# Two NLR immune receptors acquired high-affinity binding to a fungal effector through convergent evolution of their integrated domain

**DOI:** 10.1101/2021.01.26.428286

**Authors:** Aleksandra Białas, Thorsten Langner, Adeline Harant, Mauricio P Contreras, Clare EM Stevenson, David M Lawson, Jan Sklenar, Ronny Kellner, Matthew J Moscou, Ryohei Terauchi, Mark J Banfield, Sophien Kamoun

## Abstract

A subset of plant NLR immune receptors carry unconventional integrated domains in addition to their canonical domain architecture. One example is rice Pik-1 that comprises an integrated heavy metal–associated (HMA) domain. Here, we reconstructed the evolutionary history of Pik-1 and its NLR partner, Pik-2, and tested hypotheses about adaptive evolution of the HMA domain. Phylogenetic analyses revealed that the HMA domain integrated into Pik-1 before Oryzinae speciation over 15 million years ago and has been under diversifying selection. Ancestral sequence reconstruction coupled with functional studies showed that two Pik-1 allelic variants independently evolved from a weakly binding ancestral state to high-affinity binding of the blast fungus effector AVR-PikD. We conclude that for most of its evolutionary history the Pik-1 HMA domain did not sense AVR-PikD, and that different Pik-1 receptors have recently evolved through distinct biochemical paths to produce similar phenotypic outcomes. These findings highlight the dynamic nature of the evolutionary mechanisms underpinning NLR adaptation to plant pathogens.

## INTRODUCTION

<u>N</u>ucleotide-binding domain leucine-rich repeat–containing (NLR) proteins constitute an ancient class of intracellular immune receptors that confer innate immunity in plants and animals (Dodds and Rathjen, 2010; Jones et al., 2016). In plants, NLRs function by sensing pathogen-derived virulence molecules, known as effectors, and subsequently activating an immune response (Jacob et al., 2013; Kourelis and van der Hoorn, 2018). The majority of functionally validated NLRs in plants display broadly conserved domain architectures, typically consisting of the NB-ARC (<u>n</u>ucleotide-<u>b</u>inding <u>a</u>daptor shared by APAF-1, certain *<u>R</u>* gene products and <u>C</u>ED-4) domain, the LRR (leucin-rich repeat) region, and either a TIR (<u>T</u>oll/interleukin 1 receptor), CC (<u>c</u>oiled-<u>c</u>oil), or CC_R_ (RPW8-type CC) domain at the N-terminus (Kourelis and Kamoun, 2020; Shao et al., 2016). However, coevolution with pathogen effectors has led to a remarkable diversification of NLR repertoires, which form one of the most diverse protein families in plants (Lee and Chae, 2020; Prigozhin and Krasileva, 2020). An emerging paradigm in plant immunity is that some NLRs acquired novel recognition specificities through fusions of noncanonical integrated domains (IDs) that mediate perception of effectors (Cesari et al., 2014a; Wu et al., 2015). Although NLR-IDs have been described across various plant families (Gao et al., 2018; Kroj et al., 2016; Sarris et al., 2016; Van de Weyer et al., 2019), little is known about their emergence and subsequent evolution. In addition, our knowledge about how NLRs adapt to rapidly evolving pathogen effectors remains sparse.

Given that many IDs exhibit homology to molecules required for immune responses, they are generally thought to have derived from effector operative targets, which then act as baits for effector recognition within NLRs (Cesari et al., 2014a; Wu et al., 2015). IDs can perceive effectors by direct binding, by serving as substrates for their enzymatic activities, or by detecting effector-induced perturbations (Bao et al., 2017; Cesari et al., 2014a; Fujisaki et al., 2017; Heidrich et al., 2013; Sarris et al., 2016; Wu et al., 2015). The RGA5 (also known as Pia-2) and Pik-1 receptors are well-characterised examples of NLR-IDs. RGA5 and Pik-1 detect three unrelated effectors from the rice blast fungus, *Magnaporthe oryzae*, AVR-Pia/AVR1-CO39 and AVR-Pik, respectively, via their integrated heavy metal–associated (HMA) domains (De la Concepcion et al., 2018; Guo et al., 2018). HMAs are commonly found in a family of <u>H</u>MA <u>p</u>lant <u>p</u>roteins (HPPs) or <u>H</u>MA isoprenylated <u>p</u>lant <u>p</u>roteins (HIPPs) known to contribute to abiotic and biotic stress responses (De Abreu-Neto et al., 2013; Fukuoka et al., 2009; J. Li et al., 2020; Radakovic et al., 2018; Zschiesche et al., 2015). Recently, the AVR-Pik effectors have been shown to bind and stabilise rice HMA proteins to co-opt their function in immunity (Maidment et al., 2020; Oikawa et al., 2020), providing direct evidence that integrated HMAs indeed mimic host targets of effectors.

NLR-triggered immunity is usually accompanied by the hypersensitive response (HR), a type of localized cell death associated with disease resistance. Notably, several NLR-IDs appear to have lost the ability to autonomously trigger a defence response (Cesari et al., 2014b; Zdrzałek et al., 2020). As a consequence, they often function in pairs, where the NLR-ID serves as a sensor for pathogen effectors and its partner acts as a helper that mediates activation of an immune response (Adachi et al., 2019; Bonardi et al., 2011; Feehan et al., 2020). There are now many examples of such NLR pairs, including RRS1/RPS4 from *Arabidopsis thaliana* (Saucet et al., 2015) as well as Pik-1/Pik-2 (Ashikawa et al., 2008), Pii-2/Pii-1 (Fujisaki et al., 2017), and RGA5/RGA4 (the *Pia* locus) (Cesari et al., 2014b; Okuyama et al., 2011a) from rice. Many NLR pairs are encoded by two adjacent genes in a head-to-head orientation (Bailey et al., 2018; Van de Weyer et al., 2019). This genetic linkage likely provides an evolutionary advantage by facilitating co-segregation, coevolution, and transcriptional coregulation of functionally linked genes (Baggs et al., 2017; Griebel et al., 2014). Genetic linkage may also reduce the genetic load caused by autoimmunity (Wu et al., 2018), which is a common phenomenon observed across NLRs (Alcázar et al., 2009; Bomblies et al., 2007; Chae et al., 2016; Deng et al., 2019; Yamamoto et al., 2010).

Rice Pik-1 and Pik-2 proteins form a CC-type NLR pair. Two *Pik* haplotypes, N- and K-type, are present in the genetic pool of wild and cultivated rice (Zhai et al., 2011). While the function of the N-type haplotypes remains obscure, K-type Pik NLRs confer resistance to the rice blast fungus. In the K-type pair, Pik-1 acts as a sensor that binds the AVR-Pik effector via the Pik-1–integrated HMA domain, whereas Pik-2 is required for activation of immune response upon effector recognition (Maqbool et al., 2015; Zdrzałek et al., 2020). This NLR pair was initially cloned from Tsuyuake rice (Ashikawa et al., 2008), and has since been shown to occur in allelic variants, which include Pikp, Pikm, Piks, Pikh, and Pik* (Costanzo and Jia, 2010; Jia et al., 2009; Wang et al., 2009; Yuan et al., 2011; Zhai et al., 2011). Remarkably, the integrated HMA domain is the most sequence-diverse region among Pik-1 variants, consistent with the view that the receptor is under selection imposed by AVR-Pik (Białas et al., 2018; Costanzo and Jia, 2010; De la Concepcion et al., 2020; Zhai et al., 2014). Conversely, AVR-Pik alleles carry only five amino acid replacements, all of which map to regions located at the HMA-binding interface, indicating the adaptive nature of those polymorphisms (Longya et al., 2019). While the most ancient of the AVR-Pik allelic variants, AVR-PikD, is recognised by a wide range of Pik-1 proteins, the most recent variants, AVR-PikC and AVR-PikF, evade recognition by all known Pik-1 variants (Kanzaki et al., 2012; Longya et al., 2019). These recognition specificities are thought to reflect the ongoing arms race between rice and the rice blast fungus (Białas et al., 2018; Kanzaki et al., 2012; Li et al., 2019) and have been linked to the effector–HMA binding affinity (De la Concepcion et al., 2020, 2018; Maqbool et al., 2015). Despite the wealth of knowledge about mechanisms governing effector recognition by the Pik-1–integrated HMA domain, we know little about its evolutionary history.

Evolutionary molecular biology can inform mechanistic understanding of protein function. After decades of parallel research, molecular evolution and mechanistic research are starting to be used in conjunction to unravel the molecular basis of protein function within an evolutionary framework (Delaux et al., 2019). One approach to investigate the biochemical drivers of adaptation is to reconstruct the evolutionary trajectories of proteins of interest (Dean and Thornton, 2007; Harms and Thornton, 2013; Thornton, 2004). Using phylogenetic techniques and algorithms for <u>a</u>ncestral <u>s</u>equence reconstruction (ASR) it is now possible to statistically infer ancestral sequences, which can then be synthesized, expressed, and experimentally studied in the context of modern sequences (Ashkenazy et al., 2012; Cohen and Pupko, 2011; Pupko et al., 2000). In the field of plant–microbe interactions, experimental analyses of resurrected ancestral effector sequences have helped unravel biochemical bases of effector specialisation and adaptive evolution following a host jump (Dong et al., 2014; Tanaka et al., 2019; Zess et al., 2019). To date, ancestral reconstruction hasn’t been used to study the evolution of NLR immune receptors.

Despite remarkable advances in the field of NLR biology, there is still a significant gap in our understanding of how these receptors have adapted to fast-evolving pathogens. In this work, we used the rice Pik-1/Pik-2 system, coupled with ancestral sequence reconstruction, to test hypotheses about adaptive evolution of NLRs and their integrated domains and to bridge the gap between mechanistic and evolutionary research. We leveraged the rich genetic diversity of the *Pik* genes in grasses and discovered that they likely derived from a single ancestral gene pair that emerged before the radiation of the major grass lineages. In addition, we show that the HMA integration predates speciation of Oryzinae dated at ∼15 million years ago (MYA) (Jacquemin et al., 2011; Stein et al., 2018). Functional characterisation of a resurrected ancestral HMA, dating back to early *Oryza* evolution, revealed that different allelic variants of Pik-1, Pikp-1 and Pikm-1, convergently evolved from the weakly binding ancestral state towards high-affinity binding and recognition of the AVR-PikD effector through different biochemical paths. We conclude that for most of its evolutionary history Pik-HMA did not sense AVR-PikD and that recognition of this effector is a recent adaptation. This work provides new insights into our understanding of the dynamic nature of NLR adaptive evolution.

## RESULTS

### Pik orthologues are widely present across distantly related grass species

To determine the diversity of the *Pik-1* and *Pik-2* genes across the Poaceae family (grasses), we performed a phylogenetic analysis of the entire repertoire of CC-NLRs from representative grass species. We used NLR-Parser (Steuernagel et al., 2015) to identify NLR sequences from publicly available protein databases of eight species (**Supp. table 1**). Following rigorous filtering steps (**described in Materials and Methods**), we compiled a list of 3,062 putative CC-NLRs (**Supp. file 1**), amended with known and experimentally validated NLR-type proteins from grasses (**Supp. table 2**). Next, we constructed a maximum likelihood (ML) phylogenetic tree based on protein sequences of the NB-ARC domain of recovered CC-NLRs and discovered that the Pik-1 and Pik-2 sequences fell into two phylogenetically unrelated, but well-supported, clades (**Supp. figure 1A**). Among Pik-1– and Pik-2– related sequences we detected representatives from different, often distantly related, grass species, including members of the Pooideae and Panicoideae subfamilies. To determine the topologies within these clades, we performed additional phylogenetic analyses using codon-based sequence alignments of Pik-1 and Pik-2 clade members. Both Pik-1 and Pik-2 phylogenetic trees, calculated using the ML method, revealed the relationships within the two clades (**Supp. figure 1B**). We propose that the identified clades consist of Pik-1 and Pik-2 orthologues from a diversity of grass species.

We noted that Pik-2 from *Oryza brachyantha* was N-terminally truncated as a result of a 46-bp deletion within its 5′-region (**Supp. figure 2A**). To determine whether the *O. brachyantha* population carries a full-length *Pik-2* gene, we genotyped 16 additional *O. brachyantha* accessions (**Supp. figure 2B**). We successfully amplified and sequenced six full-length *ObPik-2* genes, none of which carried the deletion present in the reference genome. We further amplified full-length *ObPik-1* genes from the selected accessions (**Supp. table 3**), confirming that both full-length *Pik-2* as well as *Pik-1* are present in this species.

Following these results, we expanded the search of Pik orthologues to ten additional species, focusing on members of the Oryzoideae subfamily (**Supp. table 4**). Using recurrent BLASTN searches combined with manual gene annotation and phylogenetic analyses, we identified additional Pik-related NLRs resulting in 41 and 44 Pik-1 and Pik-2 sequences, respectively (**Figure 1A**). Altogether, the additional Pik orthologues gave us a broad view of their occurrence in monocots. The majority of species within the Oryzinae subtribe contain single copies of *Pik-1* and *Pik-2* per accession, whereas members of the Pooideae and Panicoideae subfamilies frequently encode multiple Pik-1 or Pik-2 paralogues, with wheat carrying as many as nine and ten *Pik-1* and *Pik-2* genes, respectively. In addition, Pik-1 and Pik-2 from the *Oryza* genus formed two subclades, corresponding to the two haplotypes previously identified at the *Pik* locus, N-type and K-type (**Supp. figure 3**) (Zhai et al., 2011). We conclude that the N- and K-type *Pik* genes have been maintained through speciation and co-exist as haplotypes in different *Oryza* species. Altogether, we discovered that Pik-1 and Pik-2 orthologues are present across a wide range of grasses, including members of the Oryzoideae, Pooideae, and Panicoideae subfamilies.

**Figure 1.**
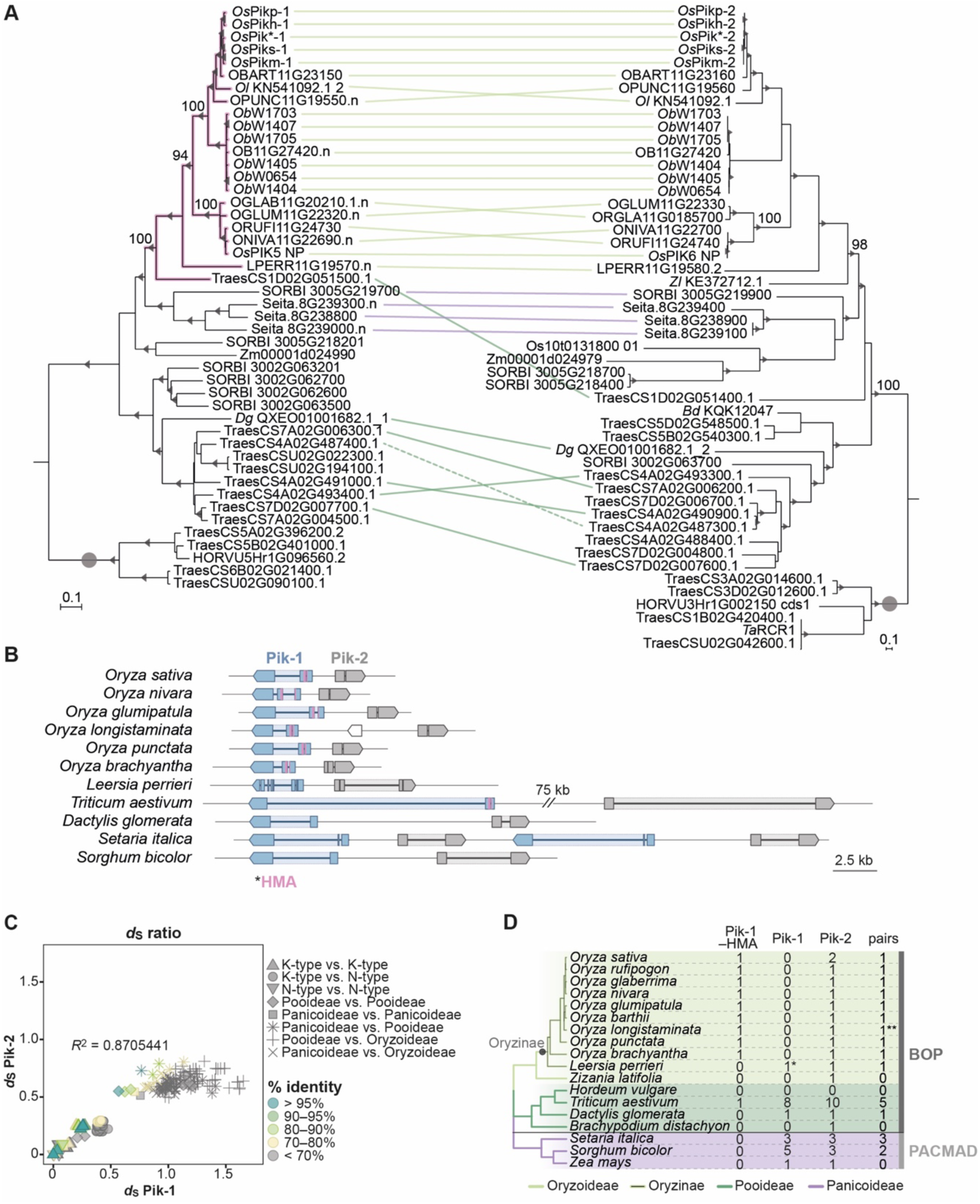
The *Pik-1*/*Pik-2* orthologues are distributed across diverse species of grasses. (**A**) The ML phylogenetic trees of Pik-1 (left) and Pik-2 (right) orthologues. The trees were calculated from 927- and 1239-nucleotide-long codon-based alignments of the NB-ARC domain, respectively, using RAxML v8.2.11 (Stamatakis, 2014), 1000 bootstrap method (Felsenstein, 1985), and GTRGAMMA substitution model (Tavaré, 1986). Best ML trees were manually rooted using the selected clades (marked with grey circle) as outgroups. The bootstrap values above 70% are indicated with grey triangles at the base of respective clades; the support values for the relevant nodes are depicted with numbers. The scale bars indicate the evolutionary distance based on nucleotide substitution rate. The Pik-1 integration clade is shown in pink. Genetically linked genes are linked with lines, with colours indicating plant subfamily: Oryzoideae (purple), Pooideae (dark green), or Panicoideae (light green); the continuous lines represent linkage in a head-to-head orientation, the dashed line indicates linkage in a tail-to-tail orientation. The interactive trees are publicly available at: https://itol.embl.de/tree/14915519290329341598279392 and https://itol.embl.de/tree/14915519290161451596745134. (**B**) Schematic illustration of the *Pik* locus in selected species. The schematic gene models of *Pik-1* (blue) and *Pik-2* (grey) are shown. The integrated HMA domain is marked with pink. The coordinates of the regions presented in this figure are summarised in **Supp. table 5**. (**C**) Comparisons of pairwise *d*_S_ rates calculated for the Pik-1 and Pik-2 receptors. The rates were calculated using Yang & Nielsen method (2000) based on 972- and 1269-nucleotide-long codon-based alignments of the NB-ARC domains of Pik-1 and Pik-2, respectively; only positions that showed over 70% coverage across the alignment were used for the analysis. The comparisons were categorised to reflect species divergence (shapes) and colour-coded to illustrate percentage identity of *d*_S_ values (% identity). The coefficient of determination (*R*^2^) was calculated for each dataset using R v3.6.3 package. (**D**) Summary of identified Pik-1 and Pik-2 homologues in plant species included in this study. The phylogenetic tree was generated using TimeTree tool (Kumar et al., 2017). The number of pairs correspond to the number of *Pik-1*/*Pik-2* genes in a head-to-head orientation separated by intergenic region of various length. (**) the species harbours a truncated gene between *Pik-1* and *Pik-2*; (*) the species has likely lost the HMA domain; Pik-1–HMA: Pik-1 with the HMA domain; Pik-1: Pik-1 without the HMA integration; BOP: <u>B</u>ambusoideae, <u>O</u>ryzoideae, <u>P</u>ooideae; PACMAD: <u>P</u>anicoideae, <u>A</u>rundinoideae, <u>C</u>hloridoideae, <u>M</u>icrairoideae, <u>A</u>ristidoideae, <u>D</u>anthonioideae.

### Genetic linkage of the *Pik* gene pair predates the split of major grass lineages

In rice, the *Pikp-1* and *Pikp-2* genes are located in a head-to-head orientation at a single locus of chromosome 11, and their coding sequences are separated by a ∼2.5-kb-long region (Ashikawa et al., 2008; Yuan et al., 2011). To determine whether this genetic linkage is conserved in grasses, we examined the genetic loci of retrieved *Pik-1* and *Pik-2* genes. A total of 14 out of 15 species in which both genes are present carry at least one *Pik* pair with adjacent *Pik-1* and *Pik-2* genes in a head-to-head orientation. Although the length of the genes and their intergenic regions vary between species (from ∼2 kb in *O. nivara* to ∼90 kb in wheat), they exhibit largely conserved gene models. Most of the *Pik-2* orthologues feature one intron in their NBD (<u>n</u>ucleotide-<u>b</u>inding <u>d</u>omain) region (Ashikawa et al., 2008) while the *Pik-1* genes typically carry one or, for the genes featuring the HMA domain, two introns (**Figure 1B**; **Supp. table 5**). In addition, in species that carry multiple copies of *Pik-1* or *Pik-2*, the copies are typically located in close proximity or, as in wheat, in large NLR-rich gene clusters (**Supp. figure 4**; **Supp. table 6**).

Given that genomic rearrangements have been reported at the *Pik* locus (Mizuno et al., 2020; Stein et al., 2018), we couldn’t exclude the possibility that genetic linkage of the *Pik-1*/*Pik-2* pair emerged more than once and is a remnant of rearrangement events. We reasoned that if the gene pair have remained genetically linked over a long evolutionary period, then they should have the same molecular age. To gain insights into the evolutionary dynamics between genetically linked Pik-1 and Pik-2 receptors, we compared their rates of synonymous substitutions (*d*_S_). For this analysis, we selected representative Pik-1 and Pik-2 NLRs that are genetically linked in a head-to-head orientation from 13 species; *Lp*Pik (*Leersia perrieri*) orthologues were excluded from the analysis because their unusual gene models interfered with sequence alignments (**Figure 1B**). Next, we assessed *d*_S_ within the coding sequences of the NB-ARC domain between pairwise genes using the Yang and Nielsen (2000) method. The rates were calculated separately for Pik-1 and Pik-2 and cross-referenced such that the pairwise values for Pik-1 were compared to the respective values for cognate Pik-2 (**Supp. table 7—source data 1**). The comparisons revealed strong positive correlation of *d*_S_ rates (*R*^2^ = 0.87) between genetically linked *Pik* genes (**Figure 1C**). This was significantly higher than observed by chance, as calculated from random Pik-1–Pik-2 cross-referencing (**Supp. figure 5**). We conclude that the Pik-1/Pik-2 pair probably became genetically linked long before the emergence of the Oryzinae clade and prior to the split of the major grass lineages—the BOP (for <u>B</u>ambusoideae, <u>O</u>ryzoideae, <u>P</u>ooideae) and PACMAD (for <u>P</u>anicoideae, <u>A</u>rundinoideae, <u>C</u>hloridoideae, <u>M</u>icrairoideae, <u>A</u>ristidoideae, <u>D</u>anthonioideae) clades—which dates back to 100–50 MYA (Hodkinson, 2018).

### The HMA integration of Pik-1 predates the emergence of Oryzinae

To better understand the evolutionary history of Pik-1 domain architecture, we looked for signatures of HMA integration among the collection of 41 Pik-1 orthologues identified. Remarkably, the presence of an HMA domain varied among *Pik-1* genes. HMA-containing Pik-1 clustered into a single well-supported clade (herein called the Pik-1 integration clade) (**Figure 1A**). All members of the Pik-1 integration clade carry the HMA domain in the same position, between the CC and NB-ARC domains of Pik-1, and feature an intron within the HMA (**Figure 1B**). This implies that these HMA domains are likely derived from a single integration event.

Using this information, we generated a sequence alignment of selected Pik-1 orthologues to define the position of the HMA integration (**Supp. figure 7**). We focused on comparisons of representative members of the Pik-1 integration clade and their closest relatives from *Setaria italica* and *Sorghum bicolor*. This revealed that the integration site most likely falls between the KLL and KTV residues (corresponding to residues 161–163 and 284– 286 of Pikp-1); however, the exact boundaries of the integration might be slightly different, given the relatively high sequence divergence around this site among the more distantly related orthologues. We further noted that the integration site encompasses a wider region than that of functionally characterised HMA domains (De la Concepcion et al., 2020, 2018), with around 20 additional amino acids (23 and 21 in Pikp-1) on each side of the annotated HMA domain.

Next, we estimated when Pik-1 acquired the HMA from the phylogeny of the plant species with Pik-1 orthologues (**Figure 1D**). We found that all *Oryza* Pik-1 orthologues carry the HMA domain, which indicates that the integration predates speciation of this genus. Although we failed to detect a full-length HMA integration in *L. perrieri*, *Lp*Pik-1 carries ∼15 amino acids characteristic of the HMA integration site (**Supp. figure 7**), indicating that the fusion probably occurred before the speciation of Oryzinae, dated at ∼15 MYA (Jacquemin et al., 2011), and was subsequently lost in *L. perrieri*. By contrast, the vast majority of examined Pik-1 from the Pooideae and Panicoideae subfamilies lack the HMA domain. The only integration in these taxonomic groups was detected in one of the nine Pik-1 paralogues of wheat included in the analysis. This observation may indicate that the Pik-1– HMA fusion may have emerged prior to radiation of the BOP clade, 100–50 MYA (Hodkinson, 2018). However, it is also possible that the integration occurred much later and that the newly emerged Pik-1–HMA gene transferred to wheat through introgression from rice progenitors. In summary, we can confidently conclude that the HMA integration of Pik-1 predates the emergence of the Oryzinae.

### The integrated HMA domain carries signatures of positive selection

In rice, the Pik-1–integrated HMA domain exhibits higher levels of polymorphisms compared with canonical domains of Pik-1 and Pik-2 (Costanzo and Jia, 2010; Kanzaki et al., 2012). To characterise the pressures underlying HMA diversification, we examined molecular signatures of selection within the Pik-1 integration clade. Wheat Pik-1–HMA was excluded from the analysis due to its high sequence divergence relative to *Oryza* orthologues, which precluded generating reliable sequence alignments. For the same reason, the remaining sequences were assigned into K- and N-type sequences based on phylogenetic relationship and analysed separately. To test for signatures of selection, we calculated rates of synonymous (*d*_S_) and nonsynonymous (*d*_N_) substitutions across the coding sequences of the HMA domain. We discovered that *d*_N_ was greater than *d*_S_ in 96 out of 115 pairwise sequence comparisons (86/105 for K- and 10/10 for N-type HMAs; *ω* = *d*_N_/*d*_S_ ranging 0–2.45 for K-type and 1.13–3.50 for N-type) (**Figure 2A, B–C—source data 2**), providing evidence that positive selection has acted on the integrated HMA domain. By contrast, only nine out of 115 pairs of the NB-ARC domain sequences of the same set of genes displayed *d*_N_ greater than *d*_S_ (**Figure 2B, C–D**); however, all of these showed *d*_S_ = 0, and were therefore inconclusive in calculating *ω* (*d*_N_/*d*_S_) ratios. A comparison of the *d*_N_ and *d*_S_ rates between the HMA and NB-ARC domains further highlighted the elevated rates of nonsynonymous substitutions within the integrated HMA domain relative to NB-ARC (**Supp. figure 8**). Overall, these results demonstrate that the integrated HMA domain exhibits marked signatures of positive selection, in contrast to the Pik-1 NB-ARC domain.

**Figure 2.**
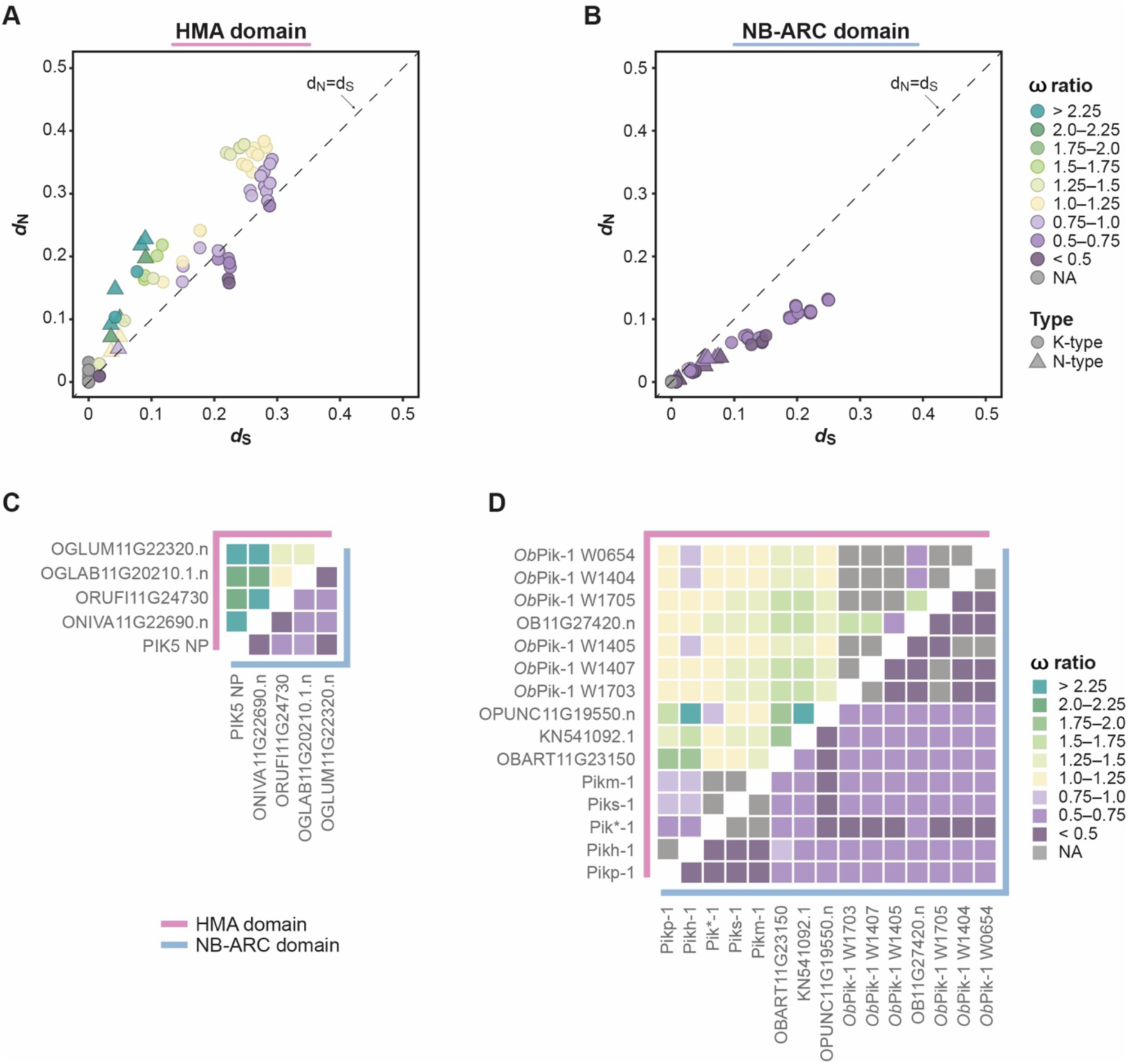
The integrated HMA domain exhibits elevated rates of ω (*d*_N_/*d*_S_) compared with the NB-ARC domain of Pik-1. (**A–B**) Pairwise comparison of nucleotide substitution rates within the Pik-1 integration clade for the (A) HMA and (**B**) NB-ARC domains, calculated using Yang & Nielsen method (2000). The diagonal line (dashed) indicates *d*_N_ = *d*_S_. The points are colour-coded to indicate ω ratio; NA: the ratio was not calculated because *d*_S_ = 0. The pairwise comparisons were separately performed for the K-type (circles) and N-type (triangles) Pik-1 sequences. (**C–D**) To highlight the differences between the ω rates for the HMA and NB-ARC domains the rates were plotted as heatmaps corresponding to the (**C**) N- and (**D**) K-type Pik-1 sequences.

Positive selection typically acts only on particular amino acids within a protein. Therefore, we aimed to detect sites within the integrated HMA domain that experienced positive selection using the ML method (Yang et al., 2000). To capture additional Pik-1–integrated HMAs, we first genotyped further wild rice species for presence of the integration. We detected the HMA integration in 21 accessions from 13 species (**Supp. table 8**); ten of those showed sufficient coverage across the entire functional region of the HMA and were used for further analysis (**Supp. figure 7; Supp. figure 9A**). We excluded the N-type HMA domains from the dataset owing to their small sample size (n = 5), which would prevent meaningful data interpretation. To detect patterns of selection within the K-type integrated HMA, we applied three pairs of ML models of codon substitution: M3/M0, M2/M1, and M8/M7 (Yang et al., 2000). As indicated by the likelihood ratio tests (LRT) and posterior probabilities, ∼26% of the HMA amino acid sites likely experienced positive selection (**Supp. figure 9B–C; Supp. file 2**). As a control, we performed the same tests on the NB-ARC domain of the K-type Pik-1 sequences. Although the discrete M3 model inferred that a subset of NB-ARC amino acids might be under diversifying selection (**Supp. figure 10**), other tests failed to detect patterns of positive selection. Based on these results, we conclude that the HMA domain exhibits strong signatures of positive selection compared with the NB-ARC domain.

### Ancestral sequence reconstruction of the Pikp-1–integrated HMA domain

To understand the evolutionary trajectory of the Pik-1–integrated HMA domain, we used representative phylogenetic trees of the K-type HMA domains to reconstruct ancestral HMA (ancHMA) sequences dating to the early stages of *Oryza* genus speciation. As an outgroup we selected HMA sequences of the integrated HMA progenitors, HPPs and HIPPs (De Abreu-Neto et al., 2013; Oikawa et al., 2020), hereafter called non-integrated HMAs, from *O. sativa* and *O. brachyantha*. To perform the reconstruction, we first tested different phylogenetic methods and focused on nodes that are well-supported in both the neighbour joining (NJ) and ML phylogenies generated from a codon-based alignment (**Supp. figure 11**). Next, we performed the ancestral sequence prediction based on protein sequence alignment, using FastML software (Ashkenazy et al., 2012), which has been previously shown to infer ancestral sequences with high accuracy (Randall et al., 2016). Multiple reconstructions yielded multiple plausible ancHMA variants (**Supp. figure 12**; **Supp. file 3**). To reduce the possibility of incorrect prediction, we selected six representative well-supported sequences for further studies.

### Reconstructed ancHMAs exhibit weaker association with AVR-PikD compared to modern Pikp-HMA

As high-affinity binding to the effector is required for the Pik-mediated immune response (De la Concepcion et al., 2020, 2019, 2018; Maqbool et al., 2015), we hypothesised that the HMA domain of Pikp-1 (Pikp-HMA) evolved towards high-affinity binding to the AVR-PikD effector. To test this hypothesis, we resurrected the six ancHMA variants determined above by synthesising their predicted sequences and incorporating them into the Pikp-1 receptor, generating Pikp-1:I-N2, Pikp-1:I-N6, Pikp-1:II-N11, Pikp-1:II-N12, Pikp-1:III-N11, and Pikp-1:III-N12 fusions (**Figure 3A**). We then tested their association with AVR-PikD in in planta co-immunoprecipitation (co-IP) experiments. The western blot analysis revealed that the ancHMA variants exhibited a range of association strengths with AVR-PikD (**Figure 3B**; **Supp. figure 13**). In every case the association with ancHMA proteins was weaker than with the present-day Pikp-HMA, indicating that binding strength has likely changed over the course of the Pikp-HMA evolutionary history. For further studies, we selected the I-N2 ancHMA variant, hereafter called ancHMA, which is the last common ancestor of Pik*-1, Pikp-1, Pikh-1, Piks-1, and Pikm-1 allelic variants of rice.

**Figure 3.**
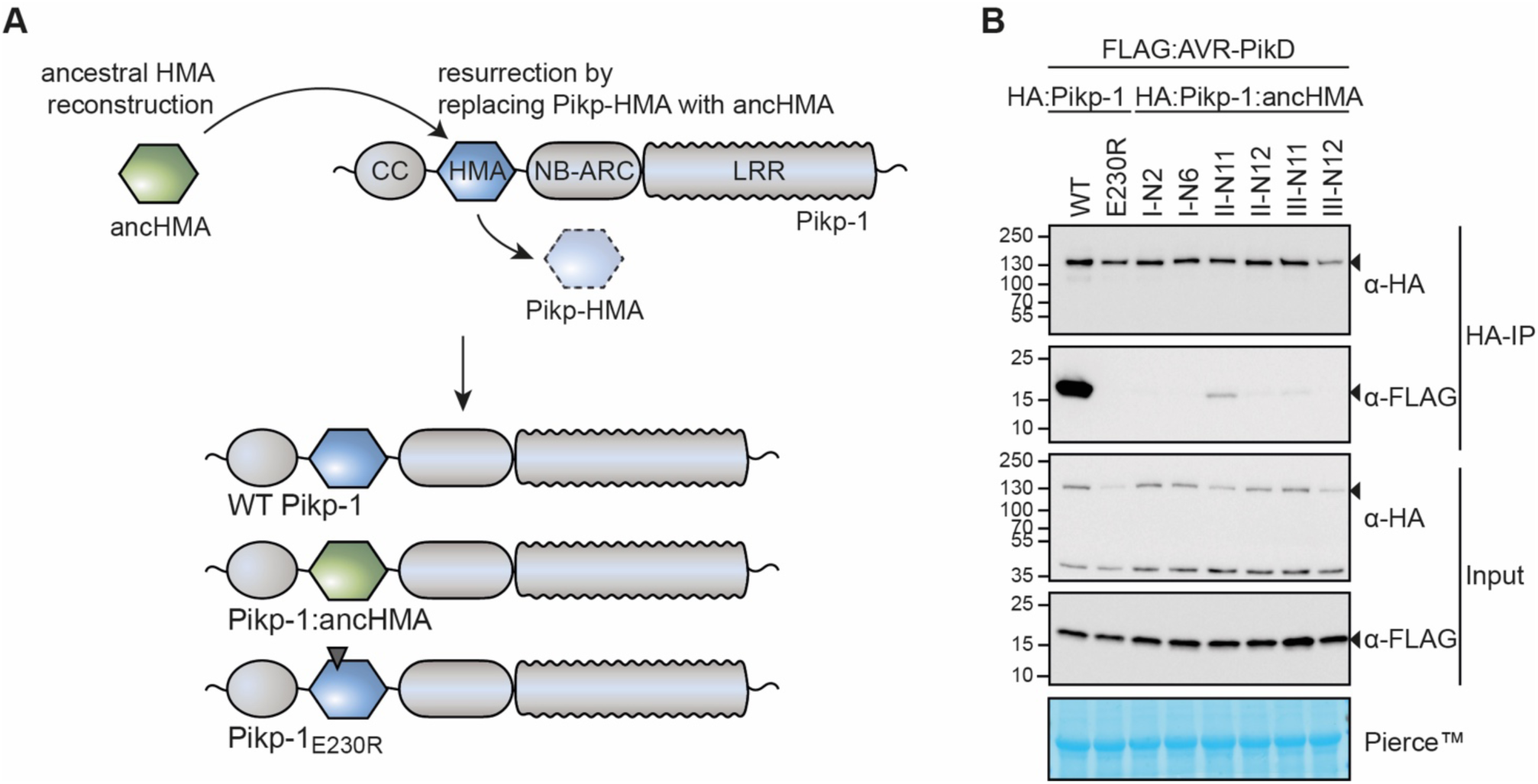
The integrated HMA domain of Pikp-1 exhibits stronger association with the AVR-PikD effector than its predicted ancestral state. (**A**) Overview of the strategy for resurrection of the ancestral HMA (ancHMA) domain. Following ancestral sequence reconstruction, the gene sequences were synthetized and incorporated into Pikp-1 by replacing the present-day Pikp-HMA domain (blue) with the ancHMA equivalent (green). (**B**) Co-IP experiment between AVR-PikD (N-terminally tagged with FLAG) and Pikp-1 (N-terminally tagged with HA) carrying ancestral sequences of the HMA. Wild-type (WT) HA:Pikp-1 and HA:Pikp-1_E230R_ were used as positive and negative controls, respectively. Immunoprecipitates (HA-IP) obtained with anti-HA probe and total protein extracts (Input) were immunoblotted with appropriate antisera (listed on the right). Rubisco loading control was performed using Pierce^TM^ staining solution. Arrowheads indicate expected band sizes. Results from three independent replicates of this experiment are shown in **Supp. figure 13**.

### The IAQVV/LVKIE region of the Pikp-HMA domain determines high-affinity AVR-PikD binding

Next, we aimed to investigate which of the structural regions in the HMA encompass adaptive mutations towards AVR-PikD binding. By combining sequence and structural information available for Pikp-HMA (De la Concepcion et al., 2018; Maqbool et al., 2015), we identified four polymorphic regions between the ancestral and modern Pikp-HMA (**Figure 4A–B**). We sequentially replaced each of these regions in Pikp-1:ancHMA with the corresponding region from Pikp-HMA. Altogether, we obtained a suite of four chimeric HMAs— ancHMA_AMEGNND_, ancHMA_LVKIE_, ancHMA_LY_, ancHMA_PI_—and assayed these for gain-of-binding to AVR-PikD in in planta co-IP experiments. Among tested constructs only the Pikp-1:ancHMA_LVKIE_ chimera associated with the effector at levels similar to Pikp-1 (**Figure 4C**; **Supp. figure 14**). This indicates that the polymorphic residues in the IAQVV/LVKIE region are critical for the evolution of enhanced AVR-PikD binding in Pikp-1.

**Figure 4.**
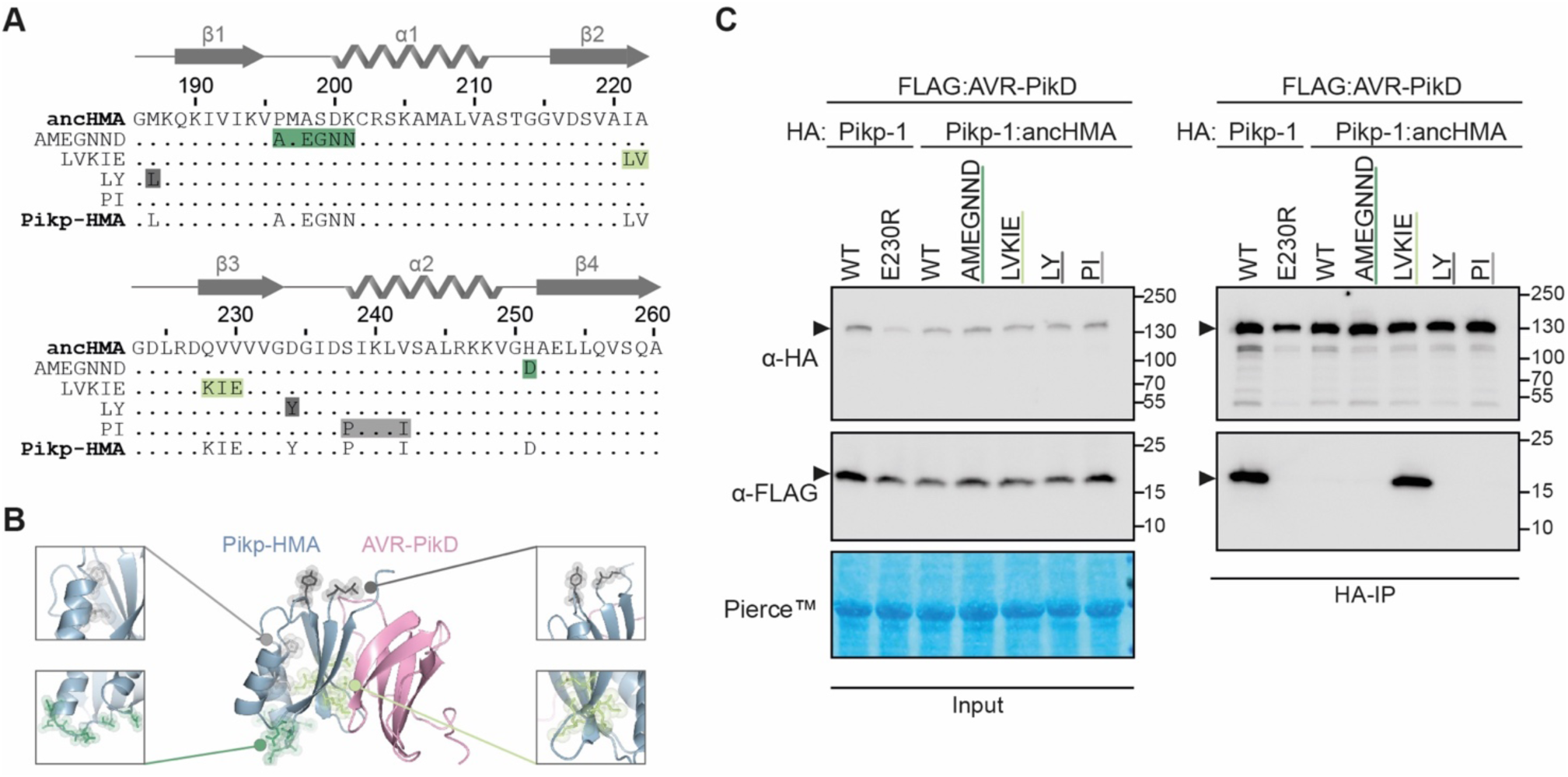
The IAQVV/LVKIE region of the Pikp-HMA domain determines high-affinity binding to AVR-PikD. (**A**) Protein sequence alignment showing the Pikp–ancHMA swap chimeras. The amino acid sequences of ancHMA, Pikp-HMA, and chimeras are aligned, with the protein model above corresponding to the Pikp-HMA structure. The colour-coded rectangles correspond to polymorphic regions used for chimeric swaps. (**B**) Schematic representation of Pikp-HMA (blue) in complex with AVR-PikD (pink) (De la Concepcion et al., 2018), with polymorphic regions between the Pikp-HMA and the ancHMA colour-coded as in the panel A. The molecular surfaces of the polymorphic residues are also shown. (**C**) Association between AVR-PikD (N-terminally tagged with FLAG) and Pikp-1, Pikp-1_E230R_, Pikp-1:ancHMA, and Pikp-1:ancHMA chimeras (N-terminally tagged with HA), labelled above, was tested in planta in co-IP experiment. Wild-type (WT) Pikp-1 and Pikp-1_E230R_ were used as a positive and negative control, respectively. Immunoprecipitates (HA-IP) obtained with anti-HA probe and total protein extracts (Input) were immunoblotted with the appropriate antisera, labelled on the left. Rubisco loading control was performed using Pierce^TM^ staining solution. Arrowheads indicate expected band sizes. Results from three independent replicates of this experiment are shown in **Supp. figure 14**.

### Two substitutions within the IAQVV/LVKIE region of ancHMA increase binding to AVR-PikD

To understand the evolutionary trajectory of the IAQVV/LVKIE region, we set out to reconstruct the evolutionary history of this region. We performed probability-based ancestral sequence reconstruction, combined with hand curation, based on protein sequence alignment and a representative phylogeny of 19 K-type integrated HMA domains, where ancHMA was separated from Pikp-HMA by five internal nodes (**Supp. figure 12**). We identified the three most ancient substitutions at the resolution of single amino acids—Ile-221-Leu, followed by Gln-228-Lys, followed by Val-229-Ile (**Figure 5A**). Discerning the order of the two most recent substitutions, Ala-222-Val and Val-230-Glu, was not possible. We generated ancHMA mutants by consecutively introducing historical substitutions into their respective ancestral backgrounds, generating ancHMA_LAQVV_, ancHMA_LAKVV_, and ancHMA_LAKIV_, as well as two plausible alternative states between LAKIV and LVKIE—ancHMA_LAKIE_ and ancHMA_LVKIV_.

**Figure 5.**
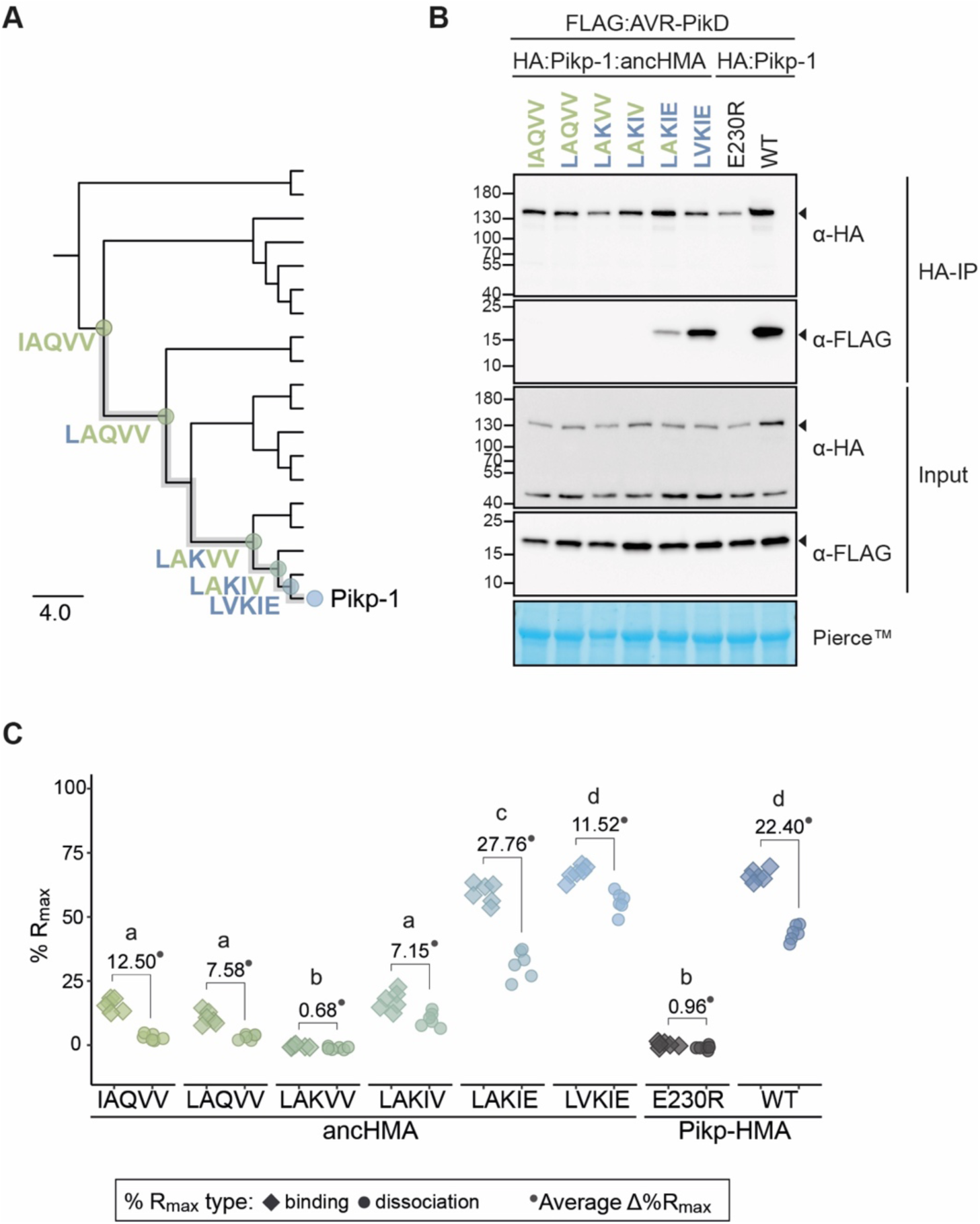
The AV-VE substitutions within the IAQVV/LVKIE region of ancHMA increase binding to AVR-PikD. (**A**) Schematic representation of an NJ phylogenetic tree of the HMA domain from *Oryza* spp. (shown in **Supp. figure 12**). The scale bar indicates the evolutionary distance based on number of base substitutions per site. Historical mutations in the IAQVV/LVKIE region acquired over the course of Pikp-HMA evolution are shown next to the appropriate nodes. The mutations are colour-coded to match the ancestral (green) and present-day (blue) states. (**B**) Co-IP experiment illustrating in planta association of AVR-PikD (N-terminally tagged with FLAG) with Pikp-1 and Pikp-1:ancHMA (N-terminally tagged with HA), labelled above. Wild-type (WT) HA:Pikp-1 and HA:Pikp-1_E230R_ proteins were used as a positive and negative control, respectively. Immunoprecipitates (HA-IP) obtained with anti-HA probe and total protein extracts (Input) were immunoblotted with appropriate antibodies (listed on the right). Loading control, featuring rubisco, was performed using Pierce^TM^ staining. The arrowheads indicate expected band sizes. Three independent replicates of this experiment are shown in **Supp. figure 16**. (**C**) Plot illustrating calculated percentage of the theoretical maximum response (%R_max_) values for interaction of HMA analytes, labelled below, with AVR-PikD ligand (featuring C-terminal HIS tag) determined using SPR. %R_max_ was normalized for the amount of ligand immobilized on the NTA-sensor chip. The chart summarises results obtained for HMA analytes at 400 nM concentration from three independent experiments with two internal repeats. Three different concentrations of the analytes (400 nM, 200 nM, 50 nM) were tested; results for the 200 nM and 50 nM concentrations are shown in **Supp. figure 18**. Average Δ%R_max_ (•) values represent absolute differences between values for ‘binding’ and ‘dissociation’, calculated from average values for each sample, and serve as an off-rate approximate. Statistical differences among the samples were analysed with Tukey’s honest significant difference (HSD) test (p < 0.01); p-values for all pairwise comparisons are presented in **Supp. table 9**.

To determine the extent to which each of the historical mutations contributed to changes in effector binding, we cloned the ancHMA mutants into the Pikp-1 background and assayed them for AVR-PikD binding in planta. Initial results showed low accumulation levels of Pikp-1:ancHMA_LVKIV_ mutant, preventing meaningful interpretation of results obtained using this protein (**Supp. figure 15**), hence, we excluded it from further analysis; the remaining constructs accumulated to similar levels. In co-IP experiments, Pikp-1:ancHMA_LVKIE_ exhibited the strongest association with AVR-PikD followed by Pikp-1:ancHMA_LAKIE_, which displayed intermediate binding (**Figure 5B**; **Supp. figure 16**). The remaining mutants did not show gain-of-binding to AVR-PikD when compared to Pikp-1:ancHMA.

To quantify how historical substitutions in the IAQVV/LVKIE region contributed to enhancing AVR-PikD binding, we carried out surface plasmon resonance (SPR) experiments, using AVR-PikD and the full set of the ancHMA mutants (cloned to match the residues Gly-186–Ser-258 of the full-length Pikp-1, which have previously been successfully used in vitro [Maqbool et al., 2015]) purified from *E. coli* by a two-step purification method (**Supp. figure 17**). We measured binding by monitoring the relative response following AVR-PikD immobilization on the NTA-sensor chip and injection of the ancHMA proteins at three different concentrations. To capture the binding dynamics, we recorded the response at two timepoints: at the end of HMA injection (‘binding’) and 15 seconds post-injection (‘dissociation’) (**Supp. figure 18A**). We normalized the response units to the theoretical maximum response (R_max_) and expressed the results as a percentage of R_max_ (%R_max_), which gave a relative indication of binding strength. Average Δ%R_max_, calculalated from a difference between R_max_ for ‘binding’ and ‘dissociation’, was used as an off-rate approximate. AncHMA_LVKIE_ formed the strongest interaction with AVR-PikD at levels similar to Pikp-HMA, followed by ancHMA_LAKIE_, then ancHMA_LAQVV_, ancHMA_LAKIV_, and ancHMA, which showed weaker interactions; we did not record any significant binding for ancHMA_LAKVV_ (**Figure 5C—source data 3**; **Supp. figure 18B—source data 3; Supp. table 9**). These results indicate that the two most recent mutations, Ala-222-Val and Val-230-Glu, collectively referred to as AV-VE, determined HMA transition towards high-affinity AVR-PikD binding.

We noted from the panel of 19 integrated HMA sequences collected in this study that the AV-VE polymorphisms are unique to Pikp-1 and Pikh-1 of rice. The *Pikp-1* and *Pikh-1* genes are highly similar to each other; out of a total of three polymorphisms, there is only one synonymous substitution that distinguishes their nearly 3,500-bp-long coding sequences (**Supp. table 10**). Although this precludes a rigorous estimation of evolutionary divergence times of the integrated HMAs, the near-absence of synonymous nucleotide polymorphisms between Pikp-1 and Pikh-1 suggests a very recent emergence of the AV-VE polymorphisms.

### The AV-VE substitutions are sufficient to increase binding affinity towards AVR-PikD

To investigate the role of historical contingency in the evolutionary history of the Pikp-1–integrated HMA domain, we tested the impact of early historical substitutions from the ancestral IAQVV residues to the Pikp-1 LVKIE on effector binding strength. We bypassed the historical sequence by incorporating the AV-VE mutations directly into ancHMA, generating Pikp:ancHMA_IVQVE_, and examined effector binding in co-IP experiments (**Supp. figure 19**). Pikp:ancHMA_IVQVE_ showed stronger association with AVR-PikD than Pikp:ancHMA; however, we were unable to directly compare its association to Pikp:ancHMA_LVKIE_ due to uneven protein accumulation levels. These results indicate that the AV-VE substitutions are sufficient to increase binding affinity towards the AVR-PikD effector independently of the other three polymorphic residues in this IAQVV/LVKIE interface. Nontheless, we cannot exclude the possibility that prior mutations had quantitative epistatic effects on the interaction that cannot be quantified by co-IP.

### High binding affinity to AVR-PikD accounts for the capacity of Pikp-1:ancHMA to trigger an immune response

To test if effector binding by Pikp-1:ancHMA is sufficient to trigger an immune response, we performed hypersensitive response (HR) cell death assays by transiently co-expressing each of the Pikp-1:ancHMA fusions with AVR-PikD and Pikp-2 in *Nicotiana benthamiana*. We discovered that all Pikp-1:ancHMA variants are autoactive and trigger spontaneous cell death in the absence of the effector (**Supp. figure 20—source data 4**; **Supp. figure 21**). Notably, the presence of the Pikp-2 partner is required for Pikp-1:ancHMA autoactivity.

Next, we used previously generated ancHMA chimeras to delimitate the region responsible for the autoactivity phenotype of Pikp-1:ancHMA. We tested these fusions for loss of function in cell death assays by transient co-expression with Pikp-2 in *N. benthamiana* (**Supp. figure 22—source data 5**; **Supp. figure 23**). Among these, Pikp-1:ancHMA_AMEGNND_ was the only chimera to show complete loss of autoactivity. This phenotype was not due to protein instability or low protein abundance (**Figure 4C**; **Supp. figure 14**). These results suggests that the PMASDKH/AMEGNND region, located in the β1–*α*1 and *α*2–β4 loops of the Pikp-HMA domain, underpins Pikp-1:ancHMA autoactivity.

To determine whether gain of AVR-PikD binding results in a functional immune response, we performed cell death assays using Pikp-1:ancHMA mutants in the IAQVV/LVKIE region. We first removed autoactivity by introducing AMEGNND mutations into these constructs (**Figure 6A**), henceforth called Pikp-1:ancHMA_LVKIE_*, Pikp-1:ancHMA_LAKIE_*, Pikp-1:ancHMA_LAKIV_*, Pikp-1:ancHMA_LAKVV_*, Pikp-1:ancHMA_LAQVV_*. None of the resulting mutants triggered spontaneous cell death when transiently co-expressed with Pikp-2 (**Figure 6B— source data 6**; **Supp. figure 24**). Co-expression with AVR-PikD revealed that the strength of binding directly correlates with the strength of HR. The mutants that gained AVR-PikD binding in the co-IP and SPR experiments, namely Pikp-1:ancHMA_LAKIE_* and Pikp-1:ancHMA_LVKIE_*, showed HR phenotypes. The Pikp-1:ancHMA_LVKIE_* mutants triggered cell death at levels similar to Pikp-1, whereas the HR triggered by Pikp-1:ancHMA_LAKIE_* was slightly, yet significantly, reduced when compared to Pikp-1. By contrast, Pikp-1:ancHMA*, Pikp-1:ancHMA_LAKVV_*, and Pikp-1:ancHMA_LAQVV_* did not elicit cell death above background levels. All proteins accumulated at similar levels in western blot analysis (**Supp. figure 25**). Overall, these results indicate that the adaptive mutations in the IAQVV/LVKIE region towards AVR-PikD binding at high affinity also enable effector-dependent activation of the cell death immune response.

**Figure 6.**
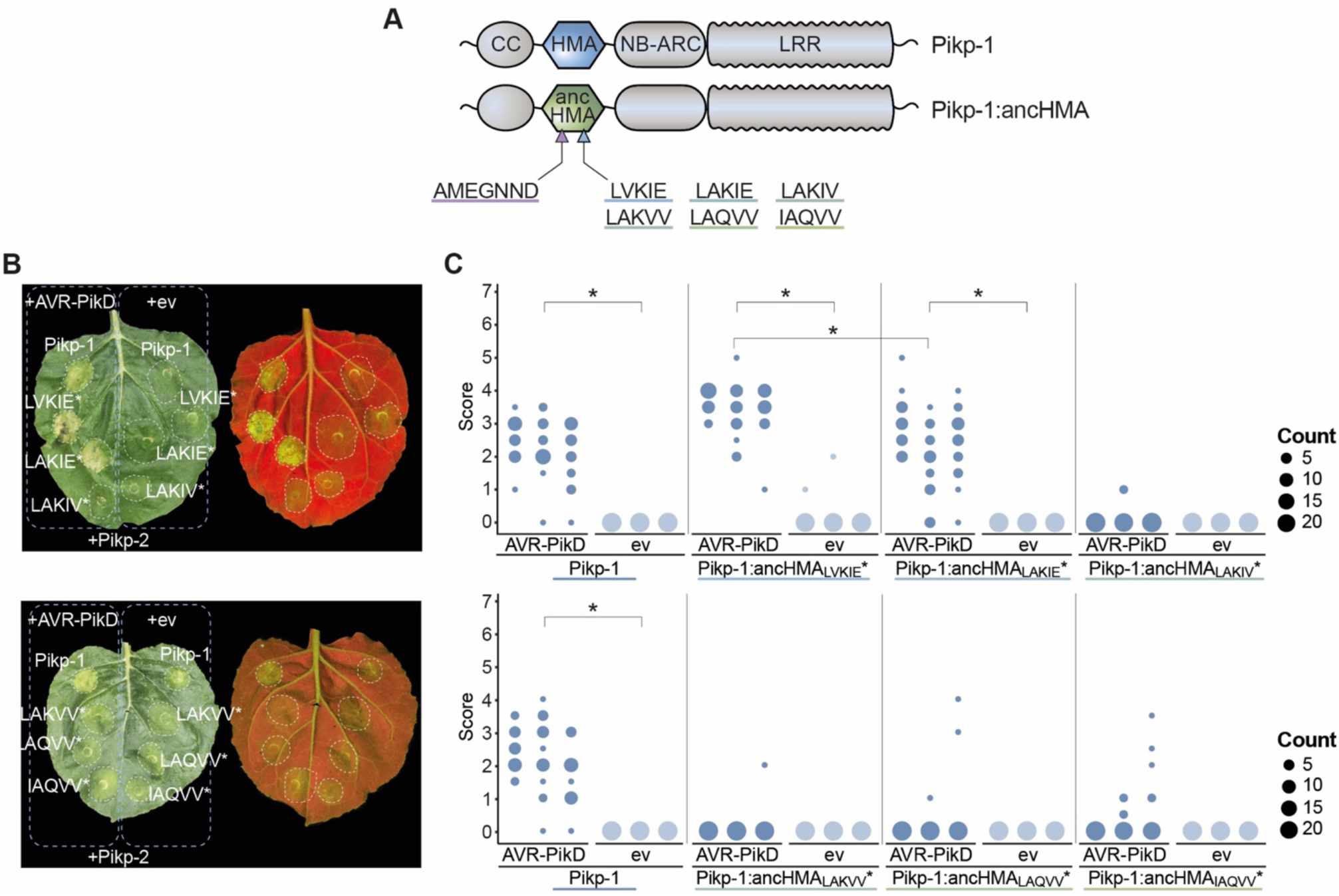
Pikp-1:ancHMA_LVKIE_* and Pikp-1:ancHMA_LAKIE_* mediate immune response towards the AVR-PikD effector. (**A**) Schematic representation of wild-type Pikp-1 and Pikp-1:ancHMA fusions used in the assay. The mutated regions are presented with arrowheads and listed. (**B**) Representative images of HR cell death assay after transient co-expression of the Pikp-1:ancHMA* mutants (C-terminally tagged with HF) with AVR-PikD (N-terminally tagged with Myc) and Pikp-2 (C-terminally tagged with HA). Empty vector (ev) was used as a negative control. All constructs were co-expressed with the gene silencing suppressor p19 (Win and Kamoun, 2003). The leaves were photographed five days after infiltration under daylight (left) and UV light (right). (**C**) HR was scored at five days post-agroinfiltration. The results are presented as dot plots, where the size of a dot is proportional to the number of samples with the same score (count) within the same biological replicate. The experiment was independently repeated at least three times with 23– 24 internal replicates; the columns within tested conditions (labelled on the bottom) correspond to results from different biological replicates. Significant differences between relevant conditions are marked with an asterisk (*); details of the statistical analysis are summarised in **Supp. figure 24**.

### A distinct region (MKANK/EMVKE) in the integrated HMA domain of Pikm-1 determines high-affinity AVR-PikD binding

As noted above, the LVKIE polymorphisms are relatively rare among Pik-1 allelic variants and *Oryza* orthologues (two out of 19 examined sequences) (**Supp. figure 26**). Other rice allelic variants of Pik-1 retain the predicted IAQVV ancestral state. Interestingly, Pikm-1, a Pik-1 allelic variant with the IAQVV residues, binds the AVR-PikD effector with high affinity and triggers an immune response upon effector recognition (De la Concepcion et al., 2018; Kanzaki et al., 2012). This led us to hypothesise that the integrated HMA domain of Pikm-1 (Pikm-HMA) has undergone a distinct evolutionary path towards AVR-PikD binding compared to Pikp-HMA.

To determine which Pikm-HMA mutations have enabled gain of AVR-PikD binding, we performed structure-informed sequence comparison of the Pikm-HMA and ancHMA domains similar to the approach described above for Pikp-1. We amended the sequence of previously predicted ancHMA with a three-amino-acid-long extension (residues 262–264 of the full-length Pikm-1) that includes residues that are polymorphic in Pikm-HMA but identical between ancHMA and Pikp-HMA. Next, we mapped five polymorphic regions that differentiate the ancHMA from modern Pikm-HMA (**Figure 7A–B**), introduced mutations in these regions in Pikm-1:ancHMA, and subjected the Pikm-1:ancHMA variants to in planta co-IP with AVR-PikD. Among the five chimeras tested in this experiment, Pikm-1:ancHMA_EMVKE_ was the only one to associate with AVR-PikD (**Figure 7C**; **Supp. figure 27**). Among the remaining chimeras, Pikm-1:ancHMA_VH_ protein was unstable and hence yielded inconclusive results. Overall, we conclude that Pikm-HMA evolved towards association with AVR-PikD through mutations in the MKANK/EMVKE region, a distinct interface from the IAQVV/LVKIE region of Pikp-1.

**Figure 7.**
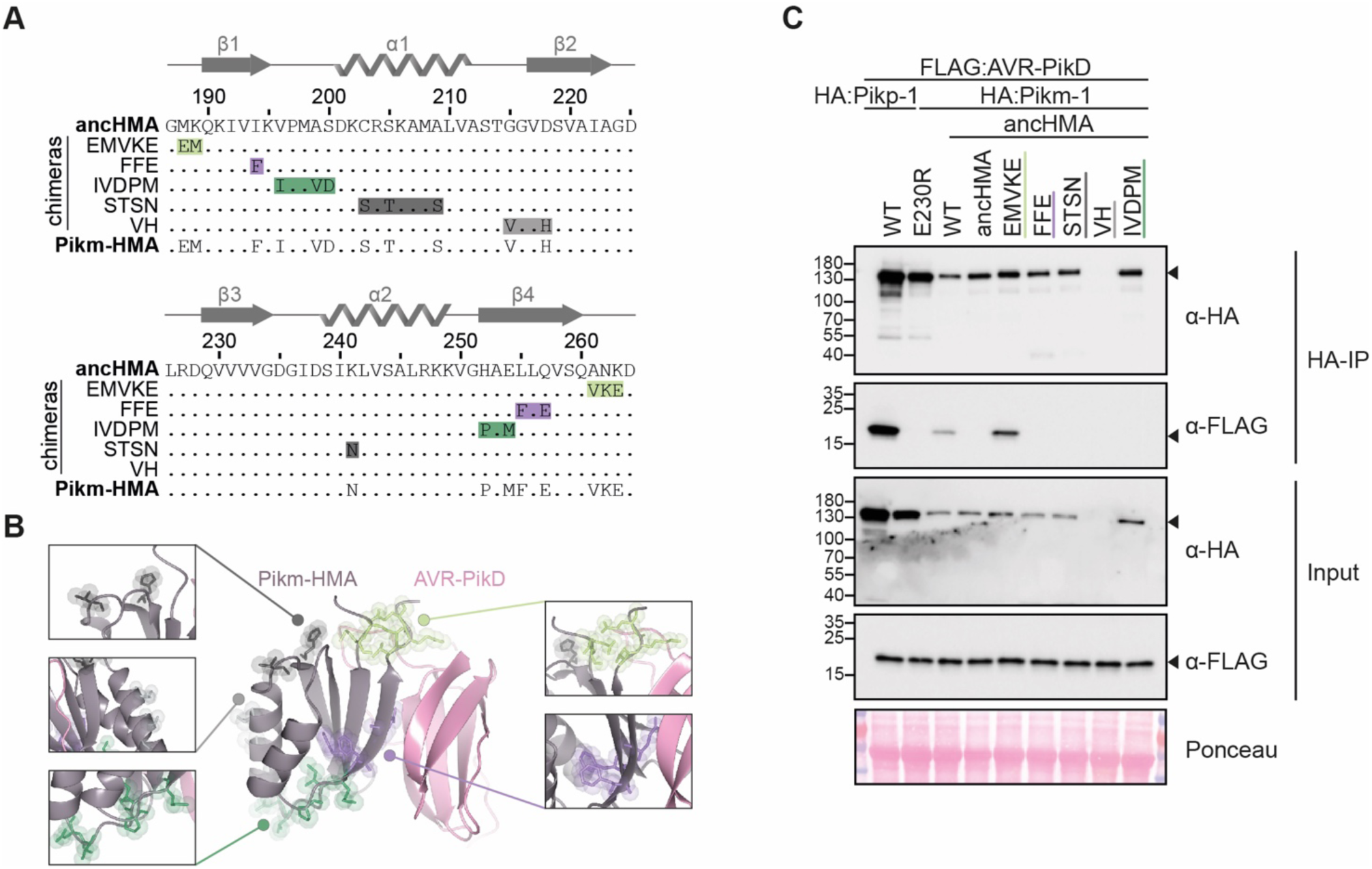
The MKANK/EMVKE region of the HMA domain of Pikm-1 determines high-affinity AVR-PikD binding. (**A**) Protein sequence alignment between the ancHMA, Pikm-HMA, and Pikm–ancHMA chimeras. The protein model above the alignment depicts Pikm-HMA seconadary structure. The colour-coded rectangles mark polymorphic regions used for chimeric swaps. (**B**) Schematic representation of the Pikm-HMA domain (purple) in complex with AVR-PikD (pink) (De la Concepcion et al., 2018), with polymorphic regions between Pikm-HMA and ancHMA colour-coded as in panel A. The molecular surfaces of the polymorphic residues are also shown. (**C**) EMVKE substitutions in the ancestral HMA restore in planta association with AVR-PikD. Co-IP experiment between AVR-PikD (N-terminally tagged with FLAG) and Pikp-1:ancHMA chimeras (N-terminally tagged with FLAG), labeled above. Wild-type (WT) Pikp-1/Pikm-1 and Pikp-1_E230R_ were used as positive and negative controls, respectively. Immunoprecipitates (HA-IP) obtained with anti-HA probe and total protein extracts (Input) were immunoblotted with the appropriate antisera (labelled on the right). Rubisco loading control was carried out using Ponceau staining. Arrowheads indicate expected band sizes. Three independent replicates of this experiment are shown in **Supp. figure 27**.

### The ANK-VKE mutations confer high-affinity AVR-PikD binding in Pikm-HMA

We reconstructed the mutational history of the MKANK/EMVKE interface to trace the evolutionary trajectory of Pikm-HMA detection of AVR-PikD (**Figure 8A**). The ancestral sequence reconstruction was performed by a combination of manual and probability-based approaches using a protein sequence alignment and a representative phylogenetic tree of the HMA domain, where ancHMA and Pikm-HMA were separated by four internal nodes (**Supp. figure 12**). However, we could only identify one node that represents an evolutionary intermediate between the ancestral MKANK and present-day EMVKE states, namely EMANK, that emerged through MK-EM mutations (Met-188-Glu and Lys-189-Met). The ANK-VKE mutations (Ala-261-Val, Asp-262-Lys, and Lys-263-Glu) were acquired at a later timepoint, and determining the order of individual mutations was not possible given the limits of the phylogenetic tree resolution.

**Figure 8.**
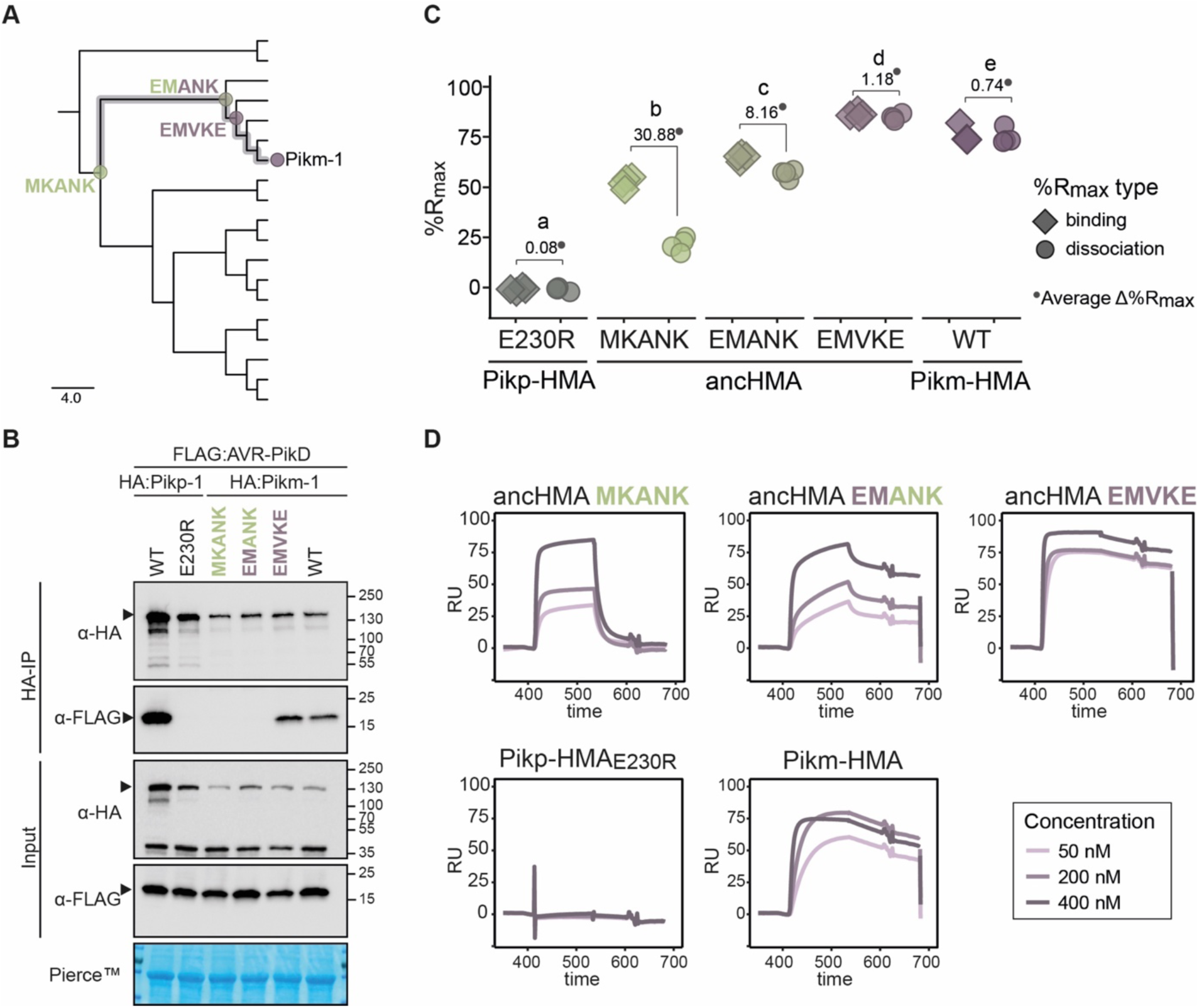
The ANK-VKE substitutions are essential for Pikm-HMA adaptation towards high-affinity binding to AVR-PikD. (**A**) Schematic representation of the NJ tree of the HMA domains from *Oryza* spp. (shown in **Supp. figure 12**). The scale bar indicates the evolutionary distance based on number of base substitutions per site. Historical substitutions in the MKANK/EMVKE region acquired over the course of Pikm-HMA evolution are shown next to the corresponding nodes. The mutations are colour-coded to match the ancestral (green) and present-day (purple) states. (A) Co-IP experiment illustrating in planta association of AVR-PikD (N-terminally tagged with FLAG) with Pikm-1 and Pikm-1:ancHMA proteins (N-terminally tagged with HA), labelled above. Wild-type (WT) Pikp-1/ Pikm-1 and Pikp-1_E230R_ constructs were used as positive and negative controls, respectively. Immunoprecipitates (HA-IP) obtained using anti-HA probes and total protein extracts (Input) were immunoblotted with the appropriate antisera (depicted on the left). The arrowheads indicate expected band sizes. Rubisco loading control was performed using Pierce^TM^ solution. Three independent replicates of this experiment are shown in **Supp. figure 31**. (**C**) Plot illustrating calculated percentage of the theoretical maximum response (%R_max_) values for interaction of HMA analytes, labelled below, with AVR-PikD ligand (C-terminally tagged with HIS) determined by SPR. %R_max_ was calculated assuming a one-to-one (HMA-to-effector) binding model for Pikm-HMA and ancHMAs, and a two-to-one for Pikp-1_E230R_. The values were normalized for the amount of ligand immobilized on the NTA-chip. The chart summarises results obtained for HMA analytes at 200 nM concentration from five independent experiments, with all the data points represented as diamonds (‘binding’) or circles (‘dissociation’). Three different concentrations of analytes (400 nM, 200 nM, 50 nM) were tested; results for 400 nM and 50 nM concentrations are shown in **Supp. figure 32**. Average Δ%R_max_ (•) values represent absolute differences between values for ‘binding’ and ‘dissociation’, calculated from average values for each sample, and serve as an off-rate approximate. Statistical differences among the samples were analysed with Tukey’s honest significant difference (HSD) test (p < 0.01); p-values for all pairwise comparisons are presented in **Supp. table 11**. (**D**) The SPR sensorgrams of the AVR-PikD and HMA proteins, corresponding to the data used in the panel C. Independent replicates of this experiment are presented in **Supp. figure 33**.

To evaluate the impact of these historical mutations, we generated the ancHMA_EMANK_ mutant that recapitulates the predicted step-by-step intermediate state of the MKANK/EMVKE region, incorporated this mutant into the Pikm-1 backbone, and assayed it for in planta association with AVR-PikD. By contrast to Pikm:ancHMA_EMVKE_, Pikm:ancHMA_EMANK_ did not gain the capacity to associate with AVR-PikD relative to Pikm:ancHMA_MKANK_ (**Figure 8B**; **Supp. figure 28**).

Next, we validated these results in vitro using the AVR-PikD protein and the full set of ancHMA mutants purified from *E. coli* (**Supp. figure 29**). To encompass the full diversity between the ancestral and present-day states of Pikm-HMA, we used HMA sequences with a five–amino acid extension at the C-terminus (ancHMA+5) compared to the constructs used in the Pikp-HMA experiments. During protein purification, we noted a shift in elution volume of the ancHMA+5 in complex with AVR-PikD relative to the elution volume of the ancHMA_LVKIE_–AVR-PikD complex in size-exclusion chromatography (**Supp. figure 30**). We concluded that this shift is consistent with different stoichiometries of the ancHMA–AVR-PikD complexes; while ancHMA_LVKIE_– AVR-PikD formed a two-to-one complex, the constructs with the extension interacted with the effector at a one-to-one ratio. Accounting for this stechiometry, we carried out SPR experiments using the same experimental design as in the Pikp-HMA assays and discovered that among tested mutants the ancHMA_EMVKE_ displayed the highest rates of interaction with AVR-PikD, followed by ancHMA_EMANK_, followed by and ancHMA_MKANK_. Although we noted that all tested HMA mutants exhibited similar binding affinity to AVR-PikD at 400 nM concentration (**Supp. figure 31—source data 7**; **Supp. table 11**), they displayed marked differences in the shapes of their sensorgrams (**Figure 8C–D—source data 7; Supp. figure 31—source data 7**; **Supp. figure 32**). First, despite high values for ‘binding’, ancHMA exhibited high off-rates, as illustrated by the pattern of ‘dissociation’ and shape of the curves. Second, ancHMA_EMVKE_ displayed high values for ‘binding’ and ‘dissociation’, with low Δ%R_max_, indicating tight and stable binding. Finally, ancHMA_EMANK_ fell in-between ancHMA and ancHMA_EMVKE_, with stable and relatively low Δ%R_max_ at the top concentration and moderate Δ%R_max_ at lower concentrations. These findings indicate that the ANK-VKE substitutions are essential for Pikm-HMA high-affinity binding of AVR-PikD. Altogether, both co-IP and SPR experiments indicate that the MKANK/EMVKE region plays an important role in high-affinity binding of the AVR-PikD effector by Pikm-HMA.

We further noted that the ANK-VKE substitutions are present in three Pik-1 alleles of rice, namely closely related Pik*-1 (Zhai et al., 2011), Pikm-1 (Ashikawa et al., 2008), and Piks-1 (Jia et al., 2009) (**Supp. figure 26**). Pikm-1 differs from Piks-1 and Pik*-1 by only two and eight amino acid polymorphisms, respectively, but no synonymous changes (**Supp. table 10**). This demonstrates a very recent emergence of these Pik-1 alleles and their associated ANK-VKE substitutions.

### Pikp-1 and Pikm-1 NLR receptors convergently evolved through distinct biochemical paths to gain high-affinity AVR-PikD binding

Our findings led us to develop an evolutionary model that depicts convergent molecular evolution of Pikp-1 and Pikm-1 towards AVR-PikD binding (**Figure 9**). To interpret this model from a structural perspective, we attempted to determine crystal structures of the ancHMA domains in complexes with AVR-PikD. Crystallisation screens of the heterologously expressed proteins resulted in crystals of the ancHMA_LVKIE_–AVR-PikD complex, which diffracted to 1.32 Å resolution (**Supp. table 12**). The structure reavealed an overall architecture of the complex similar to that of previously published co-structures of Pik-HMAs and AVR-PikD (**Supp. figure 33A**) (De la Concepcion et al., 2018; 2020; Maqbool et al., 2015). We note that the MKANK/EMVKE and IAQVV/LVKIE regions map to two of the three interaction interfaces previously described to underpin binding of AVR-PikD, and other AVR-Pik variants, to Pik-HMAs (De la Concepcion et al., 2020, 2019, 2018).

**Figure 9.**
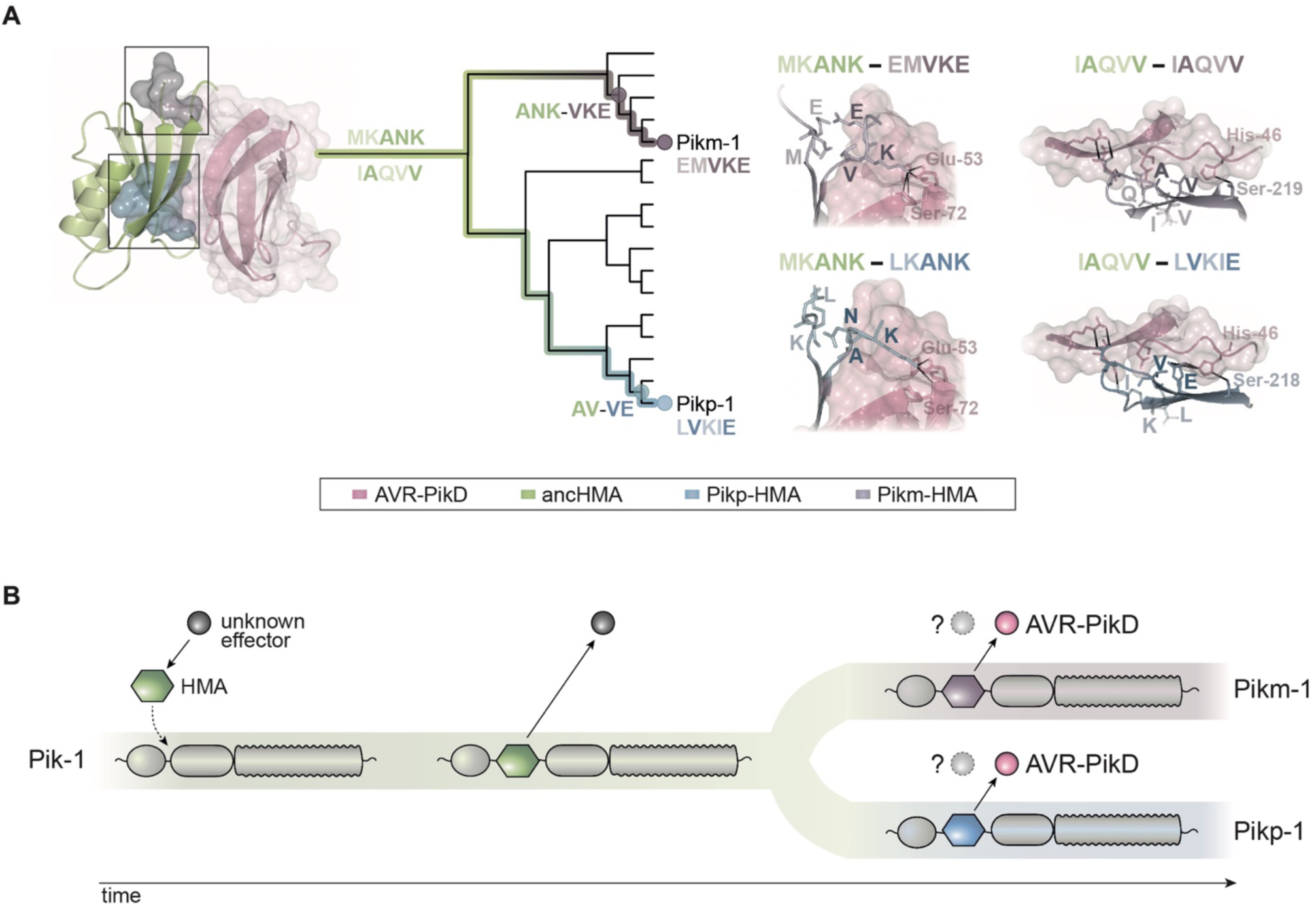
Model of molecular convergence of Pikp-1 and Pikm-1 towards AVR-PikD binding at high affinity. (**A**) The HMA domains of Pikp-1 and Pikm-1 receptors have convergently evolved through distinct evolutionary and biochemical paths to bind AVR-PikD with high affinity. The Pikp-HMA domain evolved through the AV-VE adaptations in the IAQVV/LVKIE region, whereas Pikm-HMA domain acquired the ANK-VKE mutations in the MKANK/EMVKE region. Schematic representations of the HMA–AVR-PikD structures, adapted from De la Concepcion et al. 2018, are presented with selected side chains shown as sticks and labelled; the colours of the residue labels match colours of the respective molecules. Dashed lines stand for hydrogen bonds or salt bridges. (**B**) We propose a model in which the HMA effector target integrated into Pik-1 to bait the recognition of an unknown effector. Throughout evolution the Pik-1 receptor and its integrated HMA domain diversified and led to emergence of the Pikp-1 and Pikm-1 allelic variants that bind newly emerged AVR-PikD effector.

To gain insights into the structural determinants of effector binding in the IAQVV/LVKIE region, we generated a homology model of the ancHMA in complex with AVR-PikD (**Supp. figure 33B**). We further validated modelled interactions by examining the published structure of Pikm-HMA (De la Concepcion et al., 2018), whose IAQVV/LVKIE region is identical to ancHMA. Close inspection of these structures revealed that the Val-230-Glu (V-230-E) substitution enhances the interaction with AVR-PikD through hydrogen bond formation with His-46 (**Figure 9A; Supp. figure 33C**). This bond is formed by Glu-230 (E-230) of ancHMA_LVKIE_ but absent in Pikm-HMA and ancHMA, which carry Val-230 (V-230) at the structurally equivalent position.

Next, we examined the structural basis of the interaction of the MKANK/EMVKE region with AVR-PikD by comparing Pikm- and Pikp-HMA structures (De la Concepcion et al., 2018) that feature EMVKE and LKANK residues (reminiscent of the MKANK amino acids present in ancHMA), respectively. In both cases, Lys-262 (K-262) is a major effector-binding determinant that forms hydrogen bonds or salt bridges with Glu-53 and Ser-72 of AVR-PikD (**Figure 9A**). However, in Pikm-HMA the position of Lys-262 is structurally shifted causing a difference in the conformation of the HMA peptide backbone, and associated side chains, compared to Pikp-HMA. Homology modelling fails to predict this change in the HMA backbone that results in tighter interaction between AVR-PikD and Pikm-HMA compared to Pikp-HMA (De la Concepcion et al., 2020, 2019, 2018). We conclude that Asn-261-Lys (N-261-K) and Lys-262-Glu (K-262-E) of the ANK-VKE substitution likely determine differential binding between the ancestral and present-day Pikm-HMA domains.

## DISCUSSION

The molecular evolution events associated with the transition of NLR integrated domains from pathogen effector targets to baits remain elusive. Here, we investigated the evolution of these unconventional domains of NLR receptors using rice Pik as a model system. First, we performed extensive phylogenetic analyses to determine that the integration of the HMA domain emerged over 15 MYA, predating the radiation of Oryzinae (**Figure 1D**).

Using sequence reconstruction and resurrection of an ancestral integrated HMA domain that dates back to early divergence of *Oryza* spp., we showed that the capacity of Pik-1 to sense and respond to AVR-PikD evolved relatively recently through distinct evolutionary and biochemical paths in two alleles of Pik-1, Pikp-1 and Pikm-1. This combination of evolutionary and biochemical approaches allowed us to develop a model of the adaptive evolution of the Pik proteins towards high-affinity AVR-PikD binding (**Figure 9**).

The molecular bases of functional transitions in NLR evolution remain poorly understood, especially over extended timescales. Here, we showed that adaptive evolution of Pikp-1 and Pikm-1 from weak to high-affinity binding to the AVR-PikD effector involves two distinct regions within the HMA domain. Overall, these interfaces seem to function in a synergistic yet interchangeable manner, such that weak interaction at one interface can be compensated by strong interaction at a different one (De la Concepcion et al., 2020, 2018). We propose that this modularity between different regions of the HMA increases the HMA’s capacity for rapid adaptive evolution as it can follow alternative mutational paths to produce similar phenotypic outcomes and counteract rapidly evolving pathogen effectors. Indeed, HMA domains can also detect another *M. oryzae* effector AVR-Pia through an alternative interface (Guo et al., 2018; Varden et al., 2019), further illustrating the capacity of the HMA domain to bait pathogen effectors through different interfaces. This may have contributed to the recurrent emergence of HMAs as NLR integrated domains. Previous studies have revealed that HMAs have independently integrated into NLR immune receptors from at least four flowering plant families (Kroj et al., 2016; Sarris et al., 2016).

The HMA domain of Pik-1 exhibits signatures of positive selection in contrast to the NB-ARC domain (**Figure 2**), likely reflecting coevolution with pathogen effectors versus overall purifying selection. This further suggests that HMA domains are malleable platforms that can accommodate accelerated mutational rates (Białas et al., 2018; Costanzo and Jia, 2010). Similar observations have previously been made in a number of plant NLRs, whose individual domains display patterns of asymmetrical evolution and distinct rates of selection, suggesting that NLRs evolve in a modular fashion (Kuang et al., 2004; Maekawa et al., 2019; Prigozhin and Krasileva, 2020; Read et al., 2020; Seeholzer et al., 2010). Moreover, having a domain responsible for effector recognition may release other domains from the pressure of diversification and reduce the risk of compromising or mis-regulating NLR activity (Cesari, 2018). In addition, coupling with a helper NLR such as Pik-2 likely provides yet another mechanism of functional compartmentalisation, further enhancing the evolvability of the sensor by freeing it from the constraint of executing the hypersensitive cell death (Adachi et al., 2019; Cesari, 2018; Wu et al., 2018).

We showed that the evolutionarily derived AV-VE in Pikp-1 (**Figure 5**) and ANK-VKE polymorphisms in Pikm-1 (**Figure 8**) enabled high-affinity binding to AVR-PikD. Although the high sequence divergence and elevated mutation rates among HMA sequences precluded rigorous dating of the emergence of these key adaptations, the low level of total nucleotide polymorphisms among closely related Pik alleles—in particular, the very few synonymous substitutions among Pikp- and Pikm-related alleles—points to a very recent emergence of the adaptive polymorphisms. Given that the rice-infecting lineage of *M. oryzae* is estimated to have arisen about 7,000–9,000 years ago (Couch et al., 2005; Latorre et al., 2020), our findings are consistent with the view that Pik-1 alleles evolved during rice domestication as previously suggested (Kanzaki et al., 2012; Zhai et al., 2011). In addition, AVR-Pik is widespread in rice-infecting isolates but absent in other blast fungus lineages (Langner et al., 2020; Latorre et al., 2020; Yoshida et al., 2016). Therefore, it is tempting to speculate that the rice agroecosystem has created the ecological context that led to Pik neofunctionalization towards recognition of the new pathogen threat imposed by the blast fungus. Different rice populations may have independently encountered fungal pathogens carrying AVR-Pik, leading to intense natural selection and independent emergence of the Pikp and Pikm adaptations.

We concluded that the Pik-1–integrated HMA domain did not function in sensing AVR-PikD for most of its over 15-million-year-long evolutionary history, inviting the question about the role of the ancestral integrated HMA. It is likely that over millions of years, prior to rice domestication, the Pik-1 HMA domain had recognized effectors other than AVR-Pik. These could be the structurally related MAX-effectors—an ancient effector family present across blast lineages and other fungal pathogens (de Guillen et al., 2015; Petit-Houdenot et al., 2020)—or effectors from other plant pathogen taxa. Indeed, the HMA domain is known to bind effectors from diverse pathogens including bacteria and oomycetes, in addition to fungi (González-Fuente et al., 2020). Karasov et al. (2014) proposed that NLRs caught in pairwise arms races (one NLR recognising one effector) are likely to be short-lived, whereas NLRs entangled in diffuse evolution (functioning against multiple effectors and/or multiple pathogens) are more likely to persist over longer timescales. Our model paints a more complex picture of the macroevolutionary dynamics of NLR-IDs. These receptors have the capacity to switch from one effector to another, while also engaging in short term arms race dynamics, as seems to be the case of Pik-1 vs AVR-Pik (Białas et al., 2018; Kanzaki et al., 2012). It is remarkable that the *Pik-1* gene and its paired *Pik-2* gene have been maintained in grass populations for tens of millions of years, even after the integration of the HMA domain. This points to a successful evolutionary strategy for generating long-lived disease resistance traits, with HMA promiscuity towards pathogen effectors at the centre of this model.

We discovered that the Pikp-1:ancHMA fusions trigger spontaneous hypersensitive cell death when co-expressed with Pikp-2, and mapped the region responsible for the autoactivity to two HMA parallel loops, β1–*α*1 and *α*2–β4 (**Supp. figure 20**; **Supp. figure 22**). Although the precise mechanism underpinning this autoactivity remains to be elucidated, we propose that coevolution of the HMA with the canonical domains of Pik-1 and/or Pik-2 drive this molecular incompatibility. Mismatching domains from different evolutionary timepoints may disrupt fine-tuned biochemical interactions between HMA and other domains. Indeed, intra- and intermolecular incompatibilities of NLRs are known causes of autoimmunity in plants (Harris et al., 2013; L. Li et al., 2020; Lukasik-Shreepaathy et al., 2012; Qi et al., 2012; Rairdan and Moffett, 2006; Tran et al., 2017; Wang et al., 2015). We further noted that some Pik-1 orthologues, namely *LpPik-1* and N-type *Pik-1* genes, carry large deletions within their HMAs, which may have emerged to eliminate autoimmunity (**Supp. figure 7**). This is consistent with the view that the risk of autoactivity acts as a strong evolutionary constraint narrowing NLR mutational pathways (Chae et al., 2014).

We uncovered a rich genetic diversity of *Pik* genes beyond *Oryza* species (Mizuno et al., 2020; Stein et al., 2018; Zhai et al., 2011) (**Figure 1**). This enabled us to date the emergence of the *Pik* pair to before the split of two major grass lineages: the BOP and PACMAD clades, which corresponds to 100–50 MYA (Hodkinson, 2018). Furthermore, we estimated that Pik-1 acquired the HMA domain prior the emergence of Oryzinae but after the split from Panicoideae, between 15 to 50–100 MYA (Hodkinson, 2018; Jacquemin et al., 2011; Stein et al., 2018). Remarkably, the vast majority of Pik-2 and Pik-1 orthologues across the Poaceae exist as genetically linked pairs in a head-to-head orientation. This applies to Pik-1 orthologues with and without the HMA domain, indicating that Pik-1 and Pik-2 pairing occurred prior to HMA integration. Tight genetic linkage of paired NLRs, such as Pik-1/Pik-2 (Ashikawa et al., 2008), RGA5/RGA4 (Cesari et al., 2013; Okuyama et al., 2011a), RRS1/RPS4 (Saucet et al., 2015), or RPP2A/RPP2B (Sinapidou et al., 2004), is thought to facilitate coregulation and coevolution, thereby ensuring proper cooperation between these NLRs and reducing the genetic load caused by autoimmunity (Baggs et al., 2017; Griebel et al., 2014; Wu et al., 2018). However, *Pik-1* and *Pik-2* paralogues also occur adjacent to the paired genes—a phenomenon previously observed in wild and cultivated rice (Mizuno et al., 2020)—raising the possibility that these *Pik* genes may form an NLR receptor network beyond the Pik-1/Pik-2 pair (Wu et al., 2018). In the future, it would be interesting to investigate the functions of paired Pik-1/Pik-2 and their paralogues and determine whether functional pairing and genetic linkage with *Pik-2* predisposed *Pik-1* for the HMA integration.

In summary, our study illustrates the value of ancestral sequence reconstruction—a method that has rarely been used in the field of plant–microbe interactions (Dong et al., 2014; Tanaka et al., 2019; Zess et al., 2019)—in transcending phylogenetic inference to yield a more elaborate evolutionary model. Ancestral sequence reconstruction combined with biochemical and biophysical studies enabled us to determine the directionality of evolution and therefore develop an experimentally validated model of NLR adaptation. The Pik-1/Pik-2 receptor pair emerged as an excellent system to not only provide a framework for drawing links between NLR structure and function but also to place this knowledge in an evolutionary context. This adds to our understanding of selection forces, historical contingency, and functional constraints shaping NLR activities. This approach illustrates how mechanistic research structured by a robust evolutionary framework can enhance our understanding of plant– microbe systems.

## MATERIALS AND METHODS

### Identification and phylogenetic analysis of CC-NLRs from grasses

NLR-parser (Steuernagel et al., 2015) was used to identify the NLR sequences from the predicted protein databases of eight representative grass species, *Brachypodium distachyon*, *Oryza brachyantha*, *Oryza sativa*, *Sorghum bicolor*, *Triticum aestivum*, *Zea mays* (downloaded from Ensembl Plants collection), and *Hordeum vulgare* and *Setaria italica* (downloaded from Phytozome v12.1 collection), listed in **Supp. table 1**. NLR sequences that were longer than 750 amino acid were screened for features of the NB-ARC and LRR domains, defined by the PF00931, PF00560, PF07725, PF13306, and PF13855 pfam models, using HMMER 3.2b2 (Eddy, 1998); signatures of the coiled-coil domain were identified using ‘motif16’ and ‘motif17’ defined in NLR-parser. Protein sequences of NLRs that contained at least two of the above features were aligned using MUSCLE v2.8.31 (Edgar, 2004). The proteins comprising of fewer than 60 amino acids N- and C-terminally of the NB-ARC domain, relative to the NB-ARC domain of Pikp-2 (Maqbool et al., 2015), were removed, as were sequences with less than 50% coverage across the alignment. The dataset was further filtered so that for each gene there was only one representative protein isoform—with the exception of sequences from *B. distachyon* and *S. bicolor* that didn’t carry gene identifiers. Filtering resulted in a final list of 3,062 CC-NLRs (**Supp file 1**) that were amended with 35 known and functionally characterized NLR-type resistance proteins from grasses, added for reference (**Supp. table 2**).

The amino acid sequences corresponding to the NB-ARC domain of the identified NLRs were aligned using MUSCLE v2.8.31 (Edgar, 2004). The alignment positions with more than 30% data missing were removed from the alignment using QKphylogeny (https://github.com/matthewmoscou/QKphylogeny). This revealed a final alignment of 241–amino acids, which was used for a phylogenetic analysis. A maximum likelihood (ML) phylogenetic tree was calculated using RAxML v8.2.11 (Stamatakis, 2014) with bootstrap values (Felsenstein, 1985) based on 1000 iterations and best-scoring JTT likelihood model (Jones et al., 1992) selected by automatic protein model assignment using the ML criterion. Best ML tree was mid-point rooted and visualized using Interactive Tree of Life (iToL) tool v5.5.1 (Letunic and Bork, 2007). The relationships of 28 and 38 proteins that grouped with rice Pikp-1 and Pikp-2, respectively, were further validated as follows. Genetic loci and gene coordinates for each of those NLRs were inspected and, if required, manually reannotated; identifiers of manually reannotated genes were amended with ‘.n’ suffix. For each gene, one splice version was selected and aligned using MUSCLE v2.8.31 (Edgar, 2004). The ML phylogenetic trees of Pik-1– and Pik-2–related NLRs were calculated based on positions within the NB-ARC domain, for which more than 70% of data were present—957 and 1218 nucleotides for Pik-1 and Pik-2, respectively. The trees were generated using RAxML v8.2.11 (Stamatakis, 2014) with bootstrap values (Felsenstein, 1985) based on 1000 iterations and GTRGAMMA substitution model (Tavaré, 1986). Best ML trees were manually rooted based on the relationships observed in above analyes and visualized using the iToL tool v5.5.1 (Letunic and Bork, 2007).

### Identification and phylogenetic analysis of Pik-1 and Pik-2 homologues

Coding sequences of representative Pik-1 and Pik-2 genes were used to identify Pik homologues from cDNA databases of *Oryza barthii*, *Oryza longistaminata*, *Oryza punctata*, *Oryza glumeapatula*, *Oryza glaberrima*, *Oryza rufipogon*, *Oryza nivara*, *Leersia perrieri*, *Zizania latifolia*, and *Dactylis glomerata*, listed in **Supp. table 4**, using BLAST v2.3.0 (Altschul et al., 1990). For each sequence with BLASTN E-value cutoff <0.01, genetic loci and gene coordinates were inspected and, if necessary, manually reannotated; identifiers of manually reannotated genes were amended with ‘.n’ suffix. Because the *Pik-1* and *Pik-2* genes are known to be genetically linked, each *Pik* locus was further examined for signatures of unpredicted *Pik* gene candidates. Next, coding sequences of the Pik-1 and Pik-2 candidate homologues were aligned using MUSCLE v2.8.31 (Edgar, 2004). Poorly aligned sequences were manually removed from the alignment and excluded from further analysis. The phylogenetic trees were calculated based on positions within the NB-ARC domain, for which more than 70% of data was present—927 and 1239 nucleotides of 46 Pik-1 and 54 Pik-2 candidates, respectively. ML phylogenetic trees were calculated using RAxML v8.2.11 (Stamatakis, 2014) with bootstrap values based on 1000 iterations (Felsenstein, 1985) and GTRGAMMA substitution model (Tavaré, 1986). Best ML trees were manually rooted according to previously observed relationship and visualized using the iToL tool v5.5.1 (Letunic and Bork, 2007).

### Phylogenetic analyses of rice HMA domains and ancestral sequence reconstruction

Selected non-integrated HMA sequences from *O. sativa* and *O. brachyantha* were obtained by BLASTP search (Altschul et al., 1990) using Pikp-1 HMA (Pikp-HMA) as a query. Amino acid and nucleotide alignments were generated using MUSCLE (Edgar, 2004). Neighbour joining (NJ) clustering method (Saitou and Nei, 1987) was used for constructing protein-based or codon-based trees based on JTT (Jones et al., 1992) or Maximum Composite Likelihood substitution models, respectively, using 1000 bootstrap tests (Felsenstein, 1985), as implemented in MEGA X (Kumar et al., 2018). ML trees were calculated using JTT (Jones et al., 1992) or GTR (Tavaré, 1986) substitution models as implemented in MEGA X software (Kumar et al., 2018).

Three independent protein sequence alignments, generated with MUSCLE (Edgar, 2004), were used for ancestral sequence reconstruction (**Supp. table 13**). Joint and marginal ancestral sequence reconstructions were performed with FastML software (Ashkenazy et al., 2012) using JTT substitution model (Jones et al., 1992), gamma distribution, and 90% probability cut-off to prefer ancestral indel over a character. The reconstruction was performed based on NJ trees (Saitou and Nei, 1987) built with 100 iteration bootstrap method (Felsenstein, 1985). Sequences after marginal reconstruction including indels were used for further analyses.

### Testing for selection

The rates of synonymous (*d*_S_) and nonsynonymous (*d*_N_) nucleotide substitutions per site in pairwise comparisons of protein-coding DNA sequences were estimated using the Yang and Nielsen (2000) method under realistic evolutionary models, as implemented in the YN00 program in the PAML v4.9j package (Yang, 1997). The coding sequence alignments used for the analysis were generated using MUSCLE v2.8.31 (Edgar, 2004); unless stated otherwise, only positions that showed over 70% coverage across the alignment were used for the analyses.

For selection across the sites of the HMA domain, site models were implemented using the CODEML program in the PAML v4.9j software package (Yang, 1997). The three null models, M0 (one-ratio), M1 (nearly neutral), M7 (beta), and three alternative models, M3 (selection), M2 (discrete), M8 (beta & ω), were tested as recommended by Yang et al. (2000), and their likelihoods were calculated with the likelihood ratio test. The difference in log likelihood ratio between a null model and an alternative model was multiplied by two and compared with the chi-squared (χ^2^) distribution; the degrees of freedom were calculated from the difference in the numbers of parameters estimated from the model pairs. The naïve empirical Bayes (NEB) (Yang, 2000; Yang and Nielsen, 1998) or the Bayes empirical Bayes (BEB) (Yang, 2005) were used to infer the posterior probabilities for site classes and to identify amino acids under positive selection. Raw data were extracted and visualized using the *ggplot2* R v3.6.3 package (Ginestet, 2011). ML phylogenetic tree used for the analysis was built with bootstrap values (Felsenstein, 1985) from 1000 iterations using MEGA X software (Kumar et al., 2018), based on coding sequence alignment, generated with MUSCLE v2.8.31 (Edgar, 2004).

The pairwise rates of synonymous and nonsynonymous substitutions across Pik-1 allelic variants of rice were calculated using Nei and Gojobori (1986) method, as implemented using the SNAP tool (https://www.hiv.lanl.gov/).

### Identification and cloning of *Pik-1* and *Pik-2* from *Oryza brachyantha*

Genomic DNA materials of 16 *O. brachyantha* accessions were ordered from Wild Rice Collection ‘Oryzabase’ (**Supplementary table 3**) (Kurata and Yamazaki, 2006). The accessions were first screened for deletion within the *Pik-2* gene, present in a reference genome of *O. brachyantha* (Chen et al., 2013). Selected accessions were used to amplify full-length *Pik-1* and *Pik-2* genes using 5′-TGAAGCAGATCCGAGACATAGCCT-3′ and 5′-TACCCTGCTCCTGATTGCTGACT-3′ primers designed based on the *O. brachyantha* genome sequence (Chen et al., 2013). The PCRs were run on agarose gels to check amplification and product size against positive controls. Fragments of the expected size were further gel purified, cloned into Zero Blunt® TOPO® plasmid (Thermo Fisher Scientific), and sequenced.

### Identification and cloning of the Pik-1–integrated HMA domains from wild rice relatives

Genomic DNA materials of one to three accessions of 18 wild rice species—*Oryza australiensis*, *Oryza barthii*, *Oryza brachyantha*, *Oryza eichingeri*, *Oryza glumaepatula*, *Oryza grandiglumis*, *Oryza granulata*, *Oryza latifolia*, *Oryza longiglumis*, *Oryza longistaminata*, *Oryza meridionalis*, *Oryza meyeriana*, *Oryza minuta*, Ory*za officinalis*, *Oryza punctata*, *Oryza rhizomatis*, *Oryza ridleyi*, *Oryza rufipogon*—were ordered from Wild Rice Collection ‘Oryzabase’ (Kurata and Yamazaki, 2006) and used for amplification of the Pik-1–integrated HMA (**Supp. table 8**). The 5′-AGGGAGCAATGATGCTTCACGA-3′ and 3′-TTCTCTGGCAACCGTTGTTTTGC-5′, primers were designed using the alignment of the *OsPikp-1* and *OBRAC11G13570.1* sequences and used in PCR. The amplicons were run on agarose gels to check amplification and product sizes against positive controls. Fragments of 450–720 bp in size were gel-purified, cloned into Zero Blunt® TOPO® plasmid (Thermo Fisher Scientific), and sequenced. Genotyping was performed twice and only sequences that did not show ambiguity between sequencing runs were selected for further analyses.

### Cloning for in planta assays

The rice Pikp-1, previously cloned by Maqbool et al. (2015), was amplified from pCambia1300:AscI plasmid and domesticated to remove internal *BsaI* and *BpiI* restriction enzyme recognition sites using site-directed mutagenesis by inverse PCR. The amplicons were purified and assembled using the Golden Gate method (Weber et al., 2011) in the level 0 pICH41308 (Addgene no. 47998) destination vector for subsequent Golden Gate cloning. The N-terminally tagged HA:Pikp-1 expression construct was generated by Golden Gate assembly with pICSL12008 (35S + Ω promoter, TSL SynBio), pICSL30007 (N-terminal 6×HA, TSL SynBio), and pICH41414 (35S terminator, Addgene no. 50337) modules, into the binary vector pICH47732 (Addgene no. 48001). Using the same set of Golden Gate modules, Pikp-1_E230R_ mutant was subcloned into the same binary vector, generating the N-terminally tagged HA:Pikp-1_E230R_ expression construct.

The ancestral HMA variants—corresponding to 186–260 residues of the full-length Pikp-1—were synthesised as level 0 modules for Golden Gate cloning by GENEWIZ (South Plainfield, NJ, USA). Cloning of subsequent Pikp-1:ancHMA fusions was done using two custom-made Golden Gate level 0 acceptor plasmids, p41308-PikpN and p41308-PikpC, that allowed HMA insertion in a single Golden Gate level 0 reaction, generating full-length Pikp-1 constructs with or without a stop codon, respectively. The ancestral HMA mutants— ancHMA_AMEGNND_, ancHMA_LY_, ancHMA_PI_, ancHMA_LVKIE_, and the single mutants within the LVKIE region of the ancHMA—were synthetized by GENEWIZ (South Plainfield, NJ, USA) and subcloned into p41308-PikpN and p41308-PikpC plasmids for cloning. Two of the ancHMA mutants, ancHMA_IVQVE_ and ancHMA_LVKIV_, were generated using site-directed mutagenesis by inverse PCR and cloned into the same acceptor plasmids. Using the p41308-PikpN modules, HA:Pikp-1:ancHMA expression constructs were generated by Golden Gate assembly with pICSL12008 (35S + Ω promoter, TSL SynBio), pICSL30007 (N-terminal 6×HA, TSL SynBio), and pICH41414 (35S terminator, Addgene no. 50337) into the binary vector pICH47732 (Addgene no. 48001). To generate C-terminally tagged expression constructs, the p41308-PikpC modules were assembled with pICSL13004 (Mas promoter, TSL SynBio), pICSL50001 (C-terminal HF, TSL SynBio), and pICH77901 (Mas terminator, TSL SynBio), by Golden Gate method into the same binary vector.

To generate Pikm-1:ancHMA fusions, ancHMA N2-I, ancHMA_EMVKE_, ancHMA_FFE_, ancHMA_STSN_, ancHMA_VH_, and ancHMA_IVDPM_ were synthesised by GENEWIZ (South Plainfield, NJ, USA) as Golden Gate modules. The ancHMA_EMANK_ mutant was generated by amplification and fusion of the N-terminus of ancHMA_EMVKE_ construct and the C-terminus of N2-I ancHMA variant. All ancHMA constructs corresponded to 187–264 residues of the full-length Pikm-1 protein and were subsequently assembled with custom-made p41308-PikmN (TSL SynBio) or p41308-PikmC (TSL SynBio) level 0 acceptors to generate Pikm-1:ancHMA fusions with or without a stop codon, respectively. Obtained modules were then used to generate Pikm-1:ancHMA expression constructs, featuring either N-terminal HA of C-terminal HF tags, by Golden Gate assembly using the same set of modules as previously used for Pikp-1 and pICH47732 binary vector.

### Cloning for in vitro studies

The ancHMA mutants were amplified from Golden Gate level 0 modules by PCR and cloned into and pOPIN-M vector featuring N-terminal 6×His and MBP tags with a 3C protease cleavage site, using In-Fusion cloning (Berrow et al., 2007).The AVR-PikD used for crystallography was cloned into pOPIN-S3C featuring N-terminal 6×His and SUMO tags with a 3C protease cleavage site, using In-Fusion reaction. AVR-PikD used for SPR studies was cloned previously (Maqbool et al., 2015).

### Protein–protein interaction studies: co-IP

The co-IP protocol was described previously (Win et al., 2011). Transient gene expression in planta was conducted by delivering T-DNA constructs within *Agrobacterium tumefaciens* strain GV3101::pMP90 into *Nicotiana benthamiana* leaves, and the leave tissue was collected 3 day after infiltration. Co-IP was performed using affinity chromatography with anti-HA Affinity Matrix (Roche). After co-IP and washing, the beads were resuspended in 30 μL of loading dye and eluted by incubating at 70°C for 10 minutes. Proteins were separated by SDS-PAGE and transferred onto a polyvinylidene diflouride (PVDF) membrane using a Trans-Blot turbo transfer system (Bio-Rad). The membrane was blocked with 5% non-fat dried milk powder in Tris-buffered saline and 1% Tween 20 and probed with appropriate antisera. HA-probe (F-7) horseradish peroxidase (HRP)-conjugated (Santa Cruz Biotech) was used for a single-step detection of HA tag. FLAG detection was carried using monoclonal ANTI-FLAG® M2 (Sigma) and anti-mouse HRP-conjugated antibodies in a two-step FLAG detection. A two-step detection of Myc was performed using anti-Myc (A-14, Santa Cruz Biotechnology) and anti-rabbit HRP-conjugated antibodies. Pierce ECL Western Blotting Substrate (Thermo Fisher Scientific) or SuperSignal West Femto Maximum Sensitivity Substrate (Thermo Fisher Scientific) were used for detection. Membranes were imaged using ImageQuant LAS 4000 luminescent imager (GE Healthcare Life Sciences). Equal loading was checked by staining PVDF membranes with Pierce Reversible Protein Stain Kit (Thermo Fisher Scientific), Ponceau S or Coomassie Brilliant Blue staining solutions.

### Protein–protein interaction studies: SPR

SPR experiments to investigate the effects of the IAQVV/LVKIE and MKANK/EMVKE regions were performed in the SPR buffer 1 (50 mM HEPES, pH 7.5; 300 mM NaCl; and 0.1% Tween 20) and SPR buffer 2 (50 mM HEPES, pH 7.5; 820 mM NaCl; and 0.1% Tween 20), respectively, at 25°C using Biacore T200 (GE Healthcare). The 6×His-tagged AVR-PikD (ligand) was immobilised on the Series S Sensor Chip NTA (GE Healthcare) and the HMA constructs (analytes) flowed over the effector at a flow rate of 30 μL/min. For each cycle, the chip was washed with the appropriate SPR buffer and activated with 30 μL of 0.5 mM NiCl prior immobilisation of AVR-PikD. The HMA proteins were injected over both reference and sample cells at a range of concentrations for 120 s, and buffer only flowed for 120s to record the dissociation. Between each cycle the sensor chip was regenerated with 30 μL of 0.35 M EDTA. To correct for bulk refractive index changes or machine errors, for each measurement the response was subtracted by the response in the reference cell and the response in buffer-only run (Myszka, 1999). The resulting sensorgrams were analysed using the Biacore Insight Evaluation Software (GE Healthcare).

The theoretical maximum responses (R_max_) normalized for the amount of ligand immobilized on the chip were calculated, and the level of binding was expressed as a percentage of R_max_ (%R_max_). Each experiment was repeated a minimum of three times. The data were visualised using *ggplot2* R package (Ginestet, 2011).

### Heterologous protein production and purification

Heterologous production and purification of ancHMA were performed as previously described (Varden et al., 2019). AVR-PikD and ancHMA proteins used for purification were expressed in pOPIN-S3C and pOPIN-M plasmids, respectively. AVR-PikD effector with non-cleavable C-terminal 6×His tag, used in SPR, was produced and purified as previously described (Maqbool et al., 2015). Protein intact masses were measured by static infusion of samples desalted by acetone precipitation and dissolved in 0.2% formic acid in 30% acetonitrile on Orbitrap Fusion (Thermo Scientific, UK). Data were acquired in a positive mode at 240 000 resolution and 1.6–2 kV spray voltage. The selected spectra were deisotoped and deconvoluted with Xtract software integrated in the Xcalibur package (Thermo Scientific, UK).

### Crystallisation, data collection, and structure solution

Crystallisation screens were performed at 18°C using the sitting-drop vapour diffusion technique. Drops composed of 0.3 μL of protein solution and 0.3 μL of reservoir solution were set up in MRC 96-well crystallisation plates (Molecular Dimensions), which were dispensed using an Oryx Nano or an Oryx8 robot (Douglas Instruments). Crystal growth was monitored using a Minstrel Desktop Crystal Imaging System (Rikagu). We attempted crystallisation of the ancHMA, ancHMA_LVKIE_, and ancHMA_EMVKE_ domains in complexes with AVR-PikD, but only obtained diffracting crystals for ancHMA_LVKIE_–AVR-PikD. These crystals grew after 24–48 hours in 14% (w/v) PEG 3350 and 0.2 M tri-sodium citrate, and were harvested into a cryoprotectant comprised of the precipitant augmented with 25% (v/v) ethylene glycol before flash-cooling in liquid nitrogen using LithoLoops (Molecular Dimensions). X-ray datasets were collected at the Diamond Light Source using beamline I03 (Didcot, UK) using a Pilatus3 6M hybrid photon counting detector (Dectris), with crystals maintained at 100 K by a Cryojet cryocooler (Oxford Instruments).

X-ray datasets were integrated and scaled using the DIALS xia2 pipeline (Winter, 2010) and merged with AIMLESS (Evans and Murshudov, 2013) implemented in the CCP4i2 graphical user interface (Potterton et al., 2018), with the best dataset being processed to 1.32 Å resolution in space group *P*4_1_2_1_2 with cell parameters *a* = *b* = 119.5 Å, *c* = 36.0 Å. Since the latter was isomorphous to the HMA–AVR-PikD complex previously solved (PDB accession code 5A6W, Maqbool et al., 2015), a high quality preliminary model could straightforwardly be obtained by direct refinement of the latter against the new dataset using REFMAC5 (Murshudov et al., 2011). The asymmetric unit of this preliminary model comprised one copy of AVR-PikD and two copies of ancHMA_LVKIE_. The sequences of the latter chains were subsequently corrected by manually editing the model in COOT (Emsley et al., 2010). This model was finalised by iterative rounds of manual rebuilding in COOT (Emsley et al., 2010) and restrained refinement with anisotropic thermal parameters in REFMAC5 (Murshudov et al., 2011). The resultant structure was assessed with the tools provided in COOT and MolProbity (Chen et al., 2010) and visualised using CCP4MG software (McNicholas et al., 2011).

### Homology modelling

Homology modelling of the ancHMA structure in complex with AVR-PikD was built using SWISS-MODEL (Waterhouse et al., 2018) using coordinates of Pikm-HMA–AVR-PikD structure (PDB accession 6fu9) as a template.

### Cell death assay

Expression constructs and conditions used for cell death/HR assay are listed in **Supp. table 14**. Transient expression in *N. benthamiana* leaves was conducted as previously described (Bos et al., 2006). Briefly, GV3101::pM90 *A. tumefaciens* strains carrying the appropriate expression vectors were mixed and re-suspended in infiltration buffer (10 mM 2-[N-morpholine]-ethanesulfonic acid [MES]; 10 mM MgCl_2_; and 150 μM acetosyringone, pH 5.6) to a desired density. Upper leaves of 4–5-week-old *N. benthamiana* plants were used for infiltration. The hypersensitive response (HR) cell death, was scored 5 days after agroinfiltration, using a previously published scale (Segretin et al., 2014) modified to range from 0 (no visible necrosis) to 7 (confluent necrosis).

## ACKNOWLEDGMENTS

We are thankful to several colleagues, in particular members of the Kamoun and Banfield labs, for discussions, ideas, and support. We thank Dan MacLean and other members of TSL Bioinformatics team and Mark Youles of TSL SynBio for invaluable technical support. We also thank the Diamond Light Source, UK (beamline I03 under proposal MX18565) for access to X-ray data collection facilities and Andrew Davis and Phil Robinson of JIC Bioimaging facilities for photography. This work was supported by the Gatsby Charitable Foundation, Biotechnology and Biological Sciences Research Council (BBSRC, UK), the European Research Council (ERC grant: proposal 743165) (SK, MJB), and BBSRC Doctoral Training Partnership at Norwich Research Park (grant: BB/M011216/1, project reference: 1771322) (AB).

## AUTOR CONTRIBUTIONS

Aleksandra Białas, Conceptualisation, Resources, Data curation, Formal analysis, Funding aqusition, Investigation, Methodology, Supervision, Visualisation, Writing; Thorsten Langner, Conceptualisation, Investigation, Methodology, Supervision, Writing; Adeline Harant, Investigation; Mauricio P Contreras, Investigation, Writing; Clare EM Stevenson, Data curation, Formal analysis, Methodology, Writing; David M Lawson, Data curation, Formal analysis, Methodology, Writing; Jan Sklenar, Data curation, Investigation, Writing; Ronny Kellner, Methodology; Matthew J Moscou, Resources, Methodology, Writing; Ryohei Terauchi, Resources, Funding aqusition, Methodology; Mark J. Banfield, Resources, Funding aqusition, Supervision, Writing; Sophien Kamoun, Conceptualisation, Funding aqusition, Supervision, Visualisation, Writing, Project administration

---

**Supplementary table 1.** List of databases used for NLR identification.

| **Species** | **Cultivar** | **Version** | **Assembly accession no** | **Source** |
| --- | --- | --- | --- | --- |
| ***Brachypodium distachyon*** | Bd21 | v3.0 | GCA_000005505.4 | Ensembl Plants |
| ***Hordeum vulgare*** | Morex | v2 | GCA_901482405.1 | Phytozome v12.1 |
| ***Oryza brachyantha*** | IRGC101232 | v1.4b | GCA_000231095.2 | Ensembl Plants |
| ***Oryza sativa*** | Nipponbare | IRGSP-1.0 | GCA_001433935.1 | Ensembl Plants |
| ***Setaria italica*** | Yugu1 | v2.2 | AGNK01000000.1 | Phytozome v12.1 |
| ***Sorghum bicolor*** | BTx623 | v3 | GCA_000003195.3 | Ensembl Plants |
| ***Triticum aestivum*** | Chinese Spring | v1.0 | GCA_900519105.1 | Ensembl Plants |
| ***Zea mays*** | B73 | v4 | GCA_000005005.6 | Ensembl Plants |

**Supplementary table 2.** List of known and functionally characterized NLR-type resistance proteins from grasses used as reference sequences.

| **Name** | **Accession number** | **Species** | **Reference** |
| --- | --- | --- | --- |
| **MLA10** | AY266445.1 | *Hordeum vulgare* | Halterman and Wise, 2004 |
| **RGA1-A** | KT725812.1 | *Secale cereale* | Mago et al., 2015 |
| **Os11gRGA5** | AB604627.1 | *Oryza sativa* | Okuyama et al., 2011 |
| **Os11gRGA4** | AB604622.1 | *Oryza sativa* | Okuyama et al., 2011 |
| **Piz-t** | DQ352040.1 | *Oryza sativa* | Zhou et al., 2006 |
| **Pi-ta** | AF207842.1 | *Oryza sativa* | Bryan et al., 2000 |
| **Rpg5** | EU883792.1 | *Hordeum vulgare* | Brueggeman et al., 2008 |
| **LR10** | AY270157.1 | *Triticum aestivum* | Feuillet et al., 2003 |
| **Yr10** | AF149112.1 | *Triticum aestivum* | Liu et al., 2014 |
| **Pib** | AB013448.1 | *Oryza sativa* | Wang et al., 1999 |
| **Pi9** | DQ285630.1 | *Oryza sativa* | Qu et al., 2006 |
| **Rp1-D** | XM_008664205.2 | *Zea mays* | Collins et al., 1999 |
| **Xa1** | AB002266.1 | *Oryza sativa* | Yoshimura et al., 1998 |
| **Pm8** | KF572030.1 | *Triticum aestivum* | Hurni et al., 2013 |
| **Pm3** | GU230859.1 | *Triticum aestivum* | Bhullar et al., 2010 |
| **Rdg2-a** | HM124452.1 | *Hordeum vulgare* | Bulgarelli et al., 2010 |
| **Lr21** | FJ876280.1 | *Triticum aestivum* | Huang et al., 2009 |
| **Pit** | AB379815.1 | *Oryza sativa* | Hayashi and Yoshida, 2009 |
| **Pi5-1** | EU869185.1 | *Oryza sativa* | Lee et al., 2009 |
| **Pi5-2** | EU869186.1 | *Oryza sativa* | Lee et al., 2009 |
| **Pid3** | KX791058.1 | *Oryza sativa* | Shang et al., 2009 |
| **Sr45** | LN883757.1 | *Triticum aestivum* | Steuernagel et al., 2016 |
| **Sr22** | LN883743.1 | *Triticum aestivum* | Steuernagel et al., 2016 |
| **Lr22a** | KY064064.1 | *Triticum aestivum* | Thind et al., 2017 |
| **Pik-1** | HM048900_1 | *Oryza sativa* | Zhai et al., 2011 |
| **Pik-2** | ADZ48538.1 | *Oryza sativa* | Zhai et al., 2011 |
| **Pikh-1** | HQ662330_1 | *Oryza sativa* | Costanzo and Jia, 2010b |
| **Pikh-2** | AET36550.1 | *Oryza sativa* | Costanzo and Jia, 2010b |
| **Pikm-1** | AB462324_1 | *Oryza sativa* | Ashikawa et al., 2008 |
| **Pikm-2** | BAG72135.1 | *Oryza sativa* | Ashikawa et al., 2008 |
| **Piks-1** | HQ662329_1 | *Oryza sativa* | Jia et al., 2009 |
| **Piks-2** | AET36548.1 | *Oryza sativa* | Jia et al., 2009 |
| **Pikp-1** | HM035360.1 | *Oryza sativa* | Yuan et al., 2011 |
| **Pikp-2** | ADV58351.1 | *Oryza sativa* | Yuan et al., 2011 |

**Supplementary figure 1.**
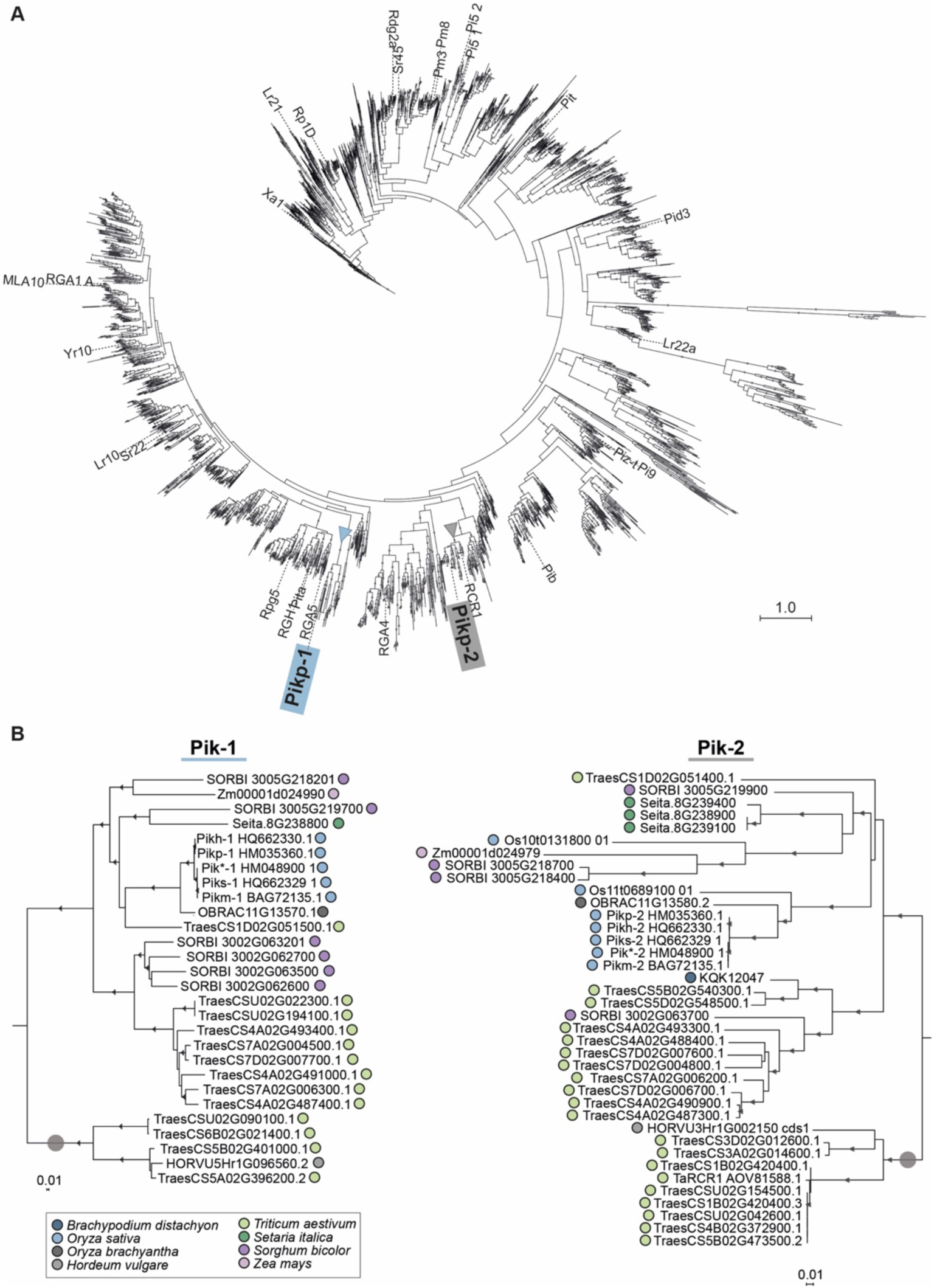
Pik-1 and Pik-2 orthologues fall into two well-supported clades. (**A**) Phylogenetic tree of CC-type NLRs of *Zea mays*, *Sorghum bicolor*, *Setaria italica*, *Triticum aestivum*, *Hordeum vulgare*, *Brachypodium distachyon*, *Oryza brachyantha*, and *Oryza sativa*. The ML tree was calculated based on 241-amino-acid-long alignment of the NB-ARC domains of 3,062 of CC-NLRs amended with 35 known and functionally characterized NLRs from grasses using RAxML v8.2.11 (Stamatakis, 2014) with bootstrap values (Felsenstein, 1985) based on 1000 iterations and the best-scoring JTT likelihood model (Jones et al., 1992). The best ML tree is shown. The scale bar indicates the evolutionary distance based on site substitution rate. The clades constituting Pik-1 and Pik-2 orthologues are marked with blue and grey triangles, respectively. Branches corresponding to the reference NLRs are labelled. The interactive tree is publicly available at: https://itol.embl.de/tree/8229133147365371602863457. (**B**) The ML phylogenetic trees of Pik-1– (left) and Pik-2–related sequences (right) constructed based on 957- and 1218-nucleotide-long codon-based alignments of the sequences of the NB-ARC domain, respectively, using RAxML v8.2.11 (Stamatakis, 2014), 1000 bootstrap method (Stamatakis, 2014), and GTRGAMMA substitution model (Tavaré, 1986). Best ML trees were manually rooted using the selected clades (marked with grey circle) as outgroups.Bootstrap values above 70% are marked with grey triangles at the base of respective clades. The scale bars indicate the evolutionary distance based on nucleotide substitution rate. The interactive trees are publicly available at: https://itol.embl.de/tree/8229133147449491602864812 and https://itol.embl.de/tree/8229133147449511602864812.

**Supplementary figure 2.**
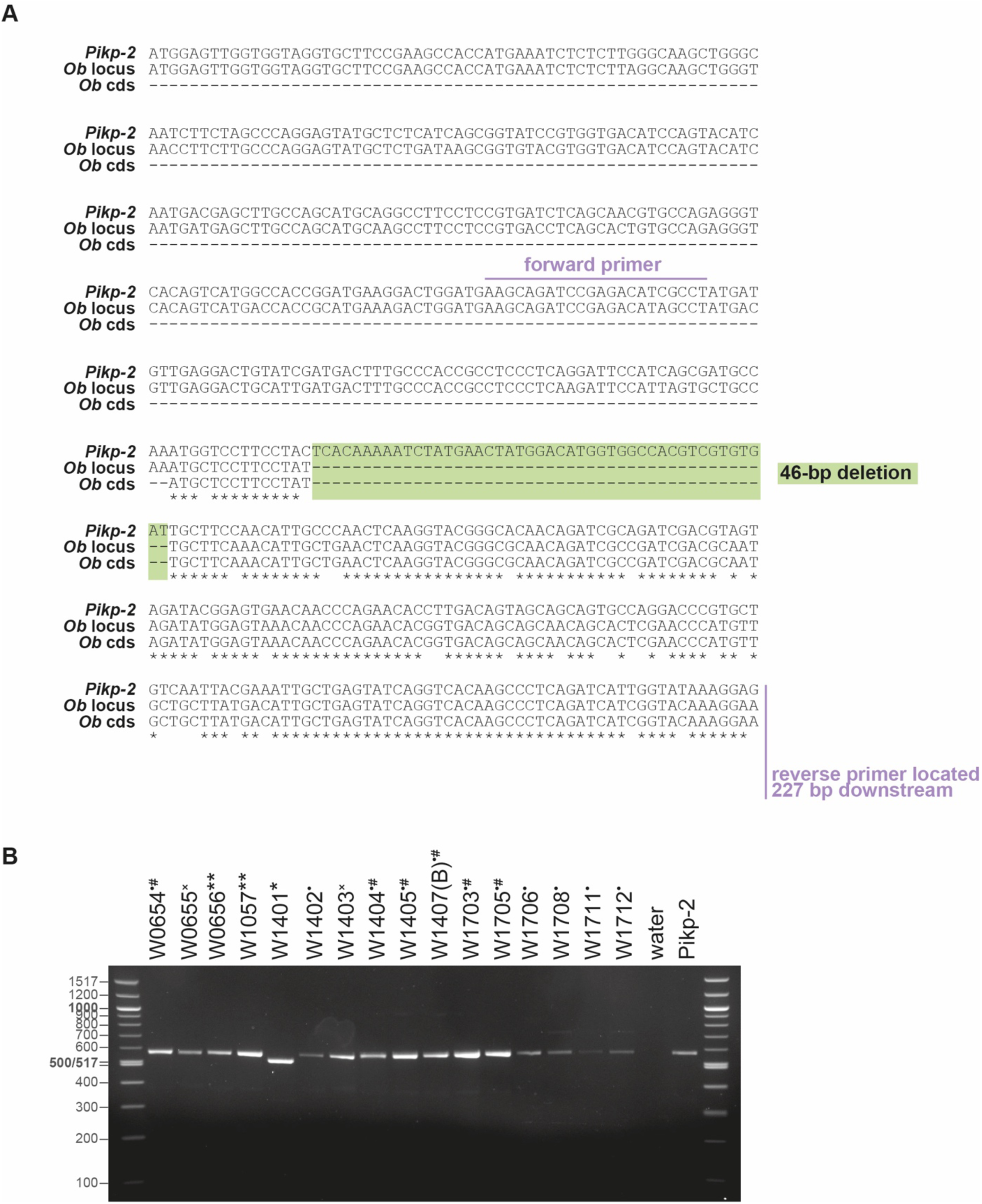
Genotyping of *Oryza brachyantha* accession. (**A**) Nucleotide alignment of *Pikp-2*, the *ObPik-2* (*Ob* locus) gene, and the *ObPik-2* coding sequence (*Ob* cds) from the reference genome (Chen et al., 2013), illustrating 46-bp-long deletion and the primers used for the genotyping. (**B**) Gel electrophoresis of *Ob*Pik-2 fragments amplified from different *O. brachyantha* accessions (labelled above). The symbols next to the accession numbers mark sequences that: carry the 46-bp deletion (*), harbour 4-bp deletion (**), carry 1-bp deletion (^×^), don’t carry any deletions (•), were used for amplification of the full-length gene (^#^). Water and *Pikp-2* were used as a negative and positive control, respectively. The left and the right lanes show molecular size markers, labelled on the left.

**Supplementary table 3.** List of *Oryza brachyantha* accessions.

| Accession | Country of origin | Used for *Pik-2* cloning | Used for *Pik-1* cloning |
| --- | --- | --- | --- |
| W0654 | Sierra Leone | Full-length | Full-length |
| W0655 | Sierra Leone | Not sequenced | Full-length |
| W0656 | Guinea | Fragment | Not amplified |
| W1057 | Guinea | Fragment | Not amplified |
| W1401 | Sierra Leone | Fragment | Not amplified |
| W1402 | Sierra Leone | Fragment | Not amplified |
| W1403 | Sierra Leone | Not sequenced | Not amplified |
| W1404 | Sierra Leone | Full-length | Full-length |
| W1405 | Sierra Leone | Full-length | Full-length |
| W1407(B) | Mali | Full-length | Full-length |
| W1703 | Mali | Full-length | Full-length |
| W1705 | Mali | Full-length | Full-length |
| W1706 | Chad | Fragment | Not amplified |
| W1708 | Cameroon | Fragment | Not amplified |
| W1711 | Cameroon | Fragment | Not amplified |
| W1712 | Cameroon | Fragment | Not amplified |

**Supplementary table 4.** List of plant datasets used for BLASTN search.

| Species | Version | Assembly accession number | Source |
| --- | --- | --- | --- |
| *Dactylis glomerata* | v1 | GCA_007115705.1 | NCBI |
| *Leersia perrieri* | v1.4 | GCA_000325765.3 | Ensembl Plants |
| *Oryza barthii* | v1 | GCA_003020155.1 | Ensembl Plants |
| *Oryza glaberrima* | v1 | GCA_000147395.2 | Ensembl Plants |
| *Oryza glumaepatula* | v1.5 | GCA_000576495.1 | Ensembl Plants |
| *Oryza longistaminata* | v1.0 | GCA_000789195.1 | Ensembl Plants |
| *Oryza nivara* | v1.0 | GCA_000576065.1 | Ensembl Plants |
| *Oryza punctata* | v1.2 | GCA_000573905.1 | Ensembl Plants |
| *Oryza rufipogon* | v1 | GCA_000817225.1 | Ensembl Plants |
| *Zizania latifolia* | v1 | GCA_000418225.1 | NCBI |

**Supplementary figure 3.**
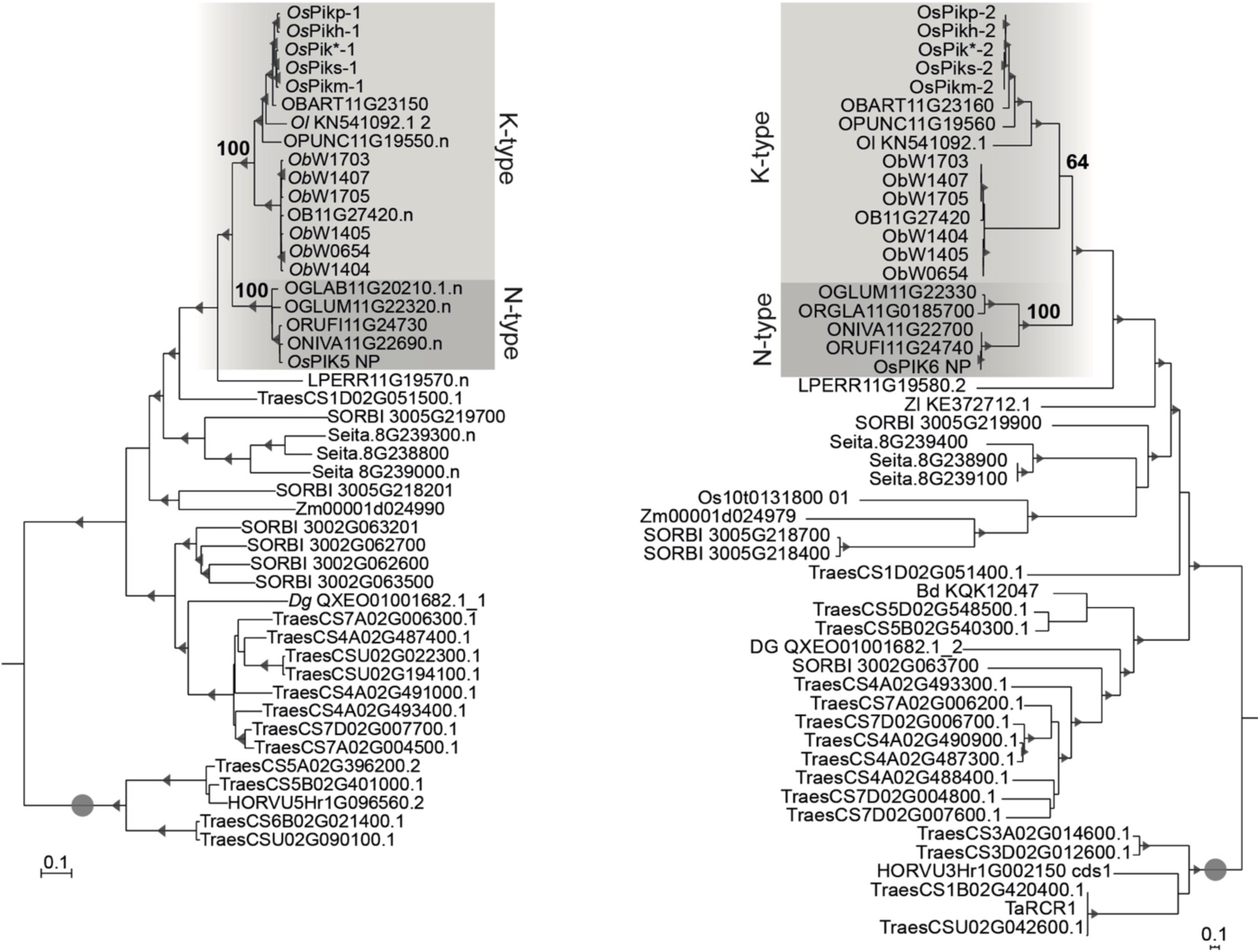
Pik-1 and Pik-2 orthologues from *Oryza* spp. fall into K- and N-type clades. The phylogenetic tree shown in Figure 3A, illustrating the divide between the N-(dark grey) and K-type (light grey) *Pik* genes. The trees were manually rooted using the selected clades (marked with grey circle) as outgroups. The bootstrap values above 70 are indicated with grey triangles at the base of respective clades; the support values for the relevant nodes are depicted with numbers. The scale bars indicate the evolutionary distance based on nucleotide substitution rate.

**Supplementary table 5.** Coordinates of genomic regions used in Figure 1B.

| **Name** | **Species** | **Contig/ Chromosome** | **Start** | **End** |
| --- | --- | --- | --- | --- |
| **Os** | *Oryza sativa* cv. K60 | Chromosome 11 | 27973977 | 28008157 |
| **Oniv** | *Oryza nivara* | Chromosome 11 | 23864010 | 23917629 |
| **Oglum** | *Oryza glumaepatula* | Chromosome 11 | 26116594 | 26116594 |
| **OI** | *Oryza longistaminata* | Contig CM003669.1 | 2965866 | 8003128 |
| **Opunc** | *Oryza punctata* | Chromosome 11 | 4972847 | 4983002 |
| **Ob** | *Oryza brachyantha* | Chromosome 11 | 15529280 | 15547390 |
| **Lp** | *Leersia perrieri* | Chromosome 11 | 20337647 | 20286182 |
| **Ta** | *Triticum aestivum* | Chromosome 1D | 33124348 | 31125148 |
| **Dg** | *Dactylis glomerata* | Scaffold QXEO01001682.1 | 1295679 | 1340805 |
| **Si** | *Setaria italica* | Scaffold 8 | 39159743 | 39261506 |
| **Sb** | *Sorghum bicolor* | Chromosome 2 | 6043453 | 6215456 |

**Supplementary figure 4.**
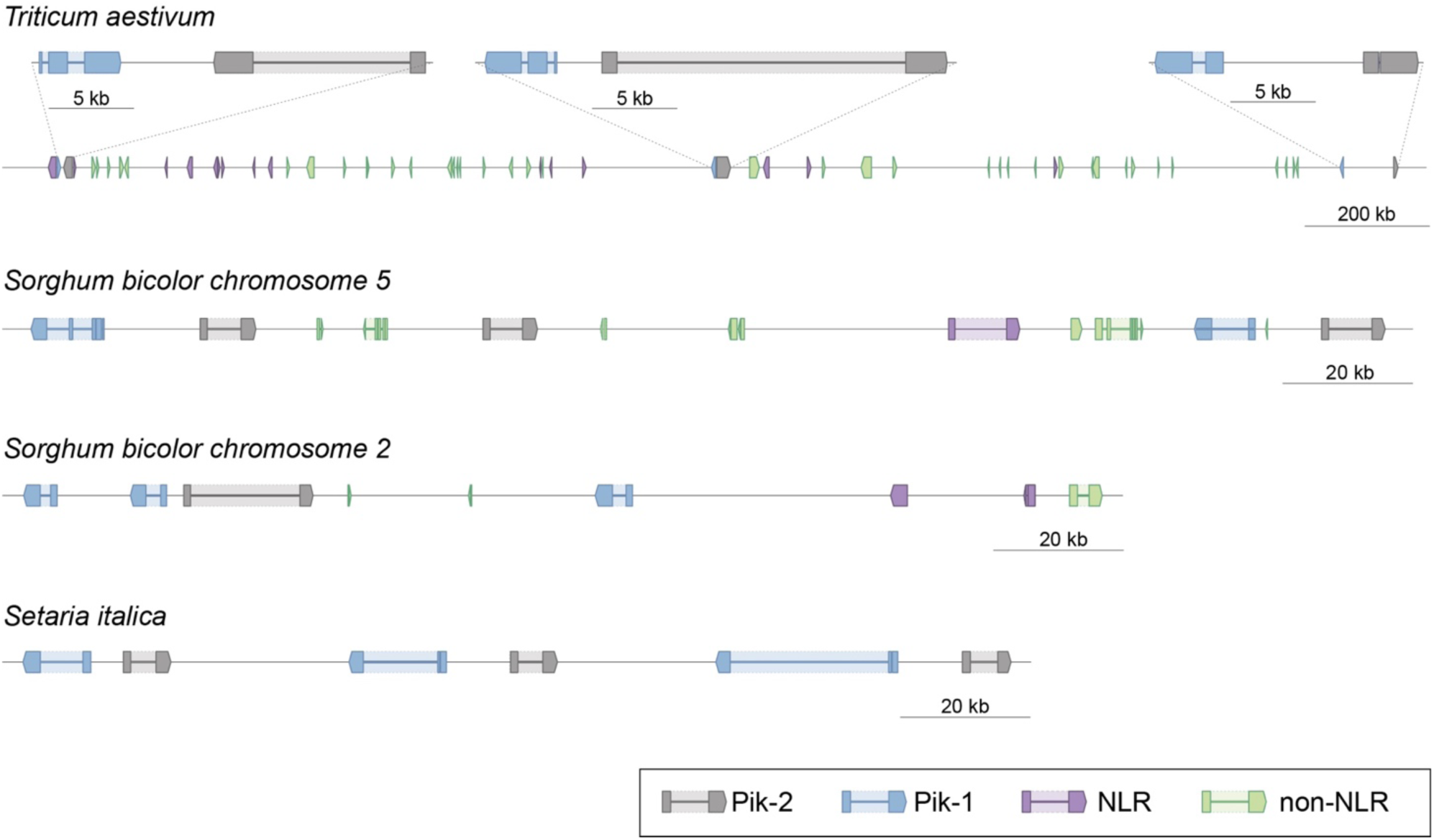
Schematic representation of selected *Pik* clusters in wheat (*T. aestivum*), sorghum (*S. bicolor*), and foxtail millet (*S. italica*). The schematic presents gene models and genetic locations of *Pik-1* (blue), *Pik-2* (grey), and other NLR genes (purple). Non-NLR genes are shown in light green. The coordinates of the regions presented in this figure are summarised in Supplementary **table 6**.

**Supplementary table 6.** Coordinates of genomic regions used in Supplementary figure 4.

| **Species** | **Chromosome / Scaffold** | **Start** | **End** |
| --- | --- | --- | --- |
| *Triticum aestivum* | Chromosome 1D | 33124348 | 31125148 |
| *Triticum aestivum* | Chromosome 4A | 739372651 | 742475497 |
| *Triticum aestivum* | Chromosome 7D | 3066514 | 4338541 |
| *Sorghum bicolor* | Chromosome 5 | 70290712 | 70702146 |
| *Sorghum bicolor* | Chromosome 2 | 5911252 | 6248983 |
| *Setaria italica* | Scaffold 8 | 39100454 | 39349775 |

**Supplementary table 7.**
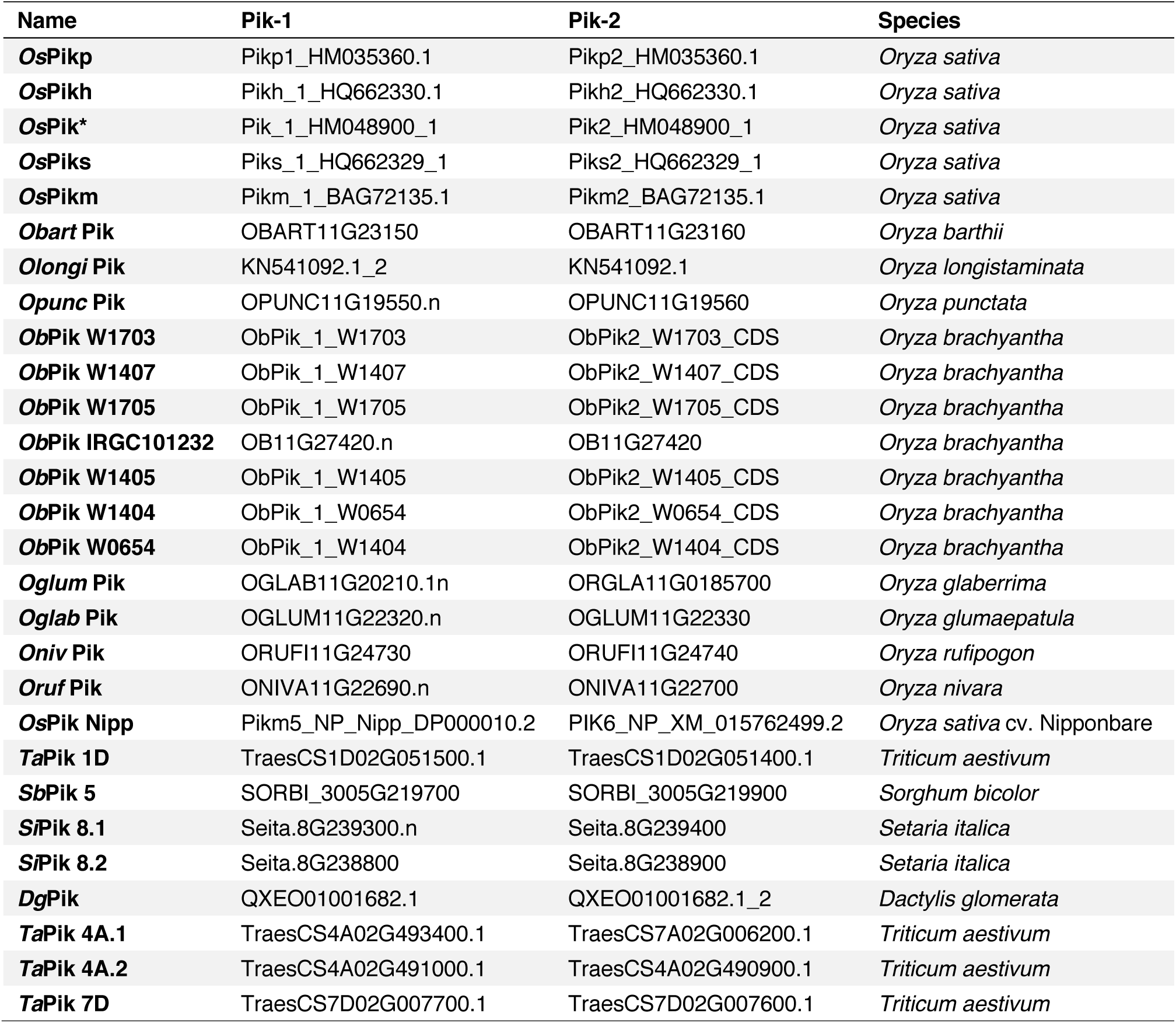
Genes used for the comparisons of *d*_S_ and rates of Pik-1–Pik-2 presented in Figure 1C.

**Supplementary figure 5.**
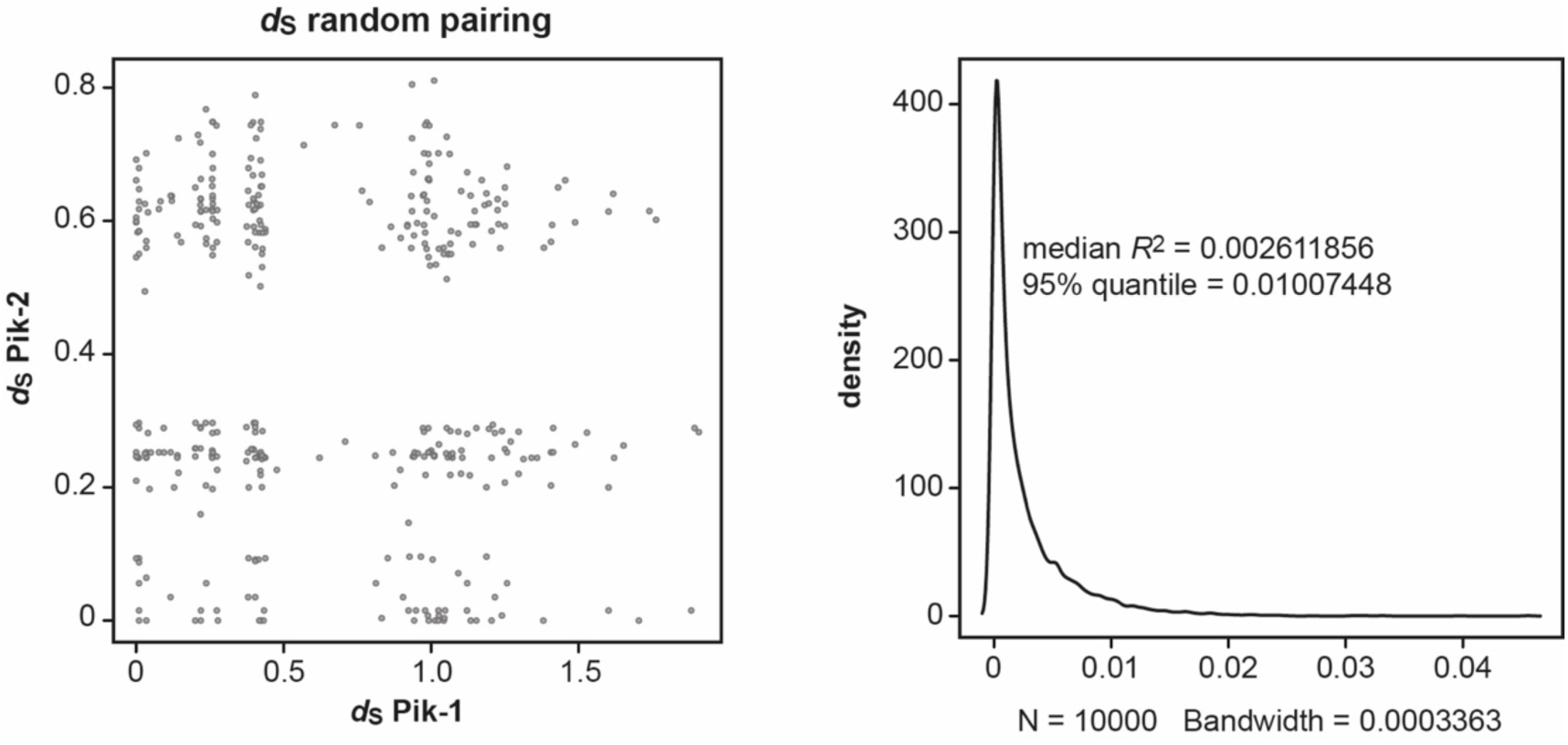
Random pairwise comparisons of *d*_S_ rates calculated for the Pik-1 and Pik-2 receptors. The synonymous (*d*_S_) rates were calculated using Yang & Nielsen method (2000) and presented in Figure 1C. The random datasets for *d*_S_ values were generated by name shuffling in the existing dataset and random sampling from it 1000 times (left panel). The coefficient of determination (*R*^2^) was calculated for every random pairing and the *R*^2^ distribution was plotted (right panel), as implemented in R v3.6.3 package. If less than 5% of the *R*^2^ for the random dataset is bigger than the *R*^2^ for the real dataset, then, according to the null model, the observed difference is very rare and can be accepted as significant with p < 0.05.

**Supplementary figure 6.**
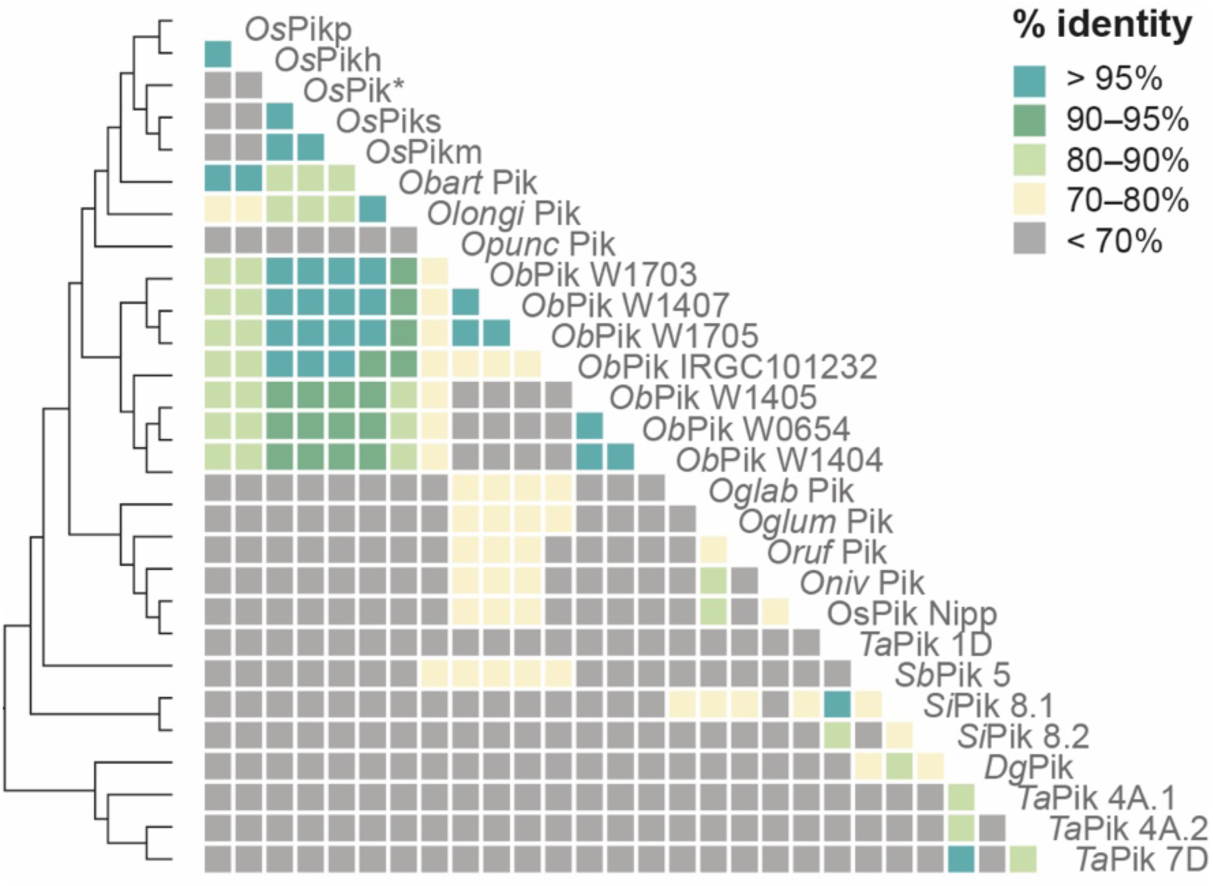
Genetically linked Pik-1 and Pik-2 have similar molecular age. Comparisons of pairwise *d*_S_ rates calculated for the Pik-1 and Pik-2 receptors. The rates were calculated using Yang and Nielsen method (2000) based on 972- and 1269-nucleotide-long codon-based alignments of the NB-ARC domains of Pik-1 and Pik-2, respectively; only positions that showed over 70% coverage across the alignment were used for the analysis. The pairwise comparisons of *d*_S_ rates are presented as a heatmap. The comparisons were ordered based on the Pik-1 phylogenetic relationship, shown on the left. The list of genes used for the pairwise comparisons is summarised in **Supplementary figure 7**.

**Supplementary figure 7.**
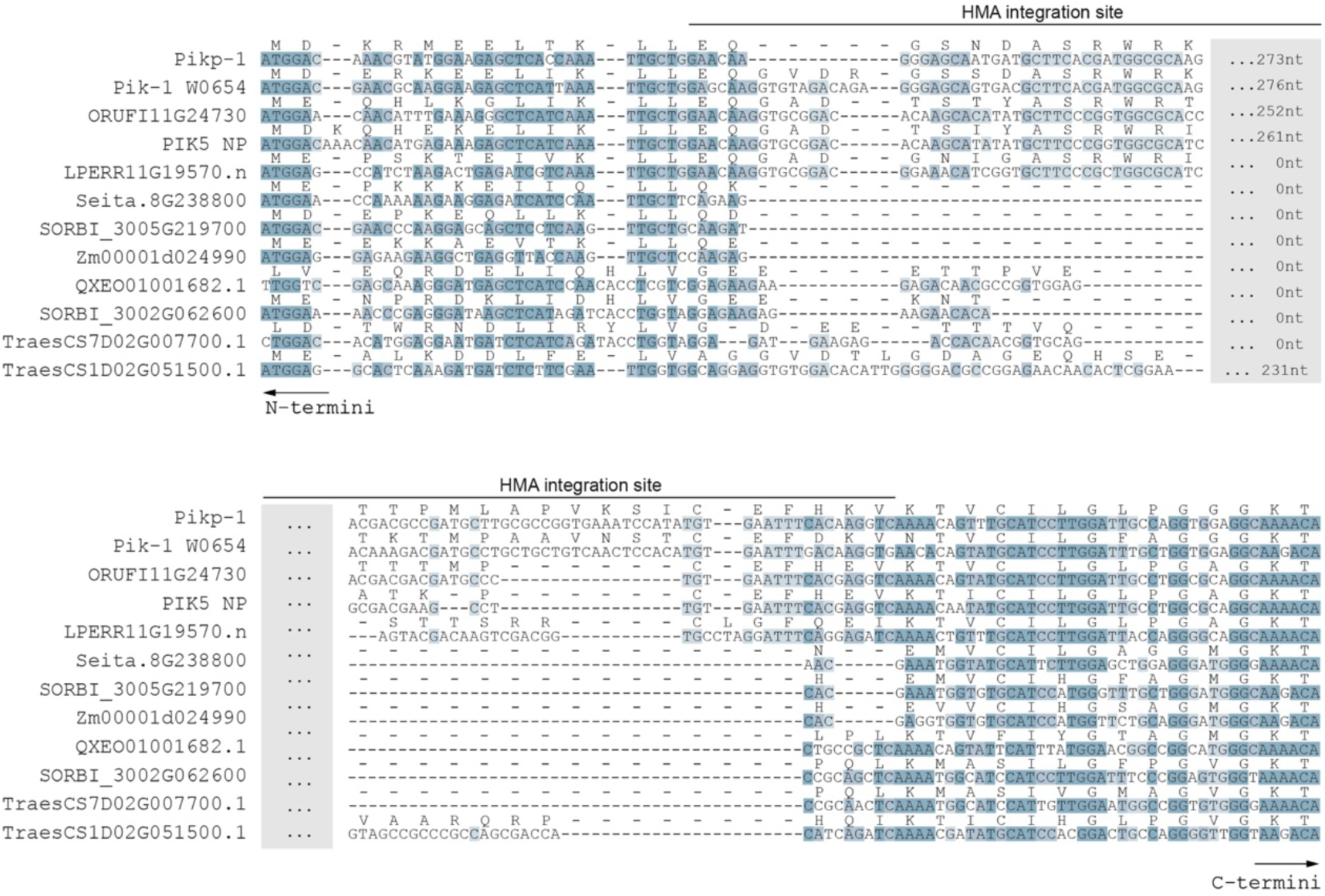
Multiple sequence alignment illustrating the conservation around the HMA integration site. The codon-based sequence alignment of the region surrounding the HMA integration site was generated using MUSCLE (Edgar, 2004). The residues are coloured based on percentage sequence identity from dark (high similarity) to light blue (low similarity).

**Supplementary figure 8.**
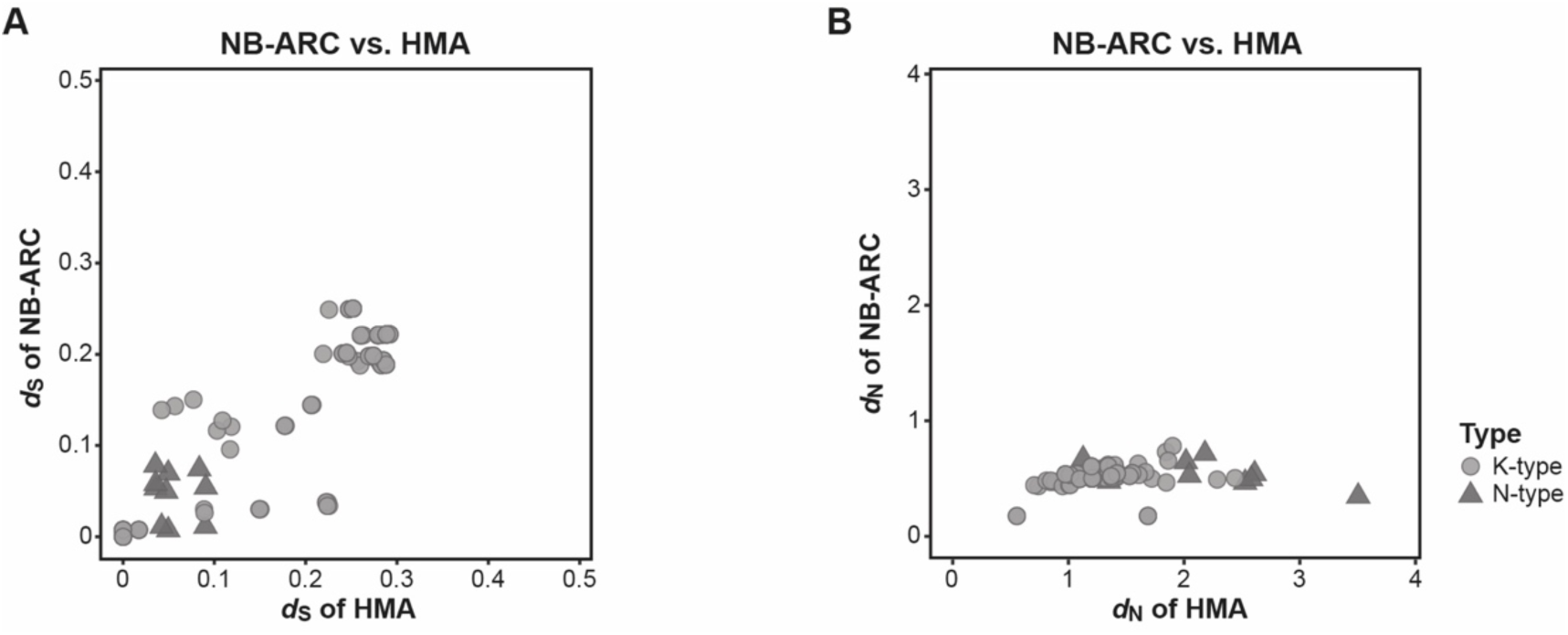
The integrated HMA domain displays elevated rates of *d*_N_ compared with the NB-ARC domain of Pik-1. Pairwise comparison of (**A**) *d*_S_ and (**B**) *d*_N_ rates between the HMA and NB-ARC domains of Pik-1. Pairwise comparison were calculated using Yang and Nielsen method (2000).

**Supplementary table 8.** Summary of the amplification experiment of the Pik-1–integrated HMA domain from wild rice species.

| Accession | Species | Origin | Amplified | Sequence confirmed |
| --- | --- | --- | --- | --- |
| W0654 | *O. brachyantha* | Sierra Leone | Yes | Yes |
| W0008 | *O. australiensis* | Australia (SE Canberra) | Yes | No |
| W1628 | *O. australiensis* | Australia (N) | No | NA |
| W1643 | *O. barthii* | Botswana | Yes | Yes |
| W1605 | *O. barthii* | Nigeria | Yes | No |
| W0042 | *O. barthii* | unspecified | Yes | Yes |
| W0698 | *O. barthii* | Guinea | Yes | Yes |
| W1526 | *O. eichingeri* | Uganda | No | NA |
| W1171 | *O. glumaepatula* | Cuba | No | NA |
| W2203 | *O. glumaepatula* | Brazil (S) | Yes | No |
| W1480(B) | *O. grandiglumis* | Brazil (N) | Yes | No |
| W0005 | *O. granulata* | Sri Lanka | No | NA |
| W0067(B) | *O. granulata* | Thailand | Yes | Yes |
| W0542 | *O. latifolia* / *O. alta* | Mexico | No | NA |
| W1539 | *O. latifolia* / *O. alta* | Argentina (N) | No | NA |
| W1228 | *O. longiglumis* | Singapore (S) | No | NA |
| W1504 | *O. longistaminata* | Tanzania | No | NA |
| W1540 | *O. longistaminata* | Republic of Congo | Yes | Yes |
| W0643 | *O. longistaminata* | The Gambia | Yes | Yes |
| W2081 | *O. meridionalis* | Australia (N) | No | NA |
| W2112 | *O. meridionalis* | Australia (NE) | No | NA |
| W1354 | *O. meyeriana* | Malaysia | Yes | No |
| W1328 | *O. minuta* | Philippines | Yes | Yes |
| W0614 | *O. officinalis* | Myanmar | Yes | Yes |
| W1200 | *O. officinalis* | Philippines | Yes | Yes |
| W1408 | *O. punctata* | Nigeria | Yes | Yes |
| W1514 | *O. punctata* | Kenya | Yes | Yes |
| W1808 | *O. rhizomatis* | Sri Lanka | Yes | No |
| W0001 | *O. ridleyi* | Thailand | Yes | No |
| W2035 | *O. ridleyi* | Philippines | No | NA |
| W2003 | *O. rufipogon* | India (SW) | Yes | Yes |
| W1715 | *O. rufipogon* | Chin (Beijing) | No | NA |
| W2117 | *O. rufipogon*/ *O. meridionalis* | Australia (NE) | Yes | No |

**Supplementary figure 9.**
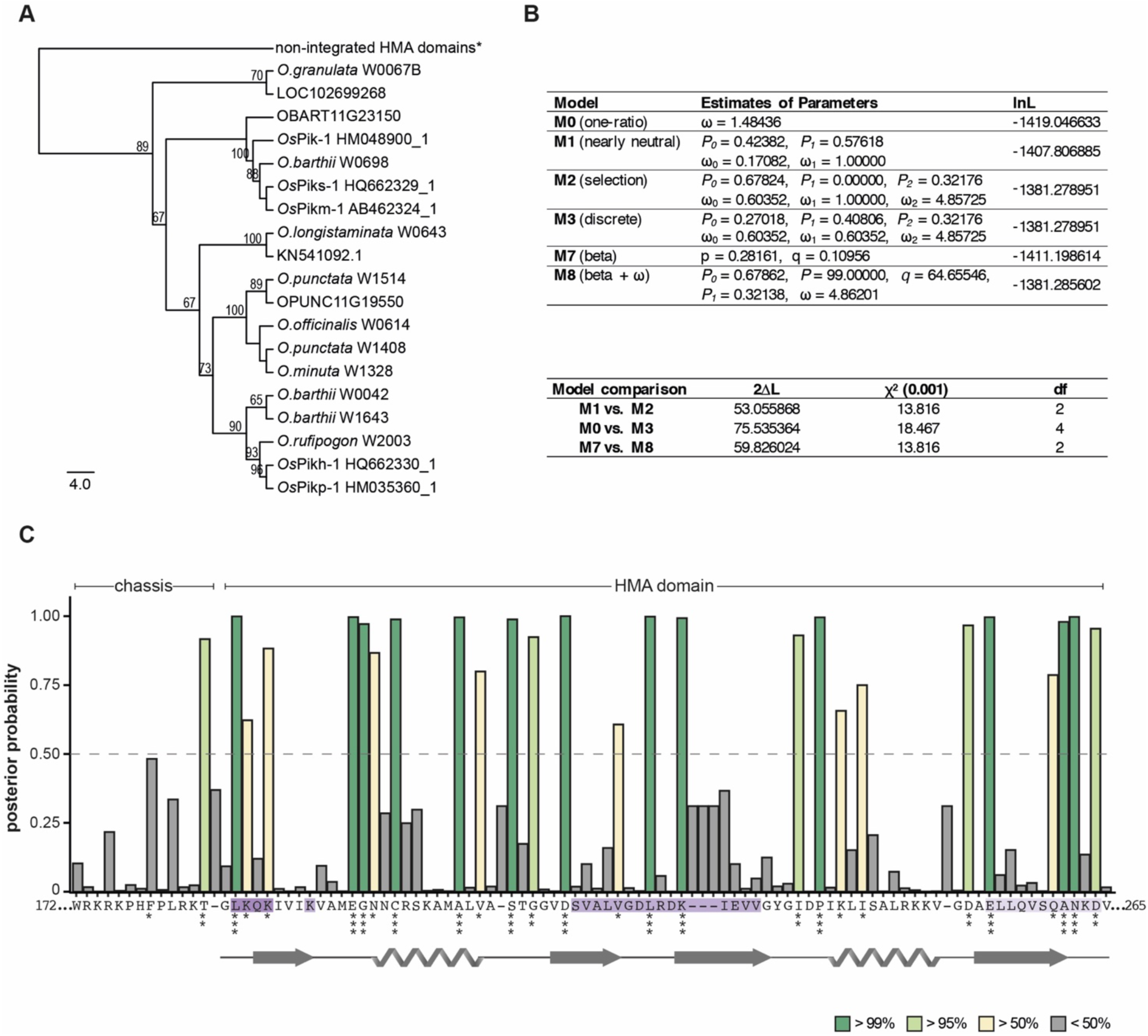
Residues within the integrated HMA domain are likely to have experienced positive selection. (**A**) The NJ tree of the HMA domain calculated using the JTT substitution model (Jones et al., 1992) and bootstrap method with 100 iterations test (Felsenstein, 1985). Alignment of 98 amino acids of integrated HMAs was generated with MUSCLE (Edgar, 2004). Bootstrap values above 65% are shown at the base of respective clades. The scale bar marks the evolutionary distance based on number of base substitutions per site. (*) A branch corresponding to non-integrated HMAs was manually added to the tree to indicate an outgroup, which was used for tree calculation but not for calculating the selection probabilities; the entire tree is presented in **Supplementary figure 12**. (**B**) Results from codon substitution models for heterogeneous selection at amino acid sites (upper panel) and likelihood ratio tests (bottom panel). (**C**) Posterior probabilities for site classes estimated under the beta & ω (M8) model inferred using Bayes empirical Bayes (BEB) (Yang, 2005). The amino acids with higher values of posterior probability are more likely to be under positive selection. The stars indicate potentially positively selected sites: (*) P>50%, (**) P>95%, (***) P>99%. The amino acid sequence and the protein model showed below the plot correspond to Pikp-HMA. The effector-interaction interfaces are marked in shades of purple.

**Supplementary figure 10.**
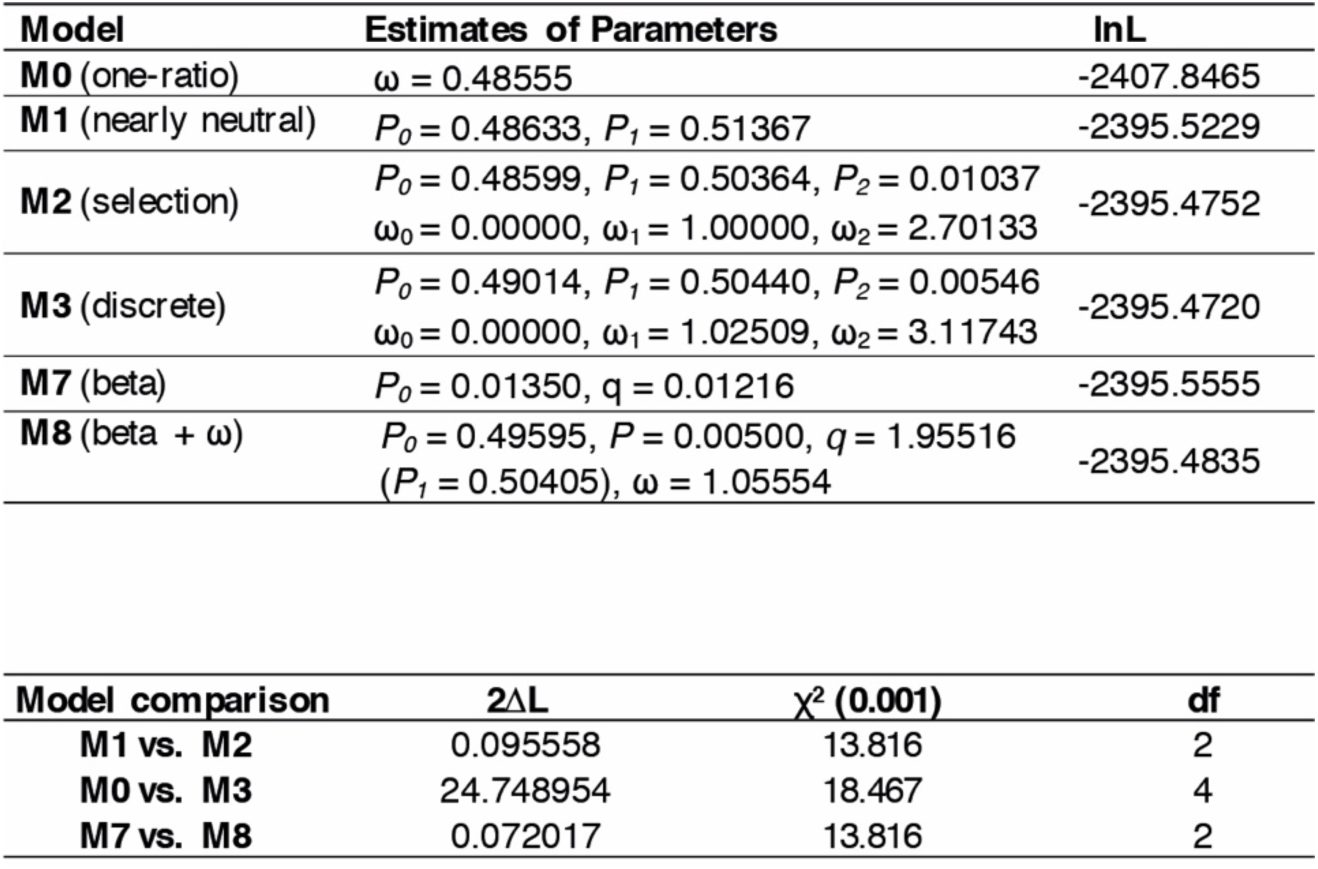
Selection test at the amino acid sites within the NB-ARC domain of the K-type *Pik-1* genes. Results from the codon substitution models for heterogeneous selection at amino acid sites (upper panel) and the likelihood ratio test (bottom panel).

**Supplementary figure 11.**
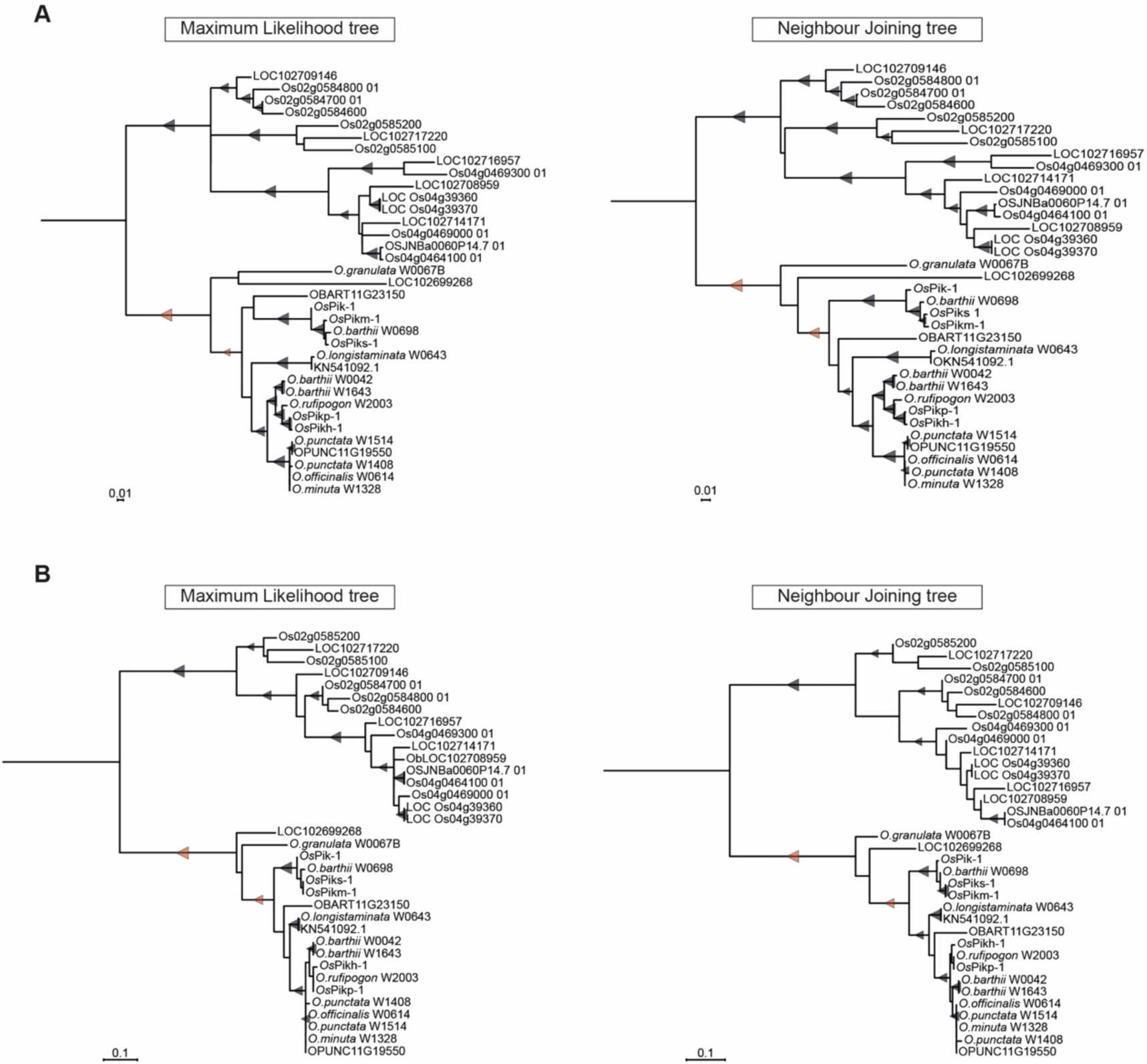
Phylogenetic analyses of the HMA domain of K-type Pik-1 NLRs. The phylogenetic trees were built using MEGA X software (Kumar et al., 2018) and bootstrap method based on 1000 iterations (Felsenstein, 1985). Codon-based 249-nucleotide-long alignment was generated using MUSCLE (Edgar, 2004); positions with less than 50% site coverage were removed prior the analysis, resulting in 234 positions in the final dataset. The relevant bootstrap values with support over 60% are shown with triangles at the base of representative clades; the size of the triangle is proportional to the bootstrap value. The scale bars indicate the evolutionary distance based on nucleotide substitution rate. Each tree was manually rooted using a clade of non-integrated HMA as an outgroup. The nodes selected for the ancestral sequence reconstruction are marked with red triangles. (**A**) ML and NJ trees calculated based on all codon positions in the alignment. (**B**) ML and NJ trees calculated based on third codon position in the alignment.

**Supplementary figure 12.**
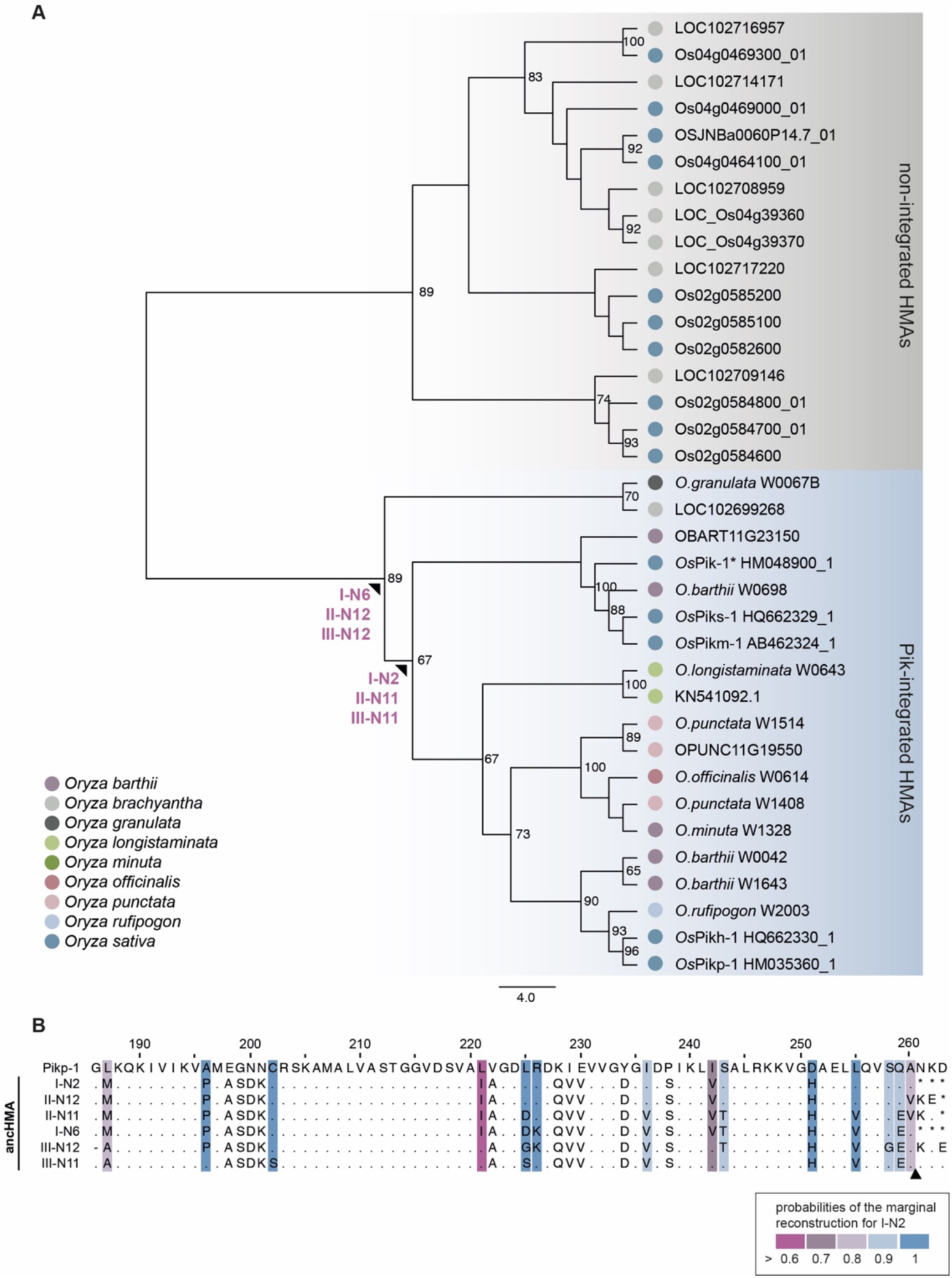
Ancestral sequence reconstruction yielded multiple plausible ancHMA sequences. (**A**) Representative NJ phylogenetic tree of the HMA domain. The tree was built using JTT substitution model (Jones et al., 1992) and bootstrap method with 100 iterations test (Felsenstein, 1985). Alignment of 98 amino acids of integrated (blue) and non-integrated (grey) HMAs was generated with MUSCLE (Edgar, 2004). Bootstrap values above 65% were shown at the base of respective clades. Nodes for which the ancestral sequence reconstruction was performed were marked with arrowheads. The scale bar indicates the evolutionary distance based on number of base substitutions per site. (**B**) Protein sequence alignment of representative ancestral HMA predictions. Amino acids for which sequence prediction was not performed were replaced with asterisk (*). The probabilities of the marginal reconstruction for I-N2 sequence were marked with coloured boxes. An arrowhead indicates the length of the construct used in further studies concerning Pikp-1.

**Supplementary figure 13.**
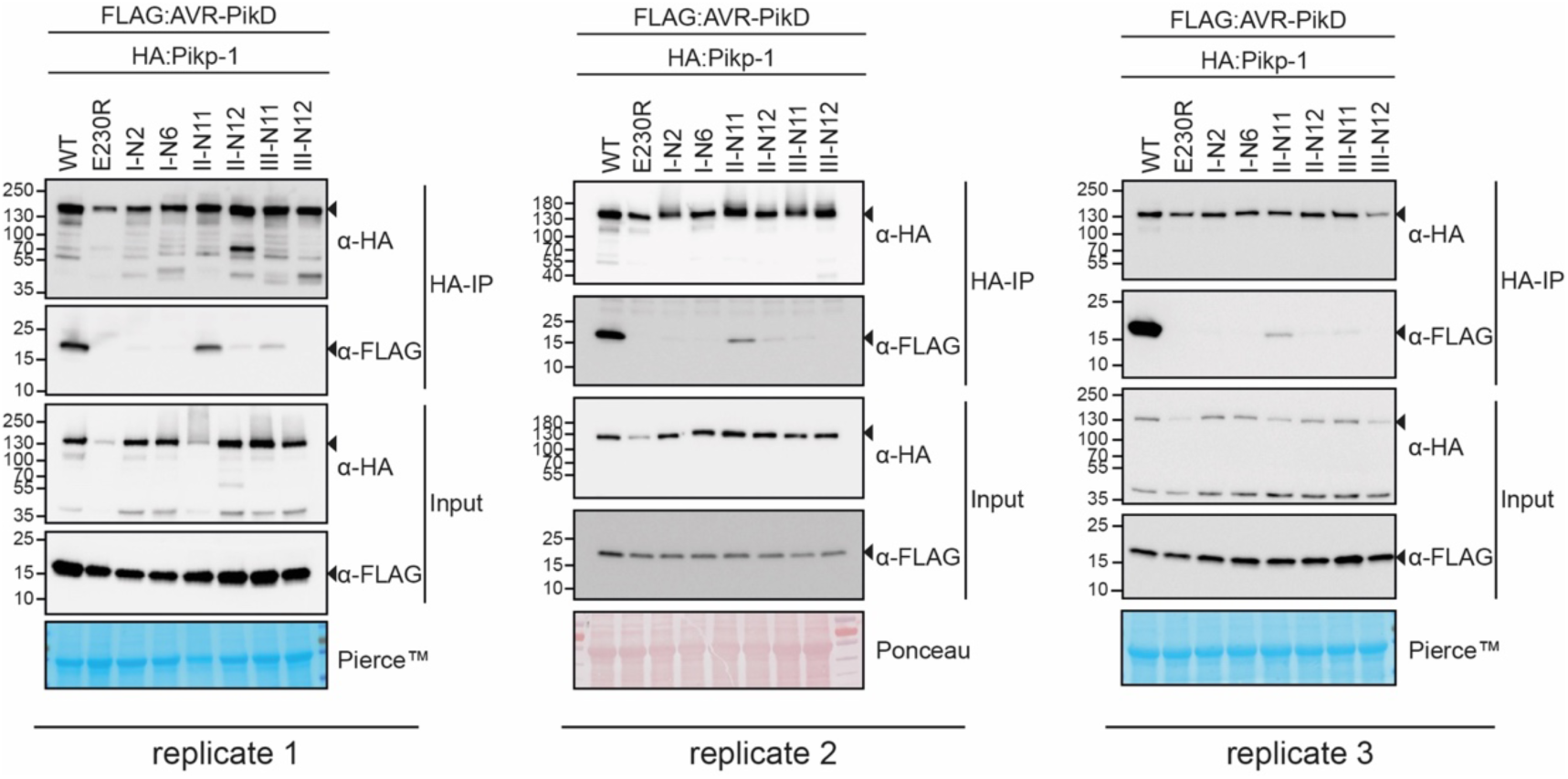
Replicates of the co-IP experiment between AVR-PikD and the reconstructed ancHMA sequences. Co-IP experiment between AVR-PikD (N-terminally tagged with FLAG) with Pikp-1 with ancestral sequences of HMA (N-terminally tagged with HA). Wild-type (WT) HA:Pikp-1 and HA:Pikp-1_E230R_ were used as positive and negative controls, respectively. Immunoprecipitates (HA-IP) o obtained using anti-HA probes and total protein extracts (Input) were immunoblotted with appropriate antisera (listed on the right). Rubisco loading control was performed using Pierce^TM^ or Ponceau staining solutions. Arrowheads indicate expected band sizes. The figure shows results from three independent experiments.

**Supplementary figure 14.**
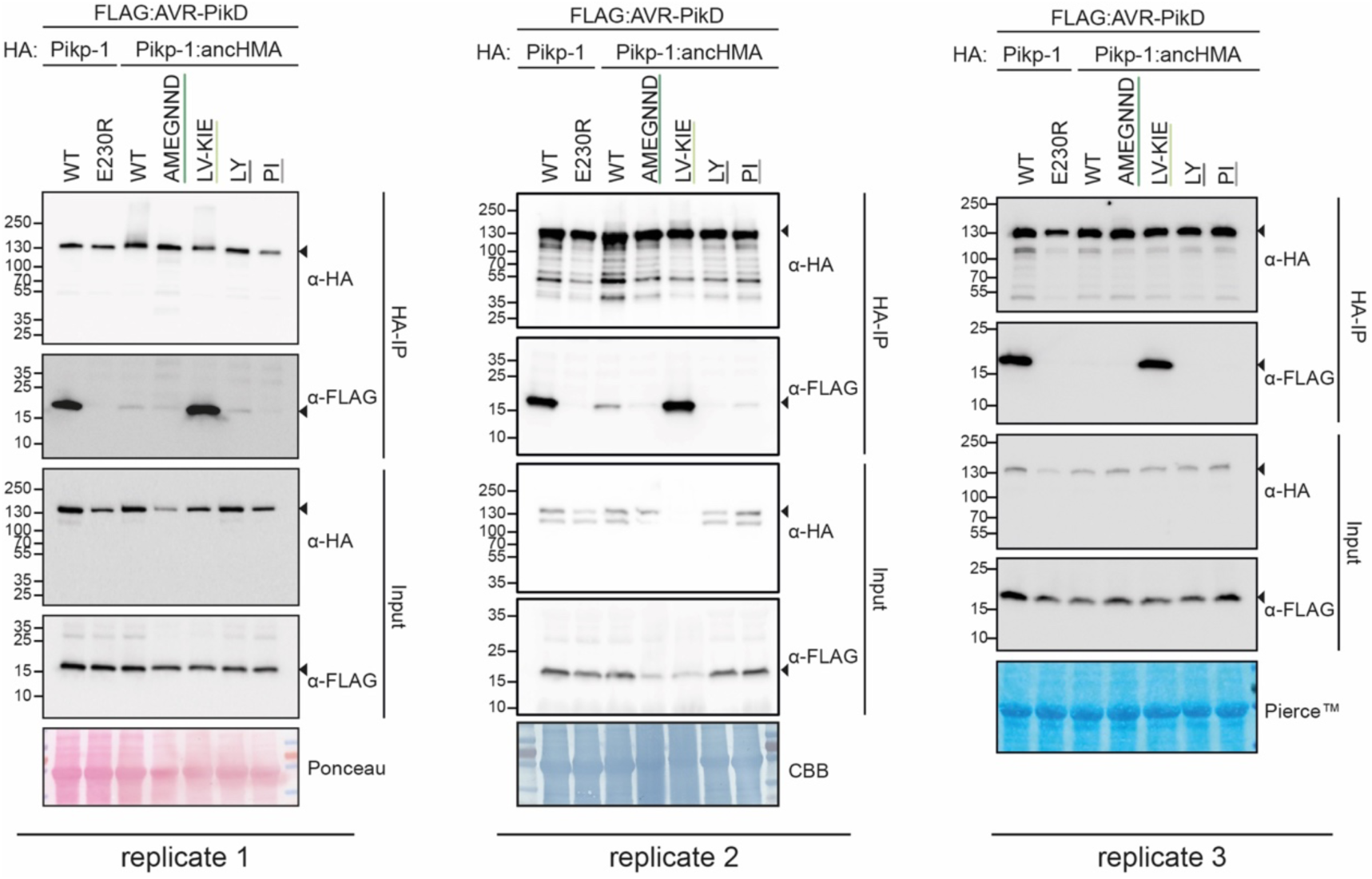
Replicates of the co-IP experiment between AVR-PikD and the Pikp-1:ancHMA chimeras. Association of AVR-PikD (N-terminally tagged with FLAG) with Pikp-1, Pikp-1_E230R_, Pikp-1:ancHMA, and Pikp-1:ancHMA chimeras (N-terminally tagged with HA), labelled above, was tested in planta by co-IP. Wild-type (WT) HA:Pikp-1 and HA:Pikp-1_E230R_ were used as a positive and negative control, respectively. Immunoprecipitates (HA-IP) obtained using anti-HA probes and total protein extracts (Input) were immunoblotted with appropriate antisera, labelled on the right. Arrowheads show expected band sizes. Rubisco loading controls were performed using Pierce^TM^, Coomassie Brilliant Blue (CBB), or Ponceau staining solutions. The figure shows results from three independent experiments.

**Supplementary figure 15.**
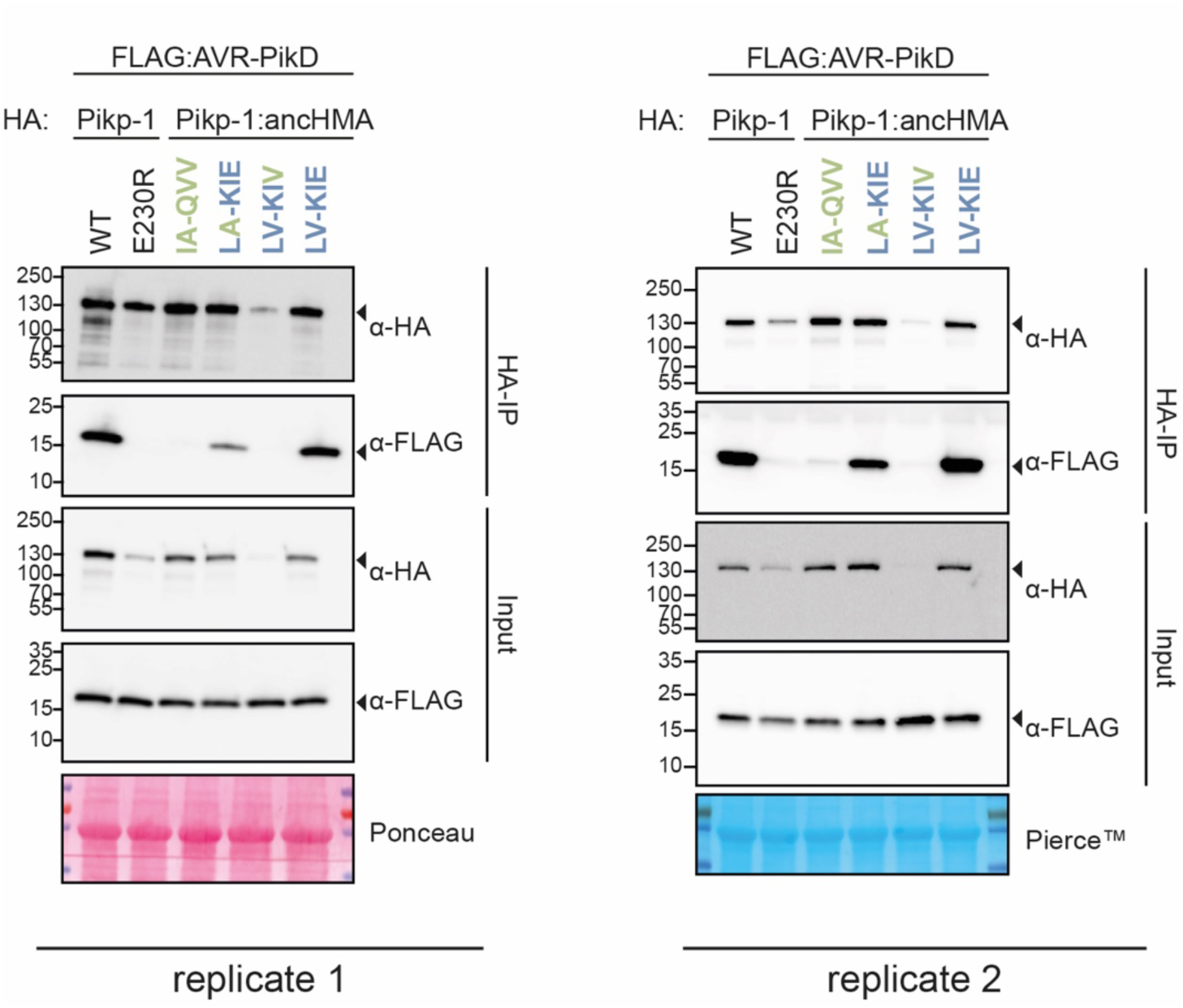
Co-IP experiment between AVR-PikD and the two plausible historical states of the IAQVV/LVKIE region within Pikp-HMA. In planta association of AVR-PikD (N-terminally tagged with FLAG) Pikp-1, Pikp-1_E230R_, Pikp-1:ancHMA, and Pikp-1:ancHMA mutants (N-terminally tagged with HA), labelled above. Wild-type (WT) HA:Pikp-1 and HA:Pikp-1_E230R_, with HA tag, were used as a positive and negative control, respectively. Immunoprecipitates (HA-IP) obtained using anti-HA probe and total protein extracts (Input) were immunoblotted with the appropriate antisera labelled on the right. Arrowheads indicate expected band sizes. Loading controls, featuring Rubisco, were performed using Pierce^TM^ or Ponceau staining solutions. The figure shows results from two independent experiments.

**Supplementary figure 16.**
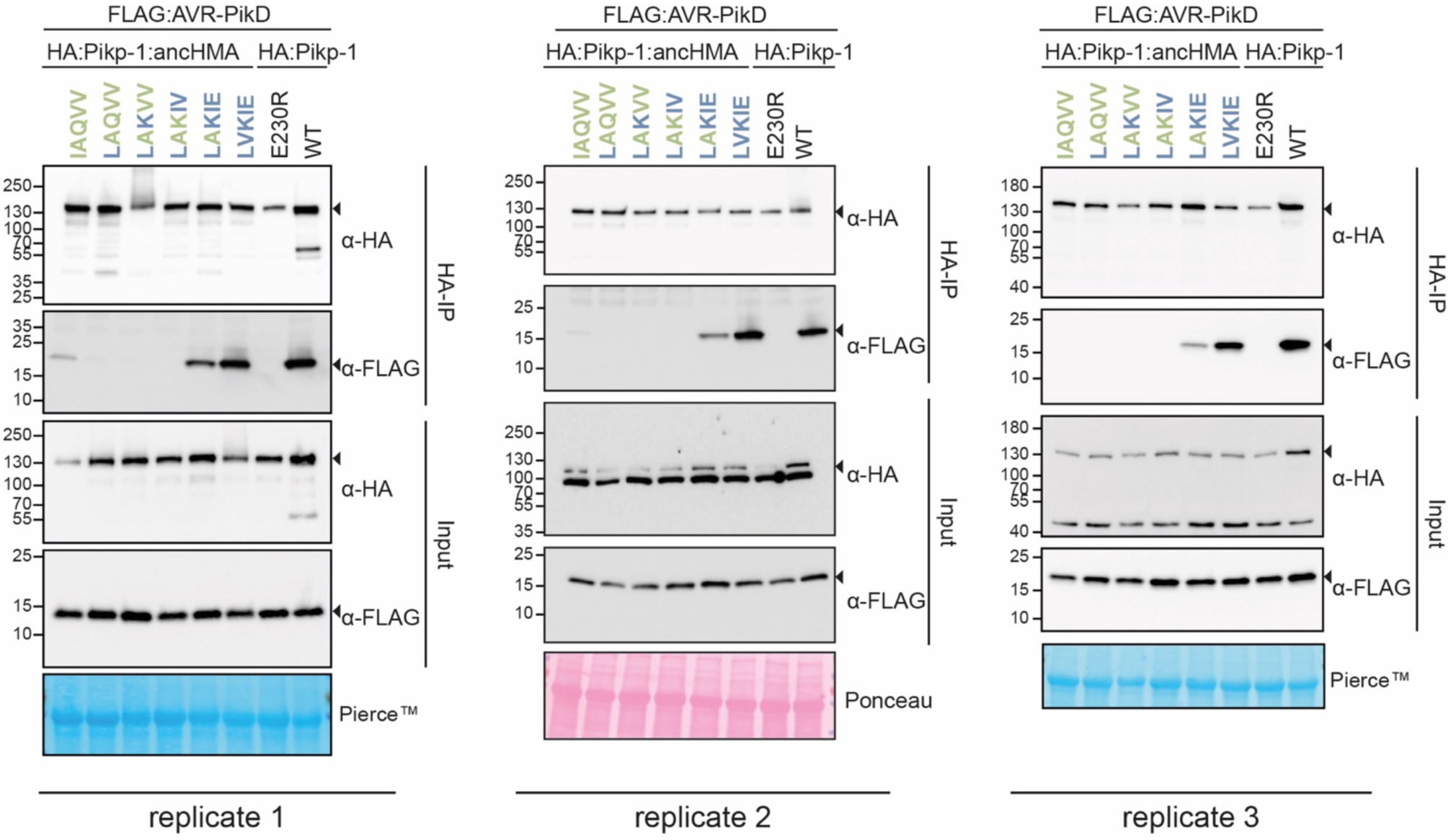
Replicates of the co-IP experiments between the Pikp-1:ancHMA IAQVV/LVKIE mutants and AVR-PikD. In planta association of AVR-PikD (N-terminally tagged with FLAG) with Pikp-1, Pikp-1_E230R_, Pikp-1:ancHMA, and Pikp-1:ancHMA mutants (N-terminally tagged with HA), labelled above. Wild-type (WT) Pikp-1 and Pikp-1_E230R_ were used as a positive and negative control, respectively. Proteins obtained by co-IP with HA-probe (HA-IP) and total protein extracts (Input) were immunoblotted with the appropriate antisera labelled on the right. Rubisco loading controls were conducted using Pierce^TM^, Coomassie Brilliant Blue (CBB), or Ponceau staining solutions. Arrowheads demonstrate expected band sizes. The figure shows results from three independent experiments.

**Supplementary figure 17.**
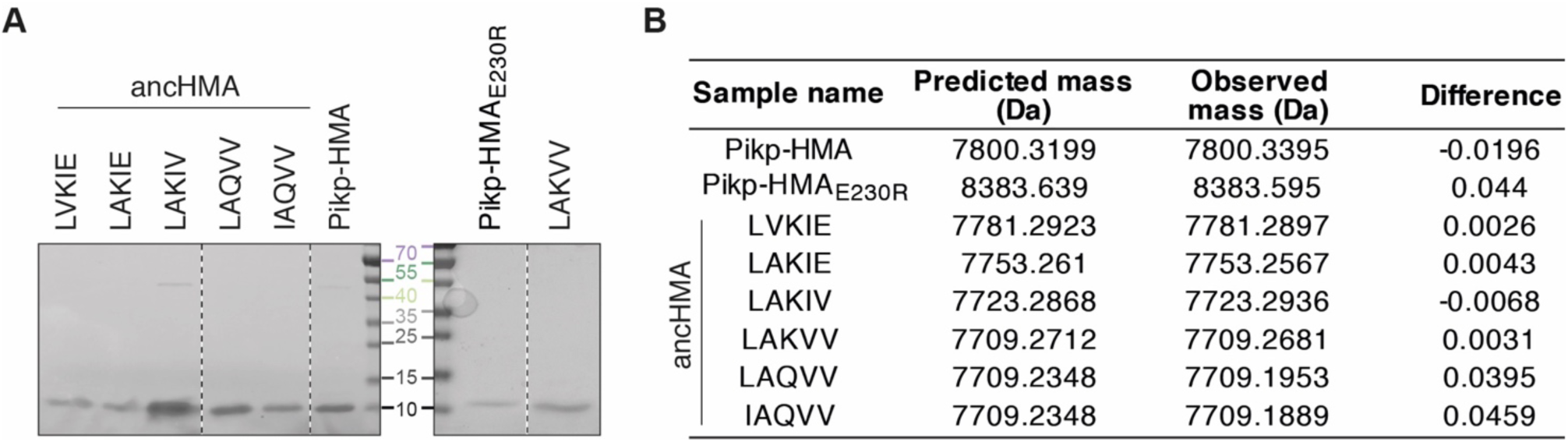
Purified proteins used in SPR studies. (**A**) Coomasie Brilliant Blue–stained SDS-PAGE gel showing purified HMA proteins used in in vitro experiments. Dashed lines signify different components of the same gel. (**B**) Table summarising intact masses (monoisotopic) of proteins from panel A.

**Supplementary figure 18.**
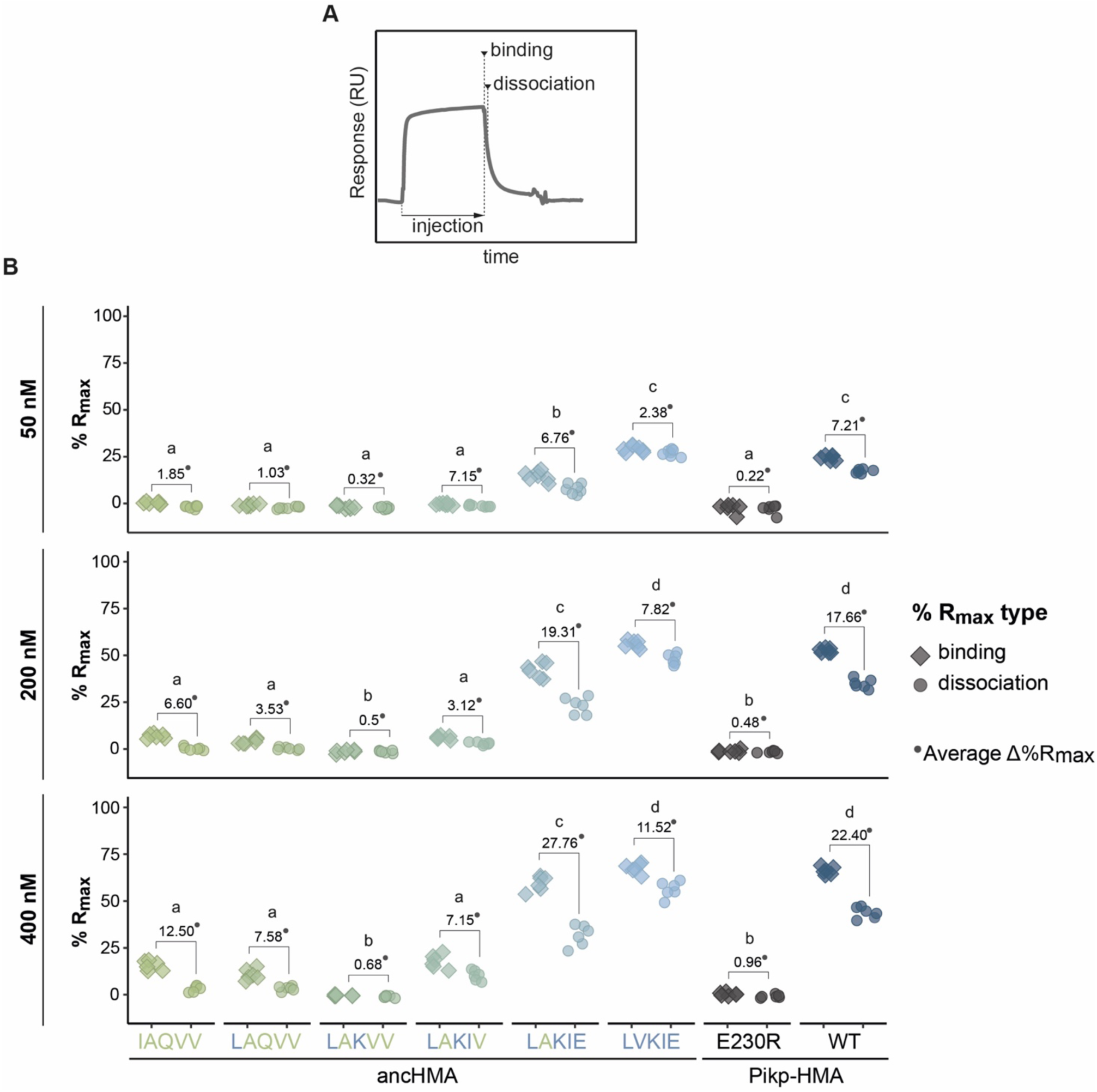
SPR results show the effect of the IAQVV-LVKIE mutations on the AVR-PikD binding, as indicated by %R_max_. (**A**) Schematic representation of the SPR sensorgrams showcasing the measurements taken to monitor binding dynamics: ‘binding’ and ‘dissociation’. (**B**) Plots illustrating calculated percentage of the theoretical maximum response values (%R_max_) for interaction of the HMA analytes, labelled below, with AVR-PikD ligand (C-terminally tagged with HIS). %R_max_ was normalized for the amount of ligand immobilized on the NTA-sensor chip. The HMA analytes were tested at three different concentrations, indicated on the left, in three independent experiments with two internal replicates. All data points are represented as diamonds or circles. Average Δ%R_max_ (•) values represent absolute differences between values for ‘binding’ and ‘dissociation’, calculated from average values for each sample, and serve as an off-rate approximate. Statistical differences among the samples were analysed with ANOVA and Tukey’s honest significant difference (HSD) test (p < 0.01); p-values for all pairwise comparisons are presented in **Supplementary table 9**.

**Supplementary table 9.** Table of p-values for all pairwise comparisons of SPR binding to AVR-PikD between the HMA mutants.

| Concentration (nM) | Sample 1 | Sample 2 | Difference | Lower confidence level | Upper confidence level | p-value |
| --- | --- | --- | --- | --- | --- | --- |
| 50 | IAQVV | E230R | 0.7594754 | -6.380544 | 7.899495 | 0.9999725 |
| 50 | LAKIE | E230R | 15.1954147 | 8.055395 | 22.335434 | 0.0000005 |
| 50 | LAKIV | E230R | 0.1973327 | -6.942687 | 7.337352 | 1 |
| 50 | LAKVV | E230R | -1.2876815 | -8.427701 | 5.852338 | 0.9990674 |
| 50 | LAQVV | E230R | -0.2110851 | -7.351104 | 6.928934 | 1 |
| 50 | LVKIE | E230R | 29.9244059 | 22.784387 | 37.064425 | 0 |
| 50 | Pikp-HMA | E230R | 21.3309677 | 14.190948 | 28.470987 | 0 |
| 50 | LAKIE | IAQVV | 14.4359393 | 7.29592 | 21.575959 | 0.0000016 |
| 50 | LAKIV | IAQVV | -0.5621428 | -7.702162 | 6.577877 | 0.9999965 |
| 50 | LAKVV | IAQVV | -2.047157 | -9.187176 | 5.092862 | 0.9838216 |
| 50 | LAQVV | IAQVV | -0.9705606 | -8.11058 | 6.169459 | 0.9998555 |
| 50 | LVKIE | IAQVV | 29.1649305 | 22.024911 | 36.30495 | 0 |
| 50 | Pikp-HMA | IAQVV | 20.5714923 | 13.431473 | 27.711512 | 0 |
| 50 | LAKIV | LAKIE | -14.998082 | -22.138101 | -7.858063 | 0.0000007 |
| 50 | LAKVV | LAKIE | -16.483096 | -23.623116 | -9.343077 | 0.0000001 |
| 50 | LAQVV | LAKIE | -15.4065 | -22.546519 | -8.266481 | 0.0000004 |
| 50 | LVKIE | LAKIE | 14.7289911 | 7.588972 | 21.86901 | 0.000001 |
| 50 | Pikp-HMA | LAKIE | 6.1355529 | -1.004466 | 13.275572 | 0.1407186 |
| 50 | LAKVV | LAKIV | -1.4850142 | -8.625033 | 5.655005 | 0.9976792 |
| 50 | LAQVV | LAKIV | -0.4084178 | -7.548437 | 6.731601 | 0.9999996 |
| 50 | LVKIE | LAKIV | 29.7270732 | 22.587054 | 36.867093 | 0 |
| 50 | Pikp-HMA | LAKIV | 21.133635 | 13.993616 | 28.273654 | 0 |
| 50 | LAQVV | LAKVV | 1.0765964 | -6.063423 | 8.216616 | 0.9997116 |
| 50 | LVKIE | LAKVV | 31.2120874 | 24.072068 | 38.352107 | 0 |
| 50 | Pikp-HMA | LAKVV | 22.6186492 | 15.47863 | 29.758669 | 0 |
| 50 | LVKIE | LAQVV | 30.135491 | 22.995472 | 37.27551 | 0 |
| 50 | Pikp-HMA | LAQVV | 21.5420528 | 14.402034 | 28.682072 | 0 |
| 50 | Pikp-HMA | LVKIE | -8.5934382 | -15.733457 | -1.453419 | 0.008645 |
| 200 | IAQVV | E230R | 8.12408613 | 4.769044 | 11.4791283 | 0 |
| 200 | LAKIE | E230R | 43.4535131 | 40.098471 | 46.8085553 | 0 |
| 200 | LAKIV | E230R | 7.26935912 | 3.914317 | 10.6244013 | 0.0000006 |
| 200 | LAKVV | E230R | -0.0357561 | -3.390798 | 3.3192861 | 1 |
| 200 | LAQVV | E230R | 4.96818917 | 1.613147 | 8.3232314 | 0.0006703 |
| 200 | LVKIE | E230R | 57.0910393 | 53.735997 | 60.4460815 | 0 |
| 200 | Pikp-HMA | E230R | 53.8341313 | 50.479089 | 57.1891735 | 0 |
| 200 | LAKIE | IAQVV | 35.329427 | 31.974385 | 38.6844692 | 0 |
| 200 | LAKIV | IAQVV | -0.854727 | -4.209769 | 2.5003152 | 0.9912868 |
| 200 | LAKVV | IAQVV | -8.1598423 | -11.514884 | -4.8048001 | 0 |
| 200 | LAQVV | IAQVV | -3.155897 | -6.510939 | 0.1991452 | 0.0782897 |
| 200 | LVKIE | IAQVV | 48.9669532 | 45.611911 | 52.3219953 | 0 |
| 200 | Pikp-HMA | IAQVV | 45.7100452 | 42.355003 | 49.0650873 | 0 |
| 200 | LAKIV | LAKIE | -36.184154 | -39.539196 | -32.829112 | 0 |
| 200 | LAKVV | LAKIE | -43.489269 | -46.844311 | -40.134227 | 0 |
| 200 | LAQVV | LAKIE | -38.485324 | -41.840366 | -35.130282 | 0 |
| 200 | LVKIE | LAKIE | 13.6375262 | 10.282484 | 16.9925683 | 0 |
| 200 | Pikp-HMA | LAKIE | 10.3806182 | 7.025576 | 13.7356603 | 0 |
| 200 | LAKVV | LAKIV | -7.3051152 | -10.660157 | -3.9500731 | 0.0000006 |
| 200 | LAQVV | LAKIV | -2.30117 | -5.656212 | 1.0538722 | 0.3776686 |
| 200 | LVKIE | LAKIV | 49.8216802 | 46.466638 | 53.1767223 | 0 |
| 200 | Pikp-HMA | LAKIV | 46.5647722 | 43.20973 | 49.9198144 | 0 |
| 200 | LAQVV | LAKVV | 5.00394529 | 1.648903 | 8.3589875 | 0.0006038 |
| 200 | LVKIE | LAKVV | 57.1267954 | 53.771753 | 60.4818376 | 0 |
| 200 | Pikp-HMA | LAKVV | 53.8698874 | 50.514845 | 57.2249296 | 0 |
| 200 | LVKIE | LAQVV | 52.1228501 | 48.767808 | 55.4778923 | 0 |
| 200 | Pikp-HMA | LAQVV | 48.8659421 | 45.5109 | 52.2209843 | 0 |
| 200 | Pikp-HMA | LVKIE | -3.256908 | -6.61195 | 0.0981342 | 0.0625642 |
| 400 | IAQVV | E230R | 15.4160802 | 10.476265 | 20.3558957 | 0 |
| 400 | LAKIE | E230R | 59.1376719 | 54.197856 | 64.0774874 | 0 |
| 400 | LAKIV | E230R | 17.2208188 | 12.281003 | 22.1606344 | 0 |
| 400 | LAKVV | E230R | -0.6186041 | -5.55842 | 4.3212114 | 0.9999089 |
| 400 | LAQVV | E230R | 10.5836822 | 5.643867 | 15.5234977 | 0.0000008 |
| 400 | LVKIE | E230R | 67.4451738 | 62.505358 | 72.3849893 | 0 |
| 400 | Pikp-HMA | E230R | 66.0719577 | 61.132142 | 71.0117733 | 0 |
| 400 | LAKIE | IAQVV | 43.7215917 | 38.781776 | 48.6614072 | 0 |
| 400 | LAKIV | IAQVV | 1.8047387 | -3.135077 | 6.7445542 | 0.9363629 |
| 400 | LAKVV | IAQVV | -16.034684 | -20.9745 | -11.094869 | 0 |
| 400 | LAQVV | IAQVV | -4.832398 | -9.772213 | 0.1074176 | 0.0591016 |
| 400 | LVKIE | IAQVV | 52.0290936 | 47.089278 | 56.9689091 | 0 |
| 400 | Pikp-HMA | IAQVV | 50.6558776 | 45.716062 | 55.5956931 | 0 |
| 400 | LAKIV | LAKIE | -41.916853 | -46.856669 | -36.977038 | 0 |
| 400 | LAKVV | LAKIE | -59.756276 | -64.696092 | -54.81646 | 0 |
| 400 | LAQVV | LAKIE | -48.55399 | -53.493805 | -43.614174 | 0 |
| 400 | LVKIE | LAKIE | 8.3075019 | 3.367686 | 13.2473174 | 0.0000904 |
| 400 | Pikp-HMA | LAKIE | 6.9342859 | 1.99447 | 11.8741014 | 0.001416 |
| 400 | LAKVV | LAKIV | -17.839423 | -22.779238 | -12.899607 | 0 |
| 400 | LAQVV | LAKIV | -6.6371366 | -11.576952 | -1.6973211 | 0.0025111 |
| 400 | LVKIE | LAKIV | 50.2243549 | 45.284539 | 55.1641705 | 0 |
| 400 | Pikp-HMA | LAKIV | 48.8511389 | 43.911323 | 53.7909544 | 0 |
| 400 | LAQVV | LAKVV | 11.2022863 | 6.262471 | 16.1421019 | 0.0000002 |
| 400 | LVKIE | LAKVV | 68.0637779 | 63.123962 | 73.0035934 | 0 |
| 400 | Pikp-HMA | LAKVV | 66.6905618 | 61.750746 | 71.6303774 | 0 |
| 400 | LVKIE | LAQVV | 56.8614916 | 51.921676 | 61.8013071 | 0 |
| 400 | Pikp-HMA | LAQVV | 55.4882755 | 50.54846 | 60.4280911 | 0 |
| 400 | Pikp-HMA | LVKIE | -1.373216 | -6.313032 | 3.5665995 | 0.9854733 |

E230R: Pikp-HMA<sub>E230R</sub>, IAQVV: anchMA<sub>IAQVV</sub>, LAQVV: anchMA<sub>LAQVV</sub>, LAKVV: anchMA<sub>LAKVV</sub>, LAKIV: anchMA<sub>LAKIV</sub>, LAKIE: anchMA<sub>LAKIE</sub>, LVKIE: anchMA<sub>LVKIE</sub>

**Supplementary table 10.**
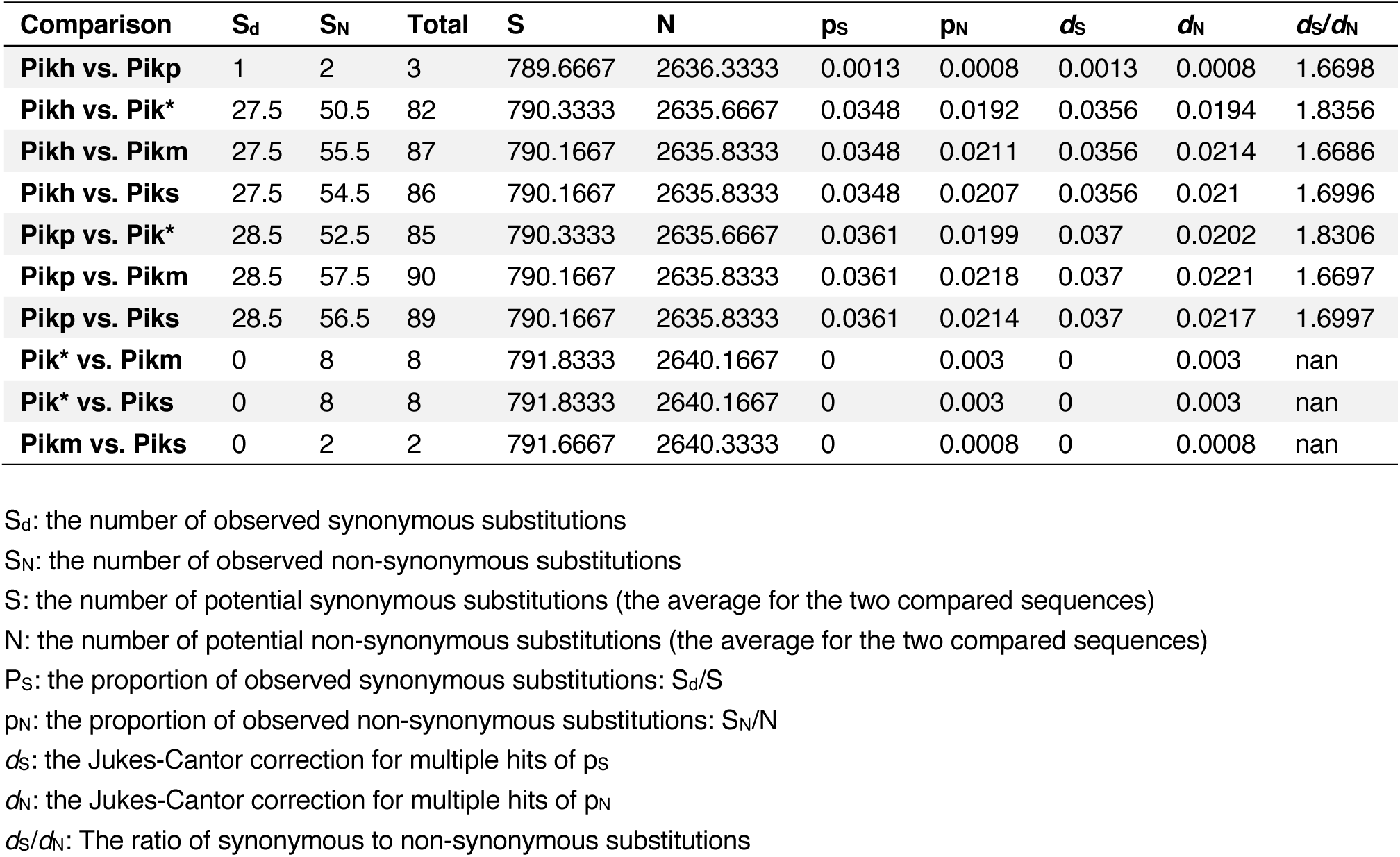
Pairwise *d*_N_ and *d*_S_ values between Pik-1 alleles from rice calculated using the method of Nei and Gojobori (1986).

**Supplementary figure 19.**
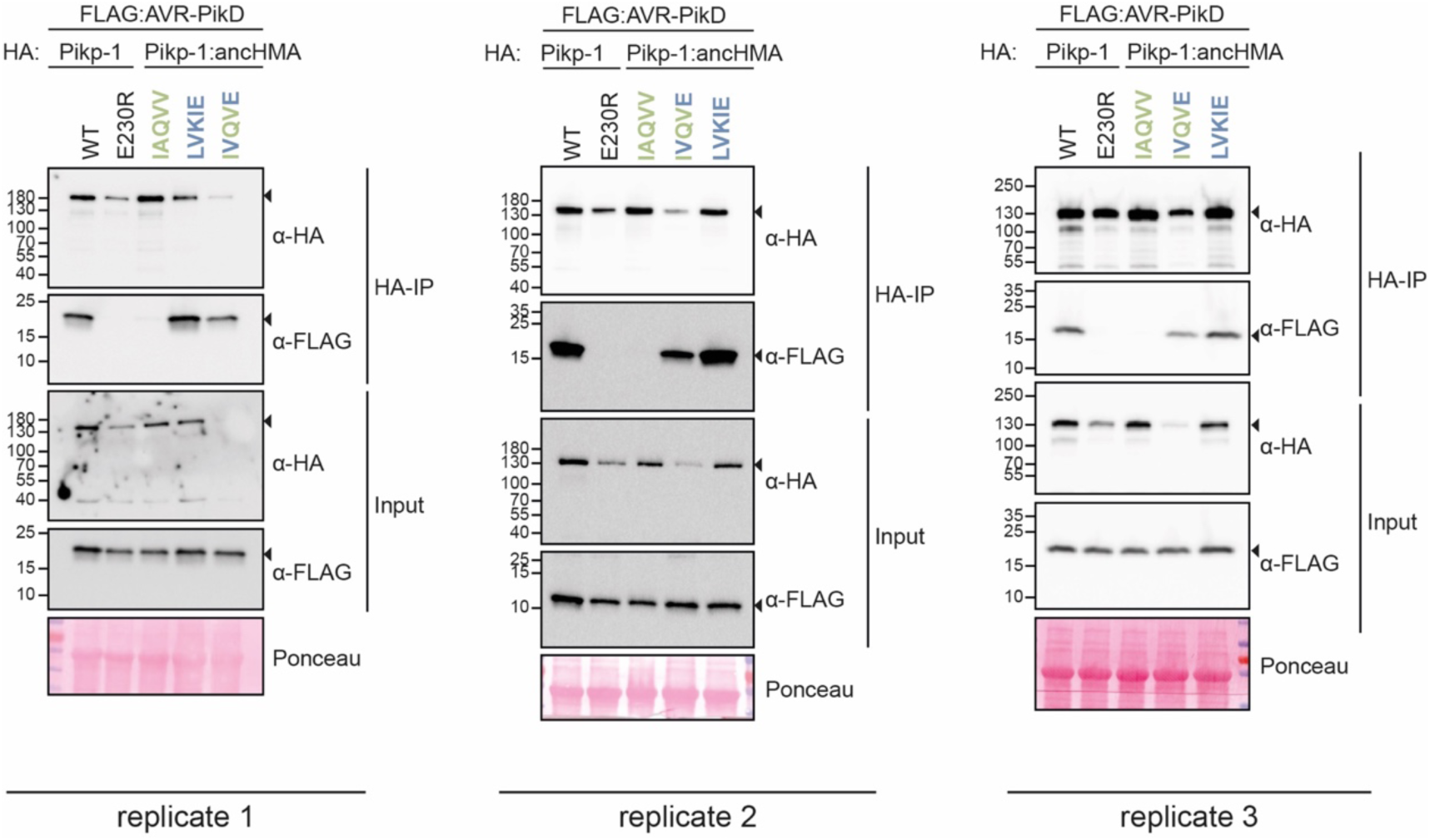
The AV-VE (Ala-222-Val and Val-230-Glu) substitutions are sufficient to increase binding affinity towards the AVR-PikD effector in co-IP. Co-IP experiments between AVR-PikD (N-terminally tagged with FLAG) and Pikp-1 and Pikp-1:ancHMA constructs (N-terminally tagged with HA), labelled above. Wild-type (WT) HA:Pikp-1 and HA:Pikp-1_E230R_ mutant were used as a positive and negative control, respectively. Immunoprecipitates (HA-IP) obtained using anti-HA resin and total protein extracts (Input) were immunoblotted with the appropriate antisera labelled on the right. Loading control, featuring rubisco, was performed using Ponceau staining. The black arrowheads point to expected band sizes. The figure shows results from three independent experiments.

**Supplementary figure 20.**
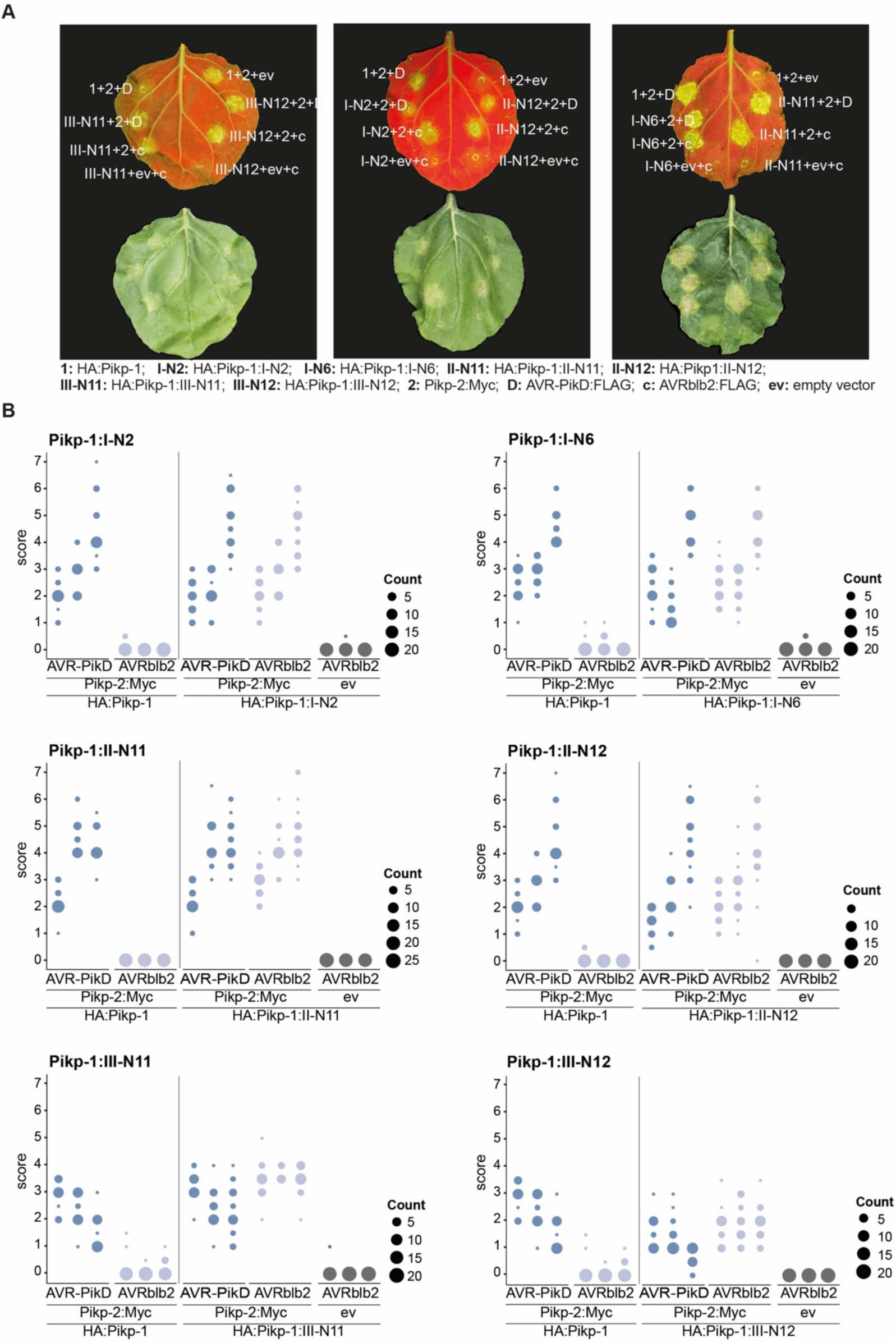
Pikp-1:ancHMA fusions are autoactive in Pikp-2–dependent manner. HR assay after transient co-expression of Pikp-1:HMA variants (N-terminally tagged with HA) with AVR-PikD (N-terminally tagged with FLAG) and Pikp-2 (C-terminally tagged with Myc). AVRblb2 and the empty vector (ev) were used as negative controls. (**A**) Representative *N. benthamiana* leaves infiltrated with samples (labelled next to the infiltration spot) were photographed five days post infiltration under UV (left) and daylight (right). (**B**) HR was scored five days after agroinfiltration. The results are presented as dots plot, where the size of a dot is proportional to the number of samples with the same score (count) within the same replicate. The experiment was repeated at least three times with 22–26 internal replicates; the columns within tested conditions (labelled on the bottom) show results from different biological replicates. The statistical analyses of these results are presented in **Supplementary figure 21**.

**Supplementary figure 21.**
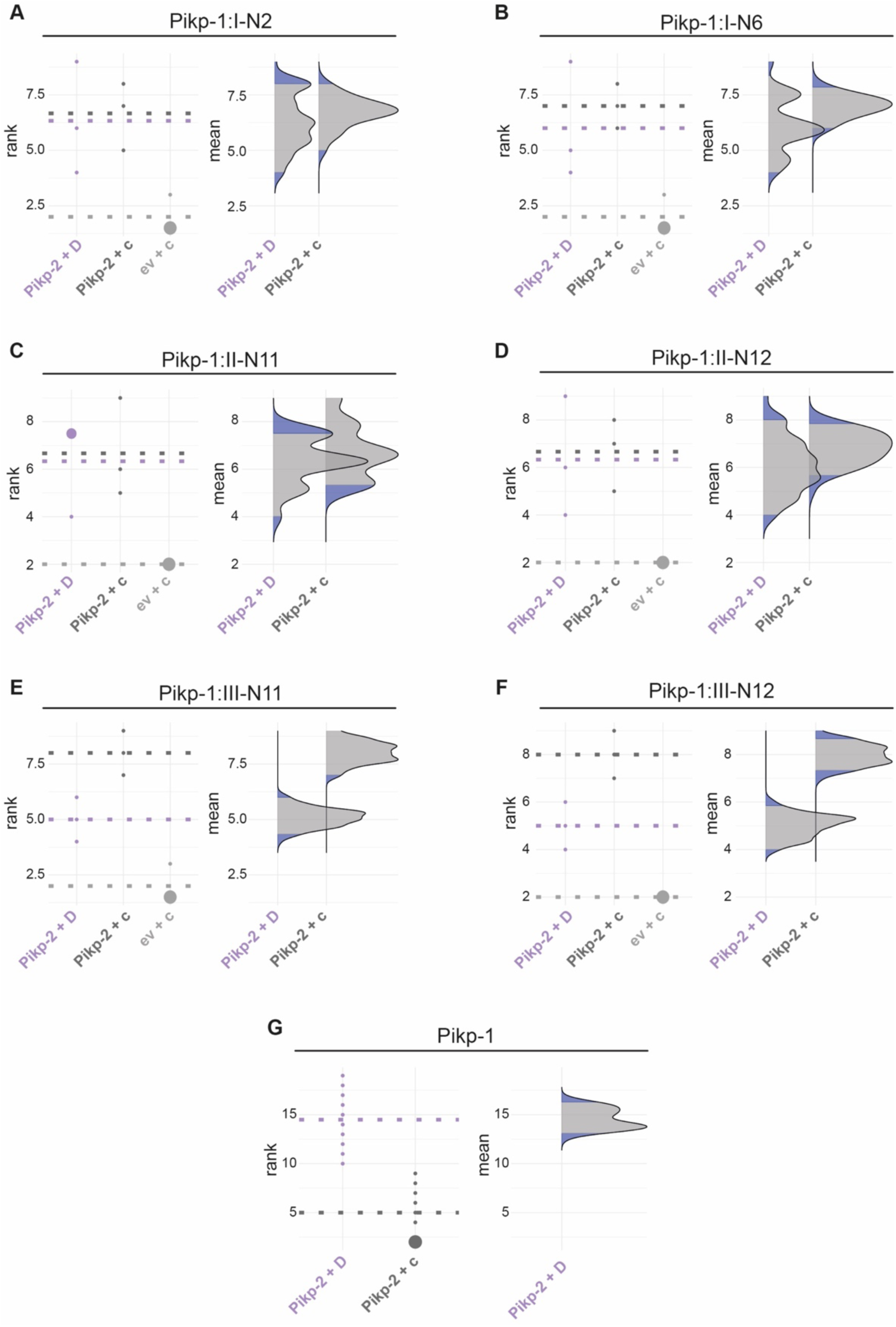
Statistical analysis of HR cell death for the Pikp-1:ancHMA fusions. The statistical analysis was conducted using an estimation method using besthr R library (MacLean, 2019). (**A–G**) Each panel corresponds to a different Pikp-1:ancHMA fusion (labelled above), co-expressed with Pikp-2 and AVR-PikD (Pikp-2 + D), Pikp-2 and AVRblb2 (Pikp-2 + c), or empty vector and AVRblb2 (ev + c). AVRblb2 and empty vector were used as controls. The left panels represent the ranked data (dots) and their corresponding mean (dashed line), with the size of a dot proportional to the number of observations with that specific value. The panels on the right show the distribution of 1000 bootstrap sample rank means, with the blue areas illustrating the 0.025 and 0.975 percentiles of the distribution. The difference is considered significant if the ranked mean for a given condition falls within or beyond the blue percentile of the mean distribution for another condition.

**Supplementary figure 22.**
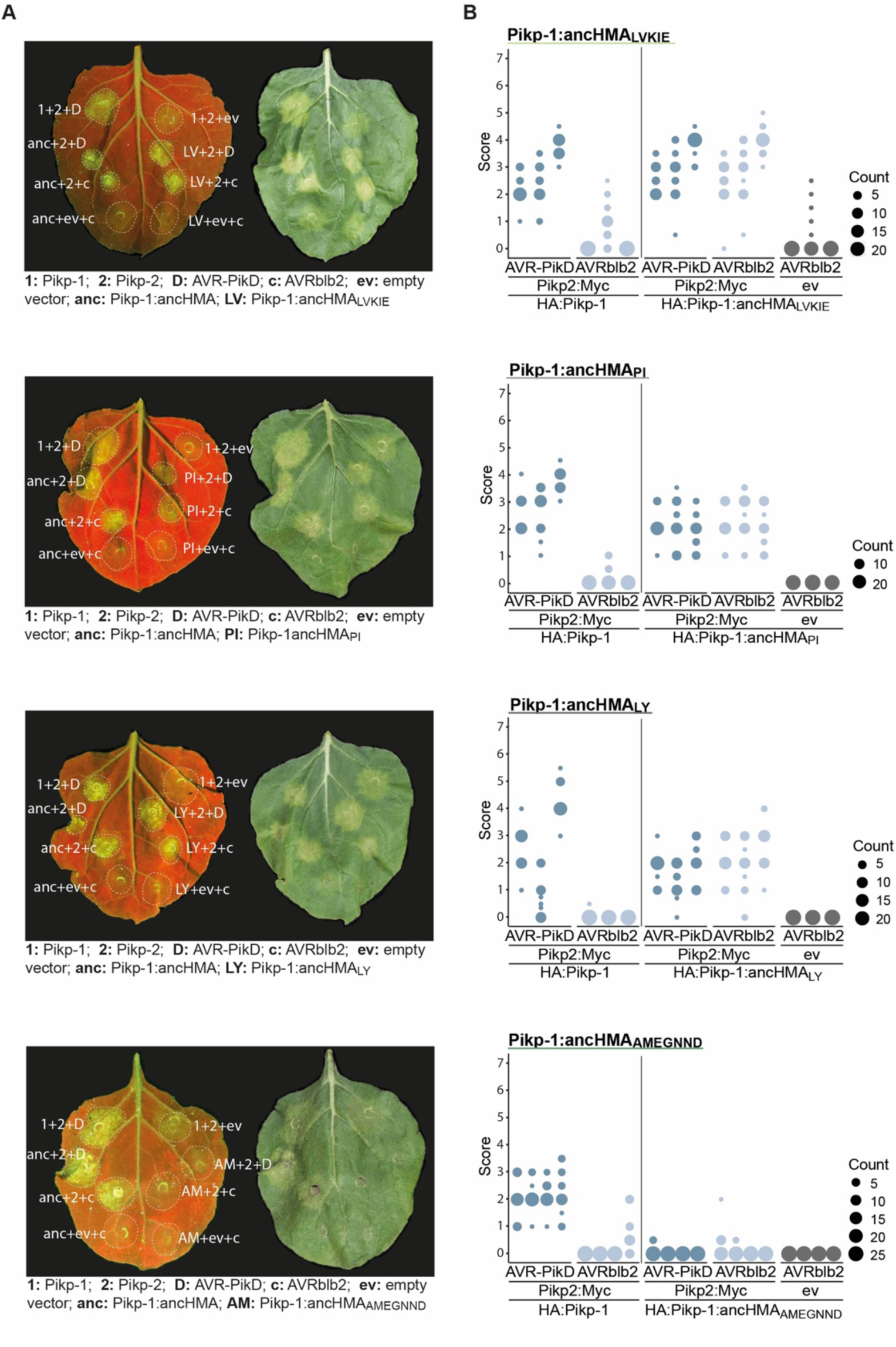
The AMEGNND mutations within ancHMA abolish autoactivatity. HR assay after transient co-expression of Pikp-1:HMA mutants (N-terminally tagged with HA) with AVR-PikD (N-terminally tagged with FLAG) and Pikp-2 (C-terminally tagged with Myc). AVRblb2 and the empty vector (ev) were used as negative controls. (**A**) Representative *N. benthamiana* leaves infiltrated with appropriate constructs (labelled next to the infiltration spot) were photographed five days post-infiltration under UV (left) and daylight (right). (**B**) HR was scored five days after agroinfiltration. The results are presented as dot plots where the size of a dot is proportional to the number of samples with the same score (count) within the same biological replicate. The experiment was independently repeated at least three times with 20–28 internal replicates; the columns within tested conditions (labelled on the bottom) illustrate results from different biological replicates. The statistical analyses of these results are presented in **Supplementary figure 23**.

**Supplementary figure 23.**
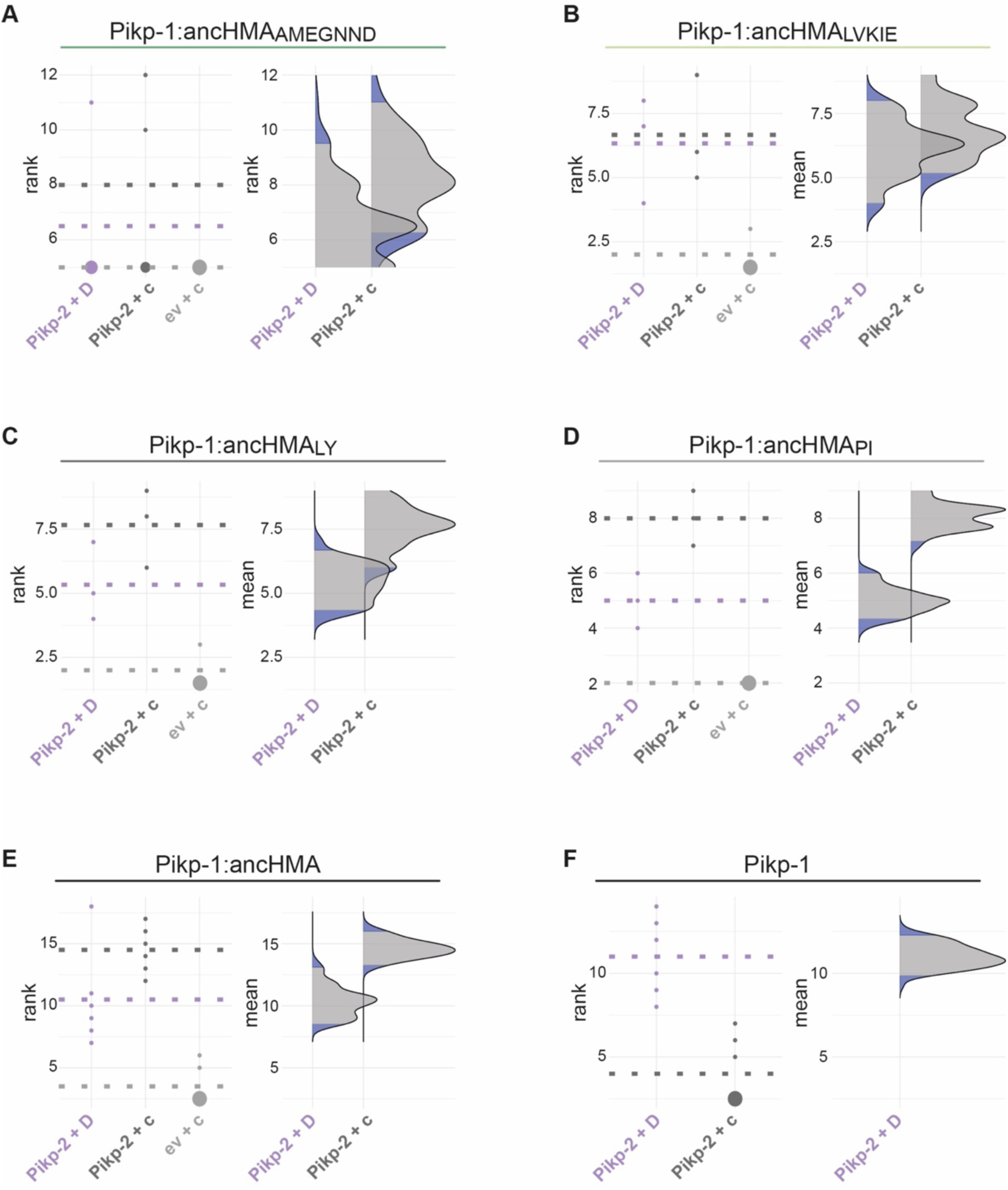
Statistical analysis of cell death assay for the Pikp-1:ancHMA chimeras. The statistical analysis was carried out using an estimation method implemented in besthr R library (MacLean, 2019). (**A–F**) Each panel corresponds to a different chimera of Pikp-1:ancHMA (labelled above), co-expressed with Pikp-2 and AVR-PikD (Pikp-2 + D), Pikp-2 and AVRblb2 (Pikp-2 + c), or empty vector and AVRblb (ev + c). AVRblb2 and empty vector were used as controls. The left panels represent the ranked data (dots) and their corresponding mean (dashed line), with the size of a dot proportional to the number of observations with that specific value. The panels on the right show the distribution of 1000 bootstrap sample rank means, with the blue areas corresponding to the 0.025 and 0.975 percentiles of the distribution. The difference is considered statistically significant if the ranked mean for a given condition falls within or beyond the blue percentile of the mean distribution for another condition.

**Supplementary figure 24.**
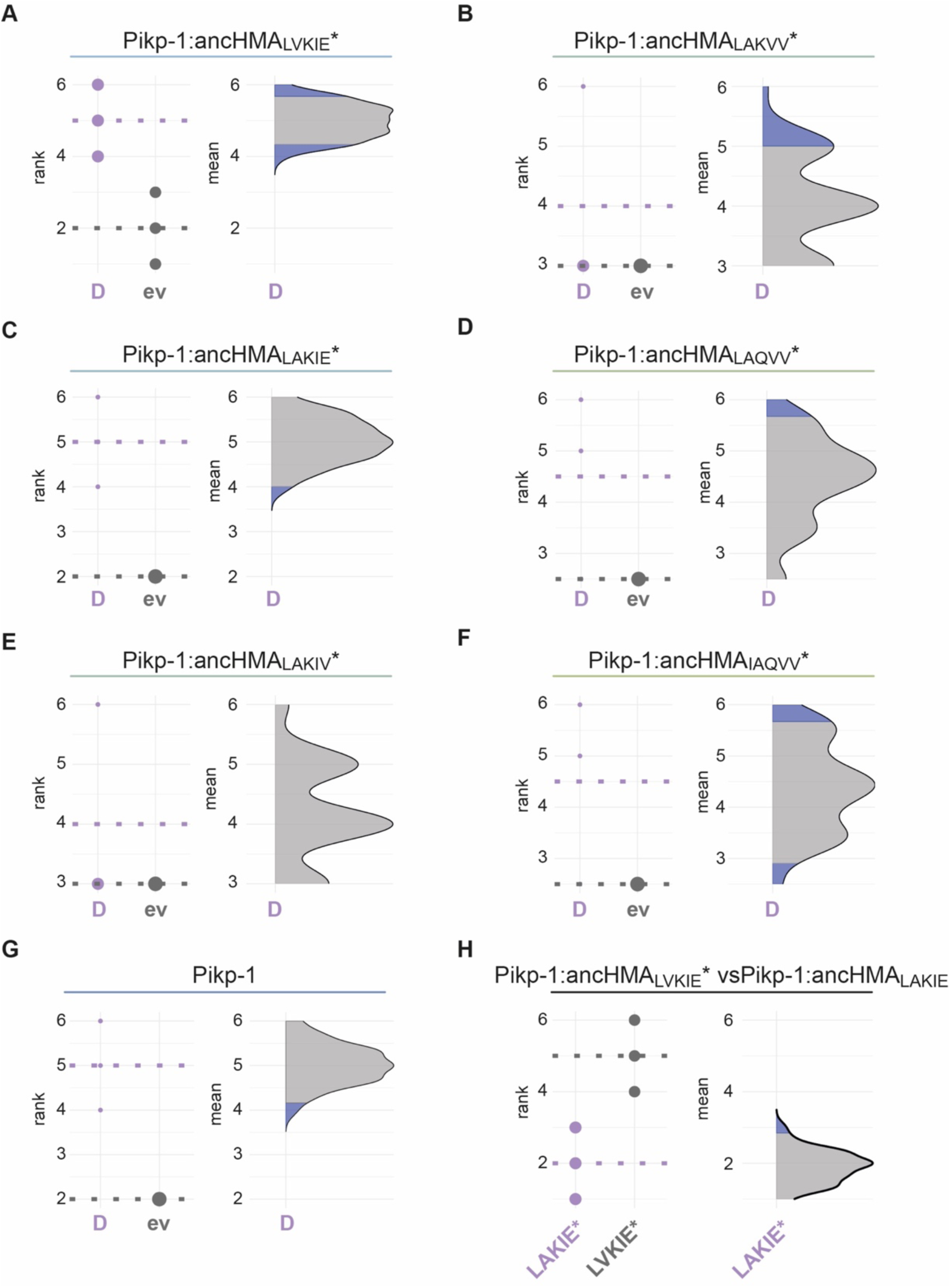
Statistical analysis of cell death for the Pikp-1:ancHMA mutants within the IAQVV/LVKIE region. The statistical analysis was performed using an estimation method implemented in besthr R library (MacLean, 2019). (**A–G**) Each panel corresponds to a different Pikp-1:ancHMA* mutant co-expressed with AVR-PikD (D), or empty vector (ev). All the constructs were co-expressed with Pikp-2. The left panels represent the ranked data (dots) and their corresponding mean (dashed line). The size of a dot centre is proportional to the number of observations with that specific value. The panels on the right show the distribution of 1000 bootstrap sample rank means, with the blue areas illustrating the 0.025 and 0.975 percentiles of the distribution. The difference is considered significant if the ranked mean for the co-expression with AVR-PikD falls within or beyond the blue percentile of the mean distribution for co-expression with the empty vector. (**H**) Statistical analysis by the estimation method of Pikp:ancHMA_LVKIE_* (LVKIE*) and Pikp:ancHMA_LAKIE_* (LAKIE*) co-expressed with AVR-PikD and Pikp-2 analysed as in panels A–G.

**Supplementary figure 25.**
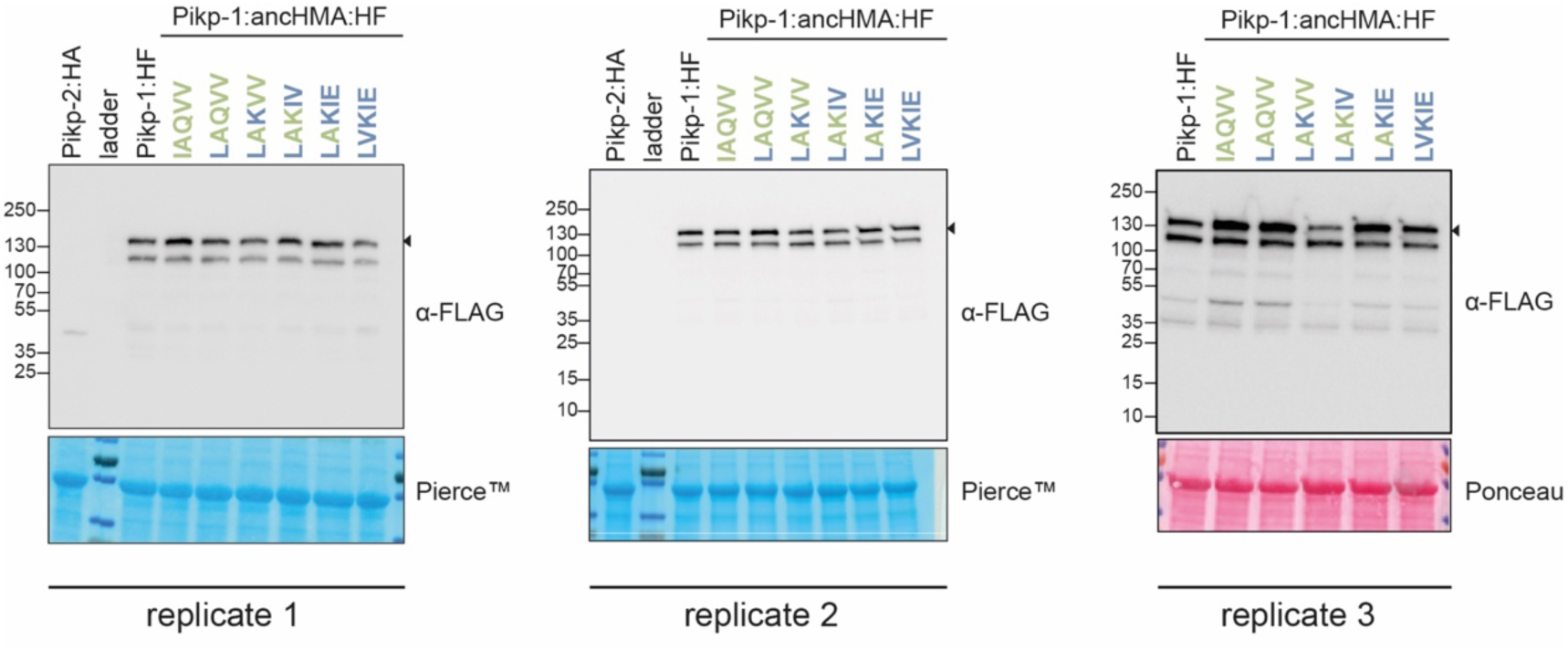
In planta accumulation of the Pikp-1:ancHMA* mutants in the IAQVV/LVKIE region. Western blot experiments of the Pikp-1:ancHMA* mutants (C-terminally tagged with HF) labelled above. Pikp-2 (C-terminally tagged with HA) was included as a negative control. Proteins were immunoblotted with the FLAG antisera (labelled on the right). Rubisco loading control was performed using Pierce^TM^ or Ponceau staining solutions. The black arrowheads indicate expected band size. The figure shows results from three independent experiments.

**Supplementary figure 26.**
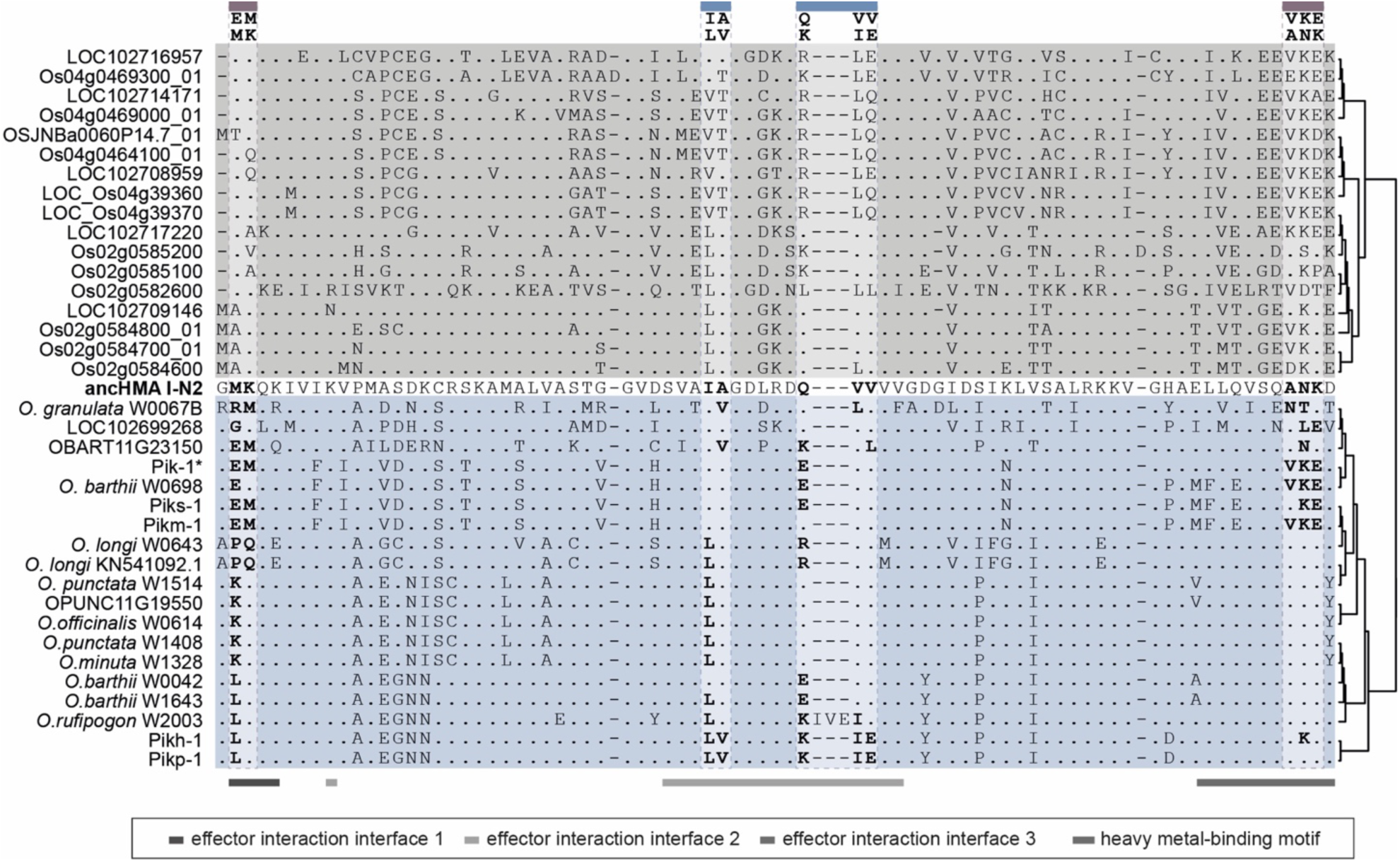
Protein sequence alignment of the HMA domain from the *Oryza* spp. Sequences of the K-type Pik-1–integrated HMA domains (blue), non-integrated HMAs from *O. sativa* and *O. brachyantha* (grey), and I-N2 ancHMA (bold) were aligned using MUSCLE (Edgar, 2004). Regions with known function were marked with horizontal lines at the bottom. The MKANK/EMVKE and IAQVV/LVKIE regions are marked above.

**Supplementary figure 27.**
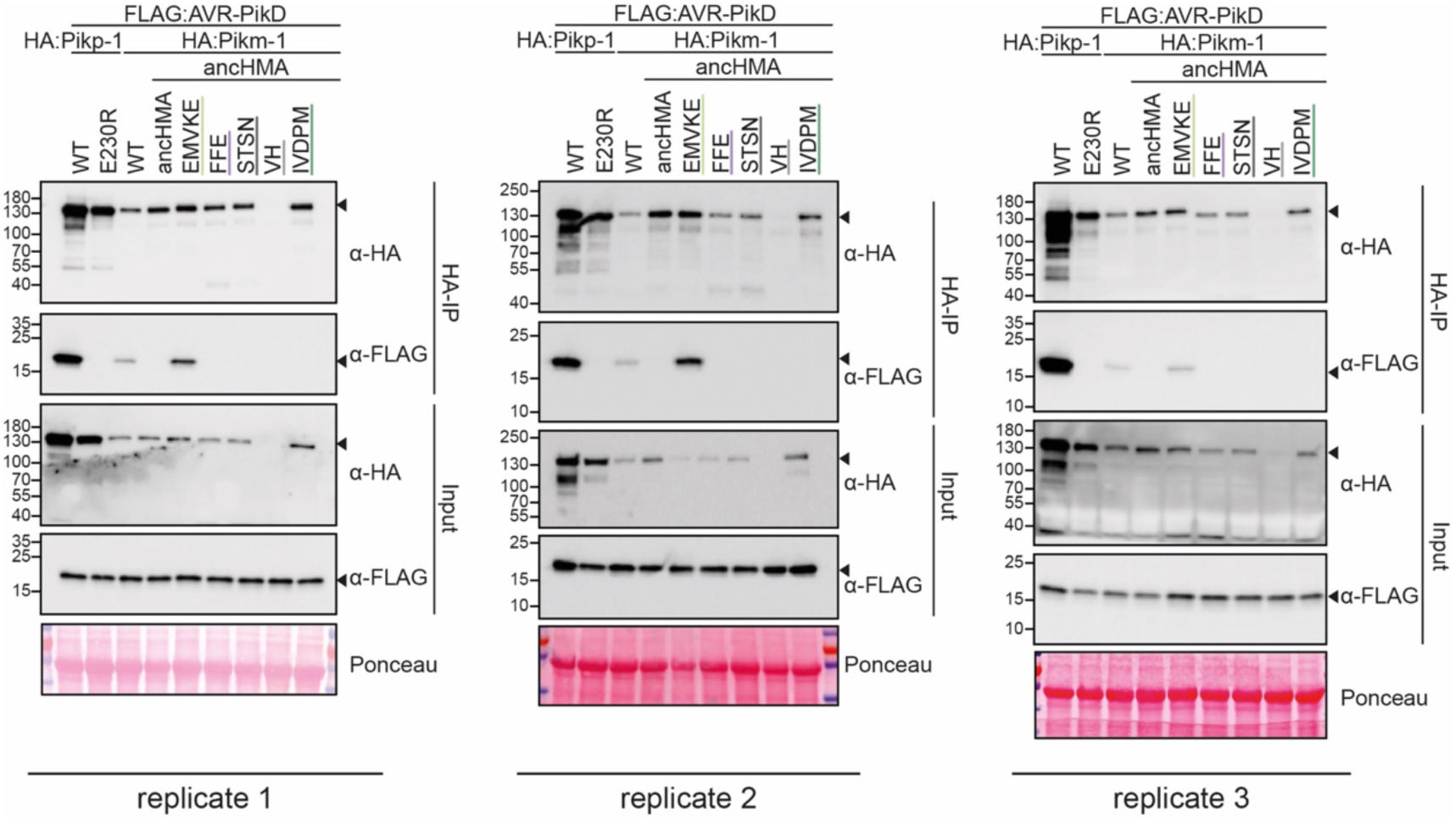
Replicates of the co-IP experiment between the Pikm-1:ancHMA chimeras and AVR-PikD. In planta association of AVR-PikD (N-terminally tagged with FLAG) with Pikp-1, Pikp-1_E230R_, Pikm-1, Pikm-1:ancHMA, and Pikm-1:ancHMA chimeras(N-terminally tagged with HA), labelled above. Wild-type (WT) Pikp-1/Pikm-1 and Pikp-1_E230R_ were used as positive and negative controls, respectively. Immunoprecipitate (HA-IP) obtained by co-IP with anti-HA probe and total protein extracts (Input) were immunoblotted with the appropriate antisera labelled on the right. Arrowheads indicate expected band sizes. Rubisco loading controls were performed using Ponceau staining. The figure presents results from three independent experiments.

**Supplementary figure 28.**
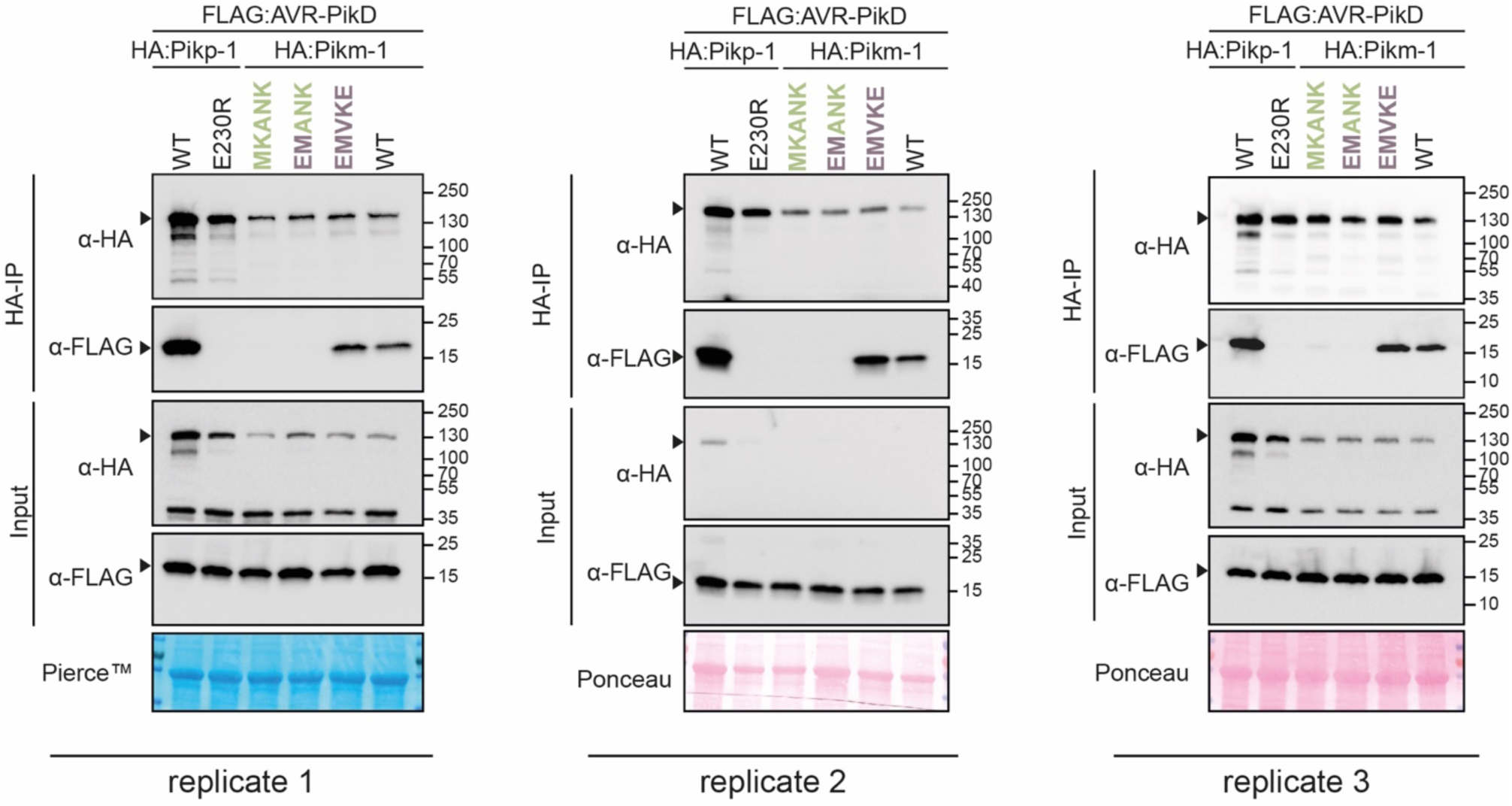
Replicates of the co-IP experiment between Pikm-1:ancHMA mutants in the MKANK/EMVKE region and AVR-PikD. In planta association of AVR-PikD (N-terminally tagged with FLAG) with Pikp-1, Pikp-1_E230R_, Pikm-1, Pikm-1:ancHMA, and Pikm-1:ancHMA mutants (N-terminally tagged with HA), labelled above. Wild-type (WT) Pikp-1/Pikm-1 and Pikp-1_E230R_ were used as positive and negative controls, respectively. Immunoprecipitates (HA-IP) obtained with anti-HA probe and total protein extracts (Input) were immunoblotted with the appropriate antisera labelled on the left. Arrowheads correspond to expected band sizes. Rubisco loading controls were performed using Pierce^TM^ or Ponceau staining. The figure depicts results from three independent experiments.

**Supplementary figure 29.**
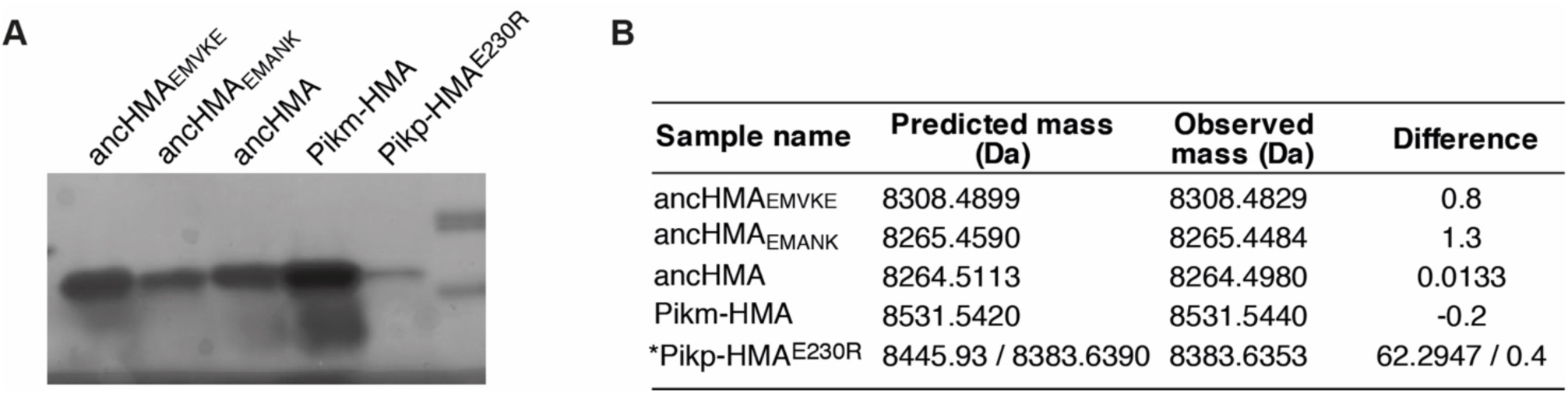
Purified proteins used in SPR studies. (**A**) Coomasie Blue–stained SDS-PAGE gel showing purified HMA proteins used in in vitro experiments. (**B**) Table summarising intact masses (monoisotopic) of proteins from panel A. (*) The Pikp-HMA^E230R^ protein appears to lack two amino acids at the N-terminus, corresponding to the linker between the protein and HIS-tag used for purification.

**Supplementary figure 30.**
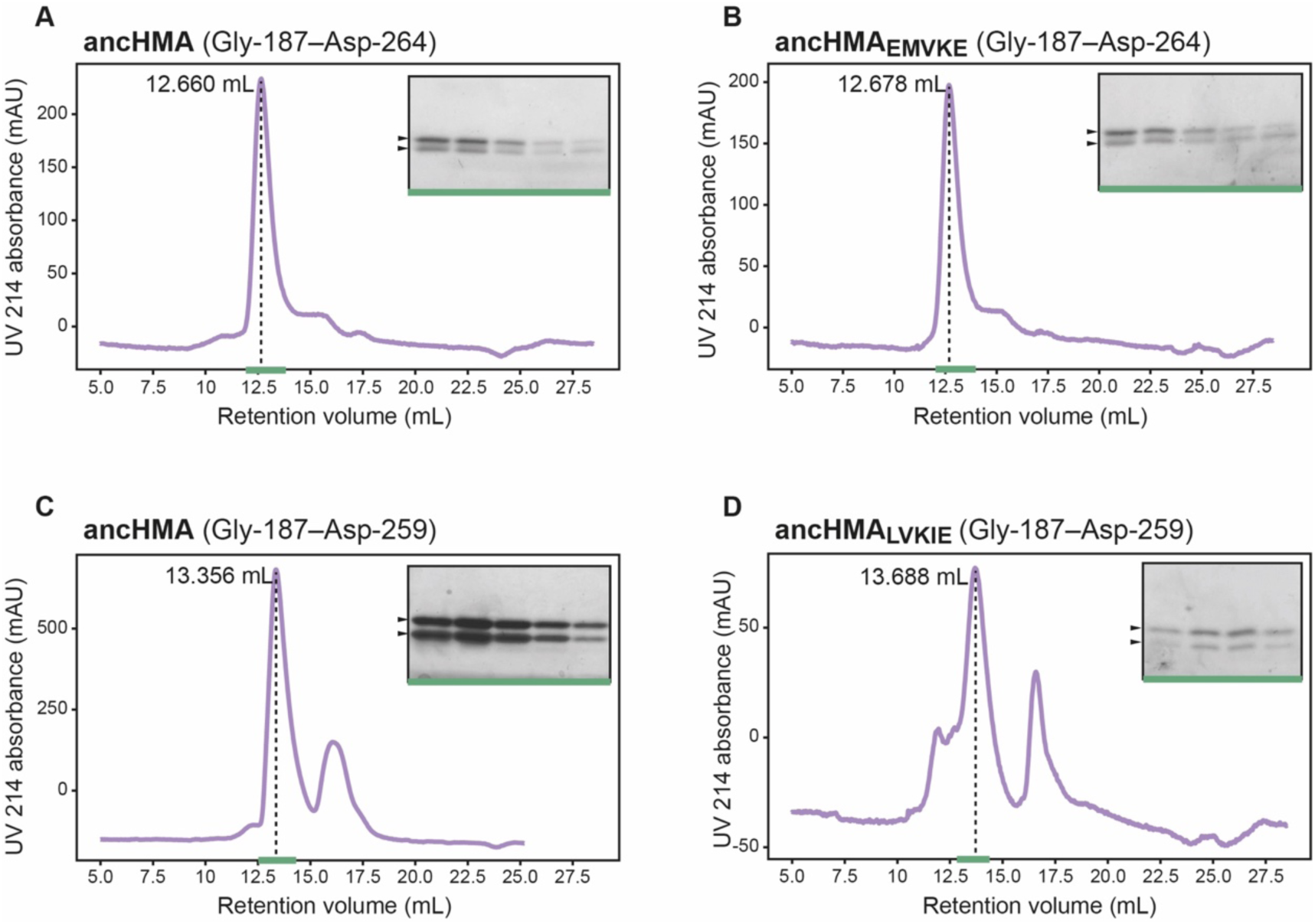
Different stoichiometry of the ancHMA–AVR-PikD complexes. Analytical gel filtration traces depicting the retention volumes of AVR-PikD in complexes with (**A**) ancHMA and (**B**) ancHMA_EMVKE_ with 5–amino acid extension, and (**C**) ancHMA and (**D**) ancHMA_LVKIE_ without the extension. The peaks corresponding to protein complexes are indicated with dashed lines, with the retention volumes shown on the left. Coomasie Blue–stained SDS-PAGE gels of relevant fractions, marked with green line, are presented of the right. Arrowheads correspond to expected protein sizes.

**Supplementary table 11.** Table of p-values for all pairwise comparisons of SPR binding to AVR-PikD between the HMA mutants.

| Concentration (nM) | Sample 1 | Sample 2 | Difference | Lower confidence level | Upper confidence level | p-value |
| --- | --- | --- | --- | --- | --- | --- |
| 400 | EMANK | E230R | 86.07761 | 62.02624 | 110.12898 | 0 |
| 400 | EMVKE | E230R | 91.209289 | 67.15792 | 115.26066 | 0 |
| 400 | MKANK | E230R | 83.519556 | 60.49213 | 106.54699 | 0 |
| 400 | Pikm | E230R | 77.298931 | 53.24757 | 101.3503 | 0 |
| 400 | EMVKE | EMANK | 5.131679 | -18.91969 | 29.18304 | 0.967442 |
| 400 | MKANK | EMANK | -2.558054 | -25.58548 | 20.46938 | 0.9971982 |
| 400 | Pikm | EMANK | -8.778679 | -32.83004 | 15.27269 | 0.8109631 |
| 400 | MKANK | EMVKE | -7.689733 | -30.71716 | 15.3377 | 0.8547095 |
| 400 | Pikm | EMVKE | -13.910358 | -37.96172 | 10.14101 | 0.4421083 |
| 400 | Pikm | MKANK | -6.220625 | -29.24805 | 16.8068 | 0.9262409 |
| 200 | EMANK | E230R | 65.82556 | 60.553591 | 71.09752 | 0 |
| 200 | EMVKE | E230R | 87.06222 | 81.790259 | 92.334189 | 0 |
| 200 | MKANK | E230R | 53.28956 | 48.017593 | 58.561522 | 0 |
| 200 | Pikm | E230R | 76.73675 | 71.464784 | 82.008713 | 0 |
| 200 | EMVKE | EMANK | 21.23667 | 15.964704 | 26.508633 | 0 |
| 200 | MKANK | EMANK | -12.536 | -17.807963 | -7.264034 | 0.0000209 |
| 200 | Pikm | EMANK | 10.91119 | 5.639228 | 16.183157 | 0.0001023 |
| 200 | MKANK | EMVKE | -33.77267 | -39.044631 | -28.500702 | 0 |
| 200 | Pikm | EMVKE | -10.32548 | -15.59744 | -5.053511 | 0.0001865 |
| 200 | Pikm | MKANK | 23.44719 | 18.175226 | 28.719156 | 0 |
| 50 | EMANK | E230R | 38.83454 | 21.734495 | 55.934593 | 0.0000044 |
| 50 | EMVKE | E230R | 76.07891 | 58.978857 | 93.178955 | 0 |
| 50 | MKANK | E230R | 27.8971 | 10.797048 | 44.997145 | 0.0005356 |
| 50 | Pikm | E230R | 63.94274 | 46.842695 | 81.042793 | 0 |
| 50 | EMVKE | EMANK | 37.24436 | 19.498803 | 54.989921 | 0.0000157 |
| 50 | MKANK | EMANK | -10.93745 | -28.683007 | 6.808111 | 0.3922477 |
| 50 | Pikm | EMANK | 25.1082 | 7.362641 | 42.853759 | 0.0027261 |
| 50 | MKANK | EMVKE | -48.18181 | -65.927369 | -30.436251 | 0.0000002 |
| 50 | Pikm | EMVKE | -12.13616 | -29.881721 | 5.609397 | 0.2926768 |
| 50 | Pikm | MKANK | 36.04565 | 18.300089 | 53.791207 | 0.000026 |

E230R: Pikp-HMA<sub>E230R</sub>, EMVKE: anchMA<sub>EMVKE</sub>, EMANK: anchMA<sub>EMANK</sub>, MKANK: anchMA, Pikm: Pikm-HMA

**Supplementary figure 31.**
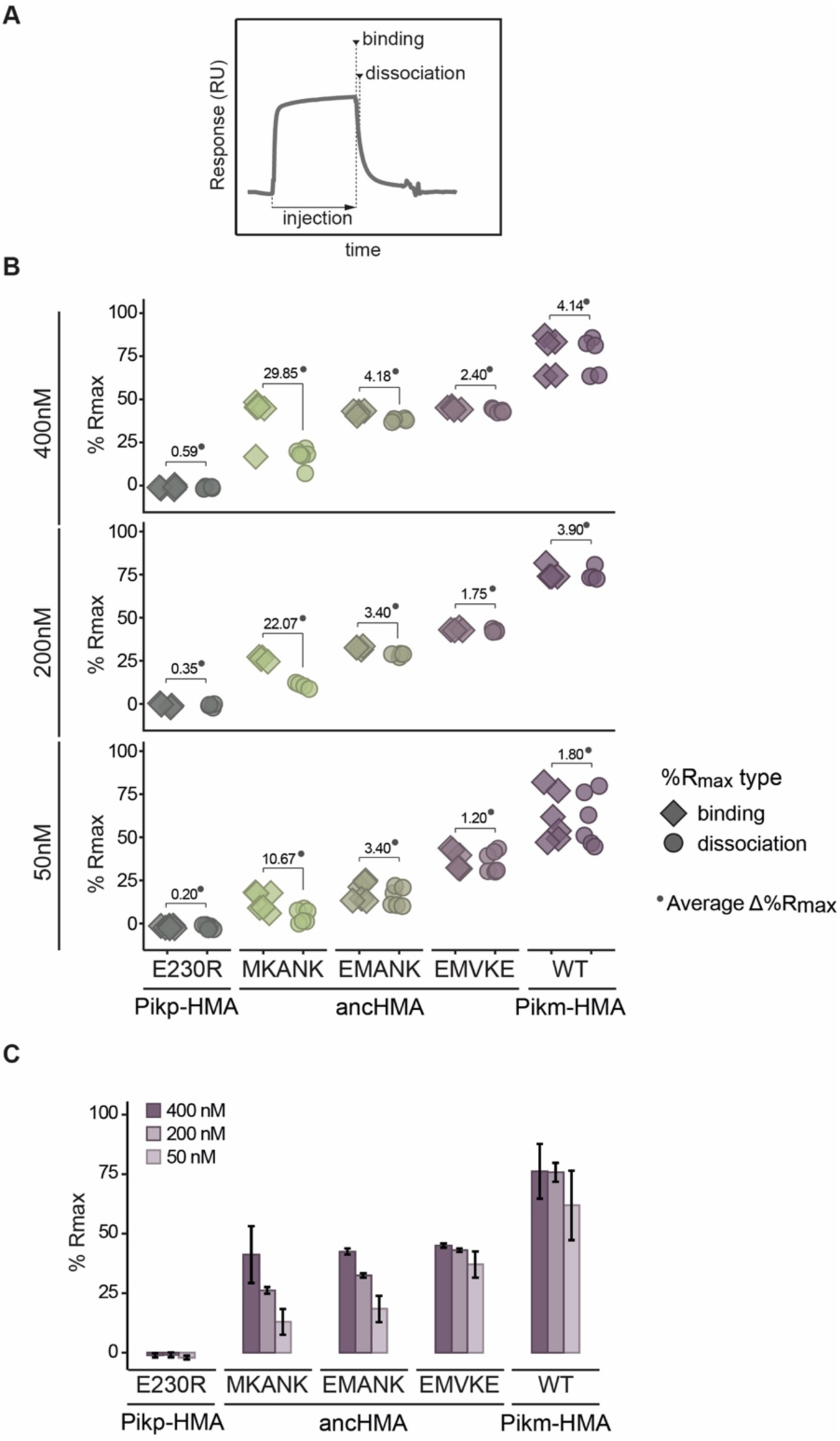
SPR results showing the effect of the step-by-step mutations within the MKANK/EMVKE region on the AVR-PikD binding in vitro, as indicated by %R_max_. (**A**) Schematic illustration of the SPR sensorgram and the timepoints corresponding to ‘binding’ and ‘dissociation’, recorded in this study. (**B**) Plots illustrating calculated percentage of the theoretical maximum response (%R_max_) values for interaction of HMA analytes, labelled below, with AVR-PikD ligand (C-terminally tagged with HIS). %R_max_ was calculated assuming a two-to-one model for Pikp-HMA_E230R_ and a one-to-one binding model for the remaining constructs. The values were normalized for the amount of ligand immobilized on the NTA-chip. The HMA analytes were tested at three different concentrations (labelled on the left) in at least four independent experiments. All of the data points are represented as diamonds or circles. Average Δ%R_max_ (•) values represent absolute differences between values for ‘binding’ and ‘‘dissociation’, calculated from average values for each sample, and serve as an off-rate approximate. Statistical differences among the samples were analysed with ANOVA and Tukey’s honest significant difference (HSD) test (p < 0.01). P-values for all pairwise comparisons are presented in **Supplementary table 11**. (**C**) The results, identical to those presented in panel B, are shown as histograms to emphasise the differences in binding dynamics between the constructs. Bars represent the average response, and the error bars represent the standard deviation.

**Supplementary figure 32.**
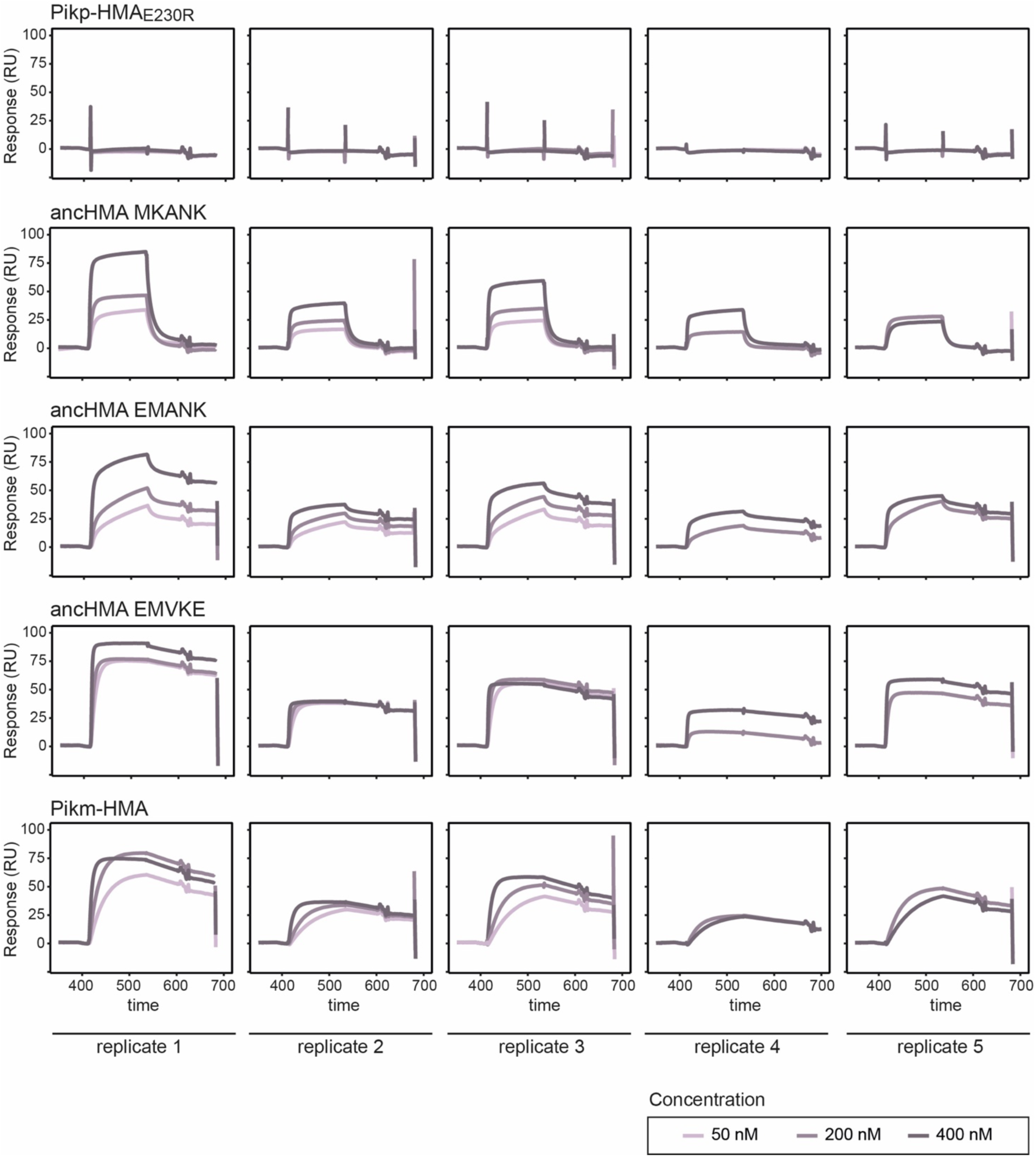
The SPR sensorgrams for the AVR-PikD–HMA binding. The SPR sensorgrams from five independent replicates are shown. His-tagged AVR-PikD was immobilised on the sample cell, giving a response level of 99 ± 33 response units (RU).

**Supplementary table 12.**
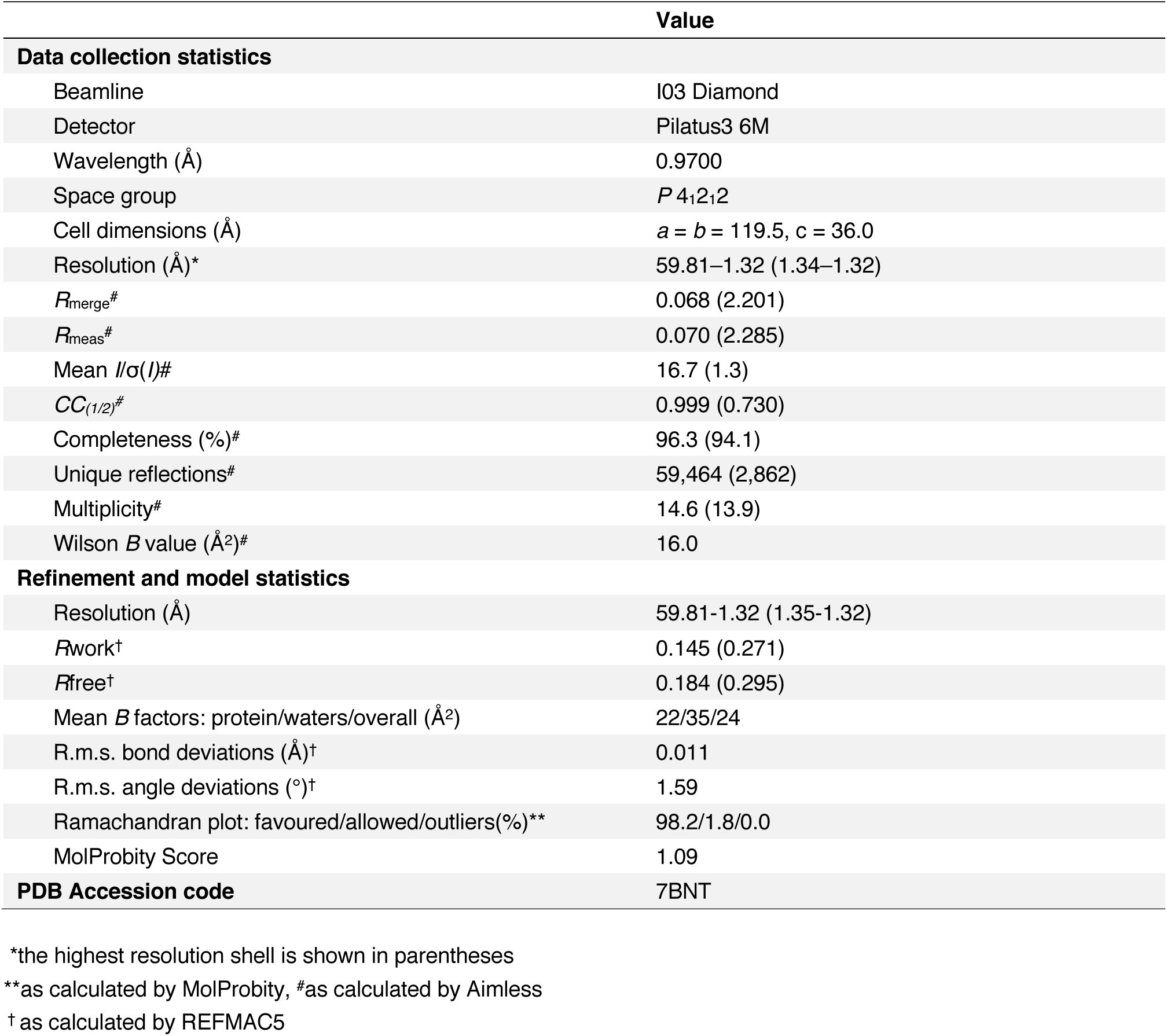
Data collection and refinement statistics for the ancHMA_LVKIE_–AVR-PikD co-crystal structure.

|  | Value |
| --- | --- |
| **Data collection statistics** |  |
| Beamline | I03 Diamond |
| Detector | Pilatus3 6M |
| Wavelength (Å) | 0.9700 |
| Space group | *P* 4<sub>1</sub>2<sub>1</sub>2 |
| Cell dimensions (Å) | *a* = *b* = 119.5, *c* = 36.0 |
| Resolution (Å)* | 59.81–1.32 (1.34–1.32) |
| *R*<sub>merge</sub><sup>#</sup> | 0.068 (2.201) |
| *R*<sub>meas</sub><sup>#</sup> | 0.070 (2.285) |
| Mean *I*/σ(*I*)<sup>#</sup> | 16.7 (1.3) |
| *CC*<sub>(1/2)</sub><sup>#</sup> | 0.999 (0.730) |
| Completeness (%)<sup>#</sup> | 96.3 (94.1) |
| Unique reflections<sup>#</sup> | 59,464 (2,862) |
| Multiplicity<sup>#</sup> | 14.6 (13.9) |
| Wilson *B* value (Å<sup>2</sup>)<sup>#</sup> | 16.0 |
| **Refinement and model statistics** |  |
| Resolution (Å) | 59.81–1.32 (1.35–1.32) |
| *R*<sub>work</sub><sup>†</sup> | 0.145 (0.271) |
| *R*<sub>free</sub><sup>†</sup> | 0.184 (0.295) |
| Mean *B* factors: protein/waters/overall (Å<sup>2</sup>) | 22/35/24 |
| R.m.s. bond deviations (Å)<sup>†</sup> | 0.011 |
| R.m.s. angle deviations (°)<sup>†</sup> | 1.59 |
| Ramachandran plot: favoured/allowed/outliers(%)<sup>**</sup> | 98.2/1.8/0.0 |
| MolProbity Score | 1.09 |
| **PDB Accession code** | 7BNT |

\*the highest resolution shell is shown in parentheses
\*\*as calculated by MolProbity, #as calculated by Aimless
† as calculated by REFMAC5

**Supplementary figure 33.**
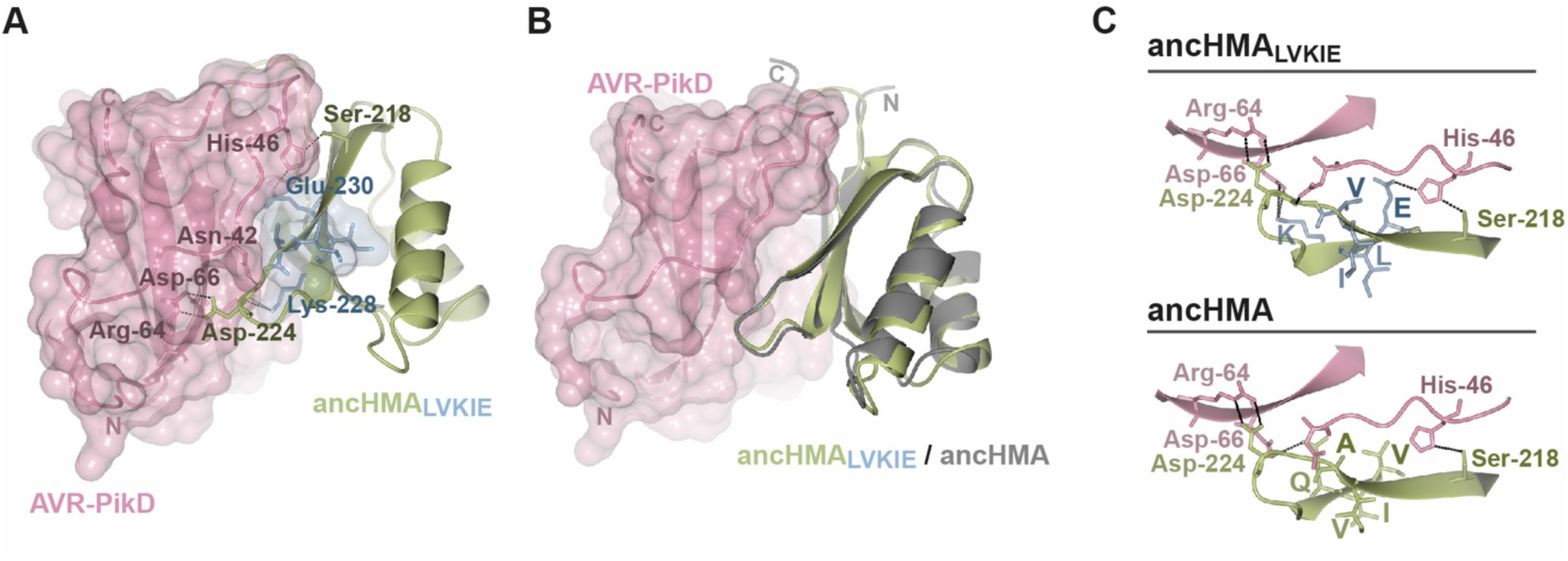
The Val-230-Glu mutation within the LVKIE region of ancHMA enhances interaction with AVR-PikD through hydrogen bond formation. (**A**) Schematic representation of the structure of ancHMA_LVKIE_ complexed with the AVR-PikD effector. The molecules are shown as ribbons with selected side chains presented as sticks and labelled; the colours of the residue labels match colours of the respective molecules. The molecular surfaces of AVR-PikD (pink) and LVKIE residues (blue) within ancHMA_LVKIE_ are also shown. Dashed lines represent hydrogen bonds or salt bridges formed between the two molecules. (**B**) Superimposition of the ancHMA_LVKIE_ structure and the ancHMA homology model (grey) in complex with AVR-PikD. (**C**) Close-up views of the IAQVV/LVKIE region of ancHMA_LVKIE_ (green and blue) and ancHMA (green) showcasing differences in binding to AVR-PikD (pink). The selected residues involved in binding are labelled with labels matching the colours of the corresponding molecules. The LVKIE residues are labelled with single-letter amino acid symbols—Ile/Leu-221 (I/L), Ala/Val-222 (A/V), Gln/Lys-228 (Q/K), Val/Ile-229 (V/I), Val/Glu-230 (V/E) for ancHMA/ancHMA_LVKIE_; and Ile-222 (I), Ala-223 (A), Gln-231 (Q), Val-232 (V), Val-233 (V) for Pikm-HMA. Dashed lines represent hydrogen bonds or salt bridges formed between the two molecules.

**Supplementary table 13.** HMA sequences used for building phylogenetic trees and ancestral sequence reconstruction (ASR).

| Description on the tree | Species | Accession number | Pik-integrated | used for ASR |  |  |
| --- | --- | --- | --- | --- | --- | --- |
|  |  |  |  | I | II | III |
| ***O.barthii_W0042*** | *O. barthii* | PCR from NIG accession no. W0042 | y | n | n | y |
| ***O.barthii_W1643*** | *O. barthii* | PCR from NIG accession no. W1643 | y | n | n | y |
| ***O.punctata_W1408*** | *O. punctata* | PCR from NIG accession no. W1408 | y | n | n | y |
| ***O.barthii_W0698*** | *O. barthii* | PCR from NIG accession no. W0698 | y | n | n | y |
| ***O.granulata_W0067B*** | *O. granulata* | PCR from NIG accession no. W0067(B) | y | n | n | y |
| ***O.longistaminata_W0643*** | *O. longistaminata* | PCR from NIG accession no. W0643 | y | n | n | y |
| ***O.officinalis_W0614*** | *O. officinalis* | PCR from NIG accession no. W0614 | y | n | n | y |
| ***O.punctata_W1514*** | *O. punctata* | PCR from NIG accession no. W1514 | y | n | n | y |
| ***O.rufipogon_W2003*** | *O. rufipogon* | PCR from NIG accession no. W2003 | y | n | n | y |
| ***O.minuta_W1328*** | *O. minuta* | PCR from NIG accession no. W1328 | y | n | n | y |
| **LOC102699268** | *O. brachyantha* | LOC102699268 | y | y | y | y |
| **OBART11G23150** | *O. barthii* | OBART11G23150 | y | n | n | y |
| **Olongi_KN541092.1** | *O. longistaminata* | KN541092.1 | y | n | n | y |
| **OPUNC11G19550** | *O. punctata* | OPUNC11G19550 | y | n | n | y |
| **OsPikp-1** | *O. sativa* | HM035360.1 | y | y | y | y |
| **OsPik-1** | *O. sativa* | HM048900_1 | y | y | y | Y |
| **OsPikh-1** | *O. sativa* | HQ662330_1 | y | y | y | Y |
| **OsPiks-1** | *O. sativa* | HQ662329_1 | y | y | y | y |
| **OsPikm-1** | *O. sativa* | AB462324.1 | y | y | y | y |
| **Ob_LOC102708959** | *O. brachyantha* | LOC102708959 | n | n | y | y |
| **Ob_LOC102709146** | *O. brachyantha* | LOC102709146 | n | n | y | y |
| **Ob_LOC102714171** | *O. brachyantha* | LOC102714171 | n | n | y | y |
| **Ob_LOC102716957** | *O. brachyantha* | LOC102716957 | n | n | y | y |
| **Ob_LOC102717220** | *O. brachyantha* | LOC102717220 | n | n | y | y |
| **Os_LOC_Os04g39360** | *O. sativa* | LOC_Os04g39360 | n | y | y | y |
| **Os_LOC_Os04g39370** | *O. sativa* | LOC_Os04g39370 | n | y | y | y |
| **Os04g0469000_01** | *O. sativa* | Os04g0469000_01 | n | y | y | y |
| **Os02g0585200** | *O. sativa* | Os02g0585200 | n | y | y | y |
| **Os02g0584800_01** | *O. sativa* | Os02g0584800_01 | n | y | y | y |
| **Os02g0584700_01** | *O. sativa* | Os02g0584700_01 | n | y | y | y |
| **Os04g0469300_01** | *O. sativa* | Os04g0469300_01 | n | y | y | y |
| **Os02g0585100** | *O. sativa* | Os02g0585100 | n | y | y | y |
| **Os02g0584600** | *O. sativa* | Os02g0584600 | n | y | y | y |
| **OSJNBa0060P14.7_01** | *O. sativa* | OSJNBa0060P14.7_01 | n | y | y | y |
| **Os04g0464100_01** | *O. sativa* | Os04g0464100_01 | n | y | y | y |
| **Os02g0582600** | *O. sativa* | Os02g0582600 | n | y | y | y |

**Supplementary table 14.**
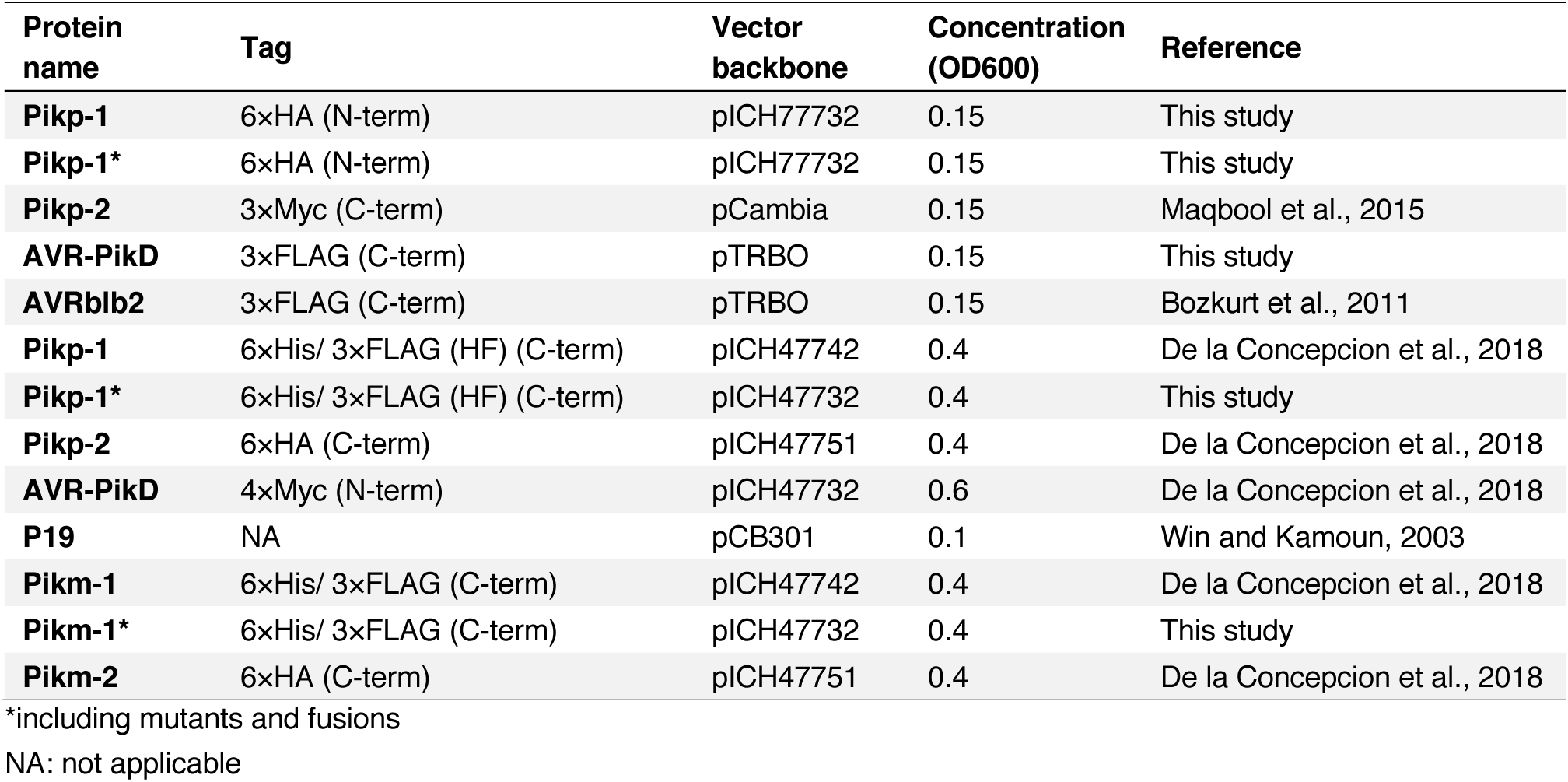
List of constructs used in cell death assays.

| Protein name | Tag | Vector backbone | Concentration (OD600) | Reference |
| --- | --- | --- | --- | --- |
| **Pikp-1** | 6xHA (N-term) | pICH77732 | 0.15 | This study |
| **Pikp-1\*** | 6xHA (N-term) | pICH77732 | 0.15 | This study |
| **Pikp-2** | 3xMyc (C-term) | pCambia | 0.15 | Maqbool et al., 2015 |
| **AVR-PikD** | 3xFLAG (C-term) | pTRBO | 0.15 | This study |
| **AVRblb2** | 3xFLAG (C-term) | pTRBO | 0.15 | Bozkurt et al., 2011 |
| **Pikp-1** | 6xHis/ 3xFLAG (HF) (C-term) | pICH47742 | 0.4 | De la Concepcion et al., 2018 |
| **Pikp-1\*** | 6xHis/ 3xFLAG (HF) (C-term) | pICH47732 | 0.4 | This study |
| **Pikp-2** | 6xHA (C-term) | pICH47751 | 0.4 | De la Concepcion et al., 2018 |
| **AVR-PikD** | 4xMyc (N-term) | pICH47732 | 0.6 | De la Concepcion et al., 2018 |
| **P19** | NA | pCB301 | 0.1 | Win and Kamoun, 2003 |
| **Pikm-1** | 6xHis/ 3xFLAG (C-term) | pICH47742 | 0.4 | De la Concepcion et al., 2018 |
| **Pikm-1\*** | 6xHis/ 3xFLAG (C-term) | pICH47732 | 0.4 | This study |
| **Pikm-2** | 6xHA (C-term) | pICH47751 | 0.4 | De la Concepcion et al., 2018 |

\*including mutants and fusions
NA: not applicable

## SUPPLEMENTARY FILES

**Supplementary File 1** Full list of all Poaceae NLRs and filtering details.xlsx

**Supplementary File 2** Site selection test for K-type HMAs.xlsx

**Supplementary File 3** ancHMA prediction probabilities.xlsx

## SOURCE DATA FILES

**Source data 1** Selection test for Pik-1 vs Pik-2 orthologues.xlsx

**Source data 2** Selection test for Pik-1-HMA vs NB-ARC.xlsx

**Source data 3** Raw data of Pikp-ancHMA Rmax SPR.xlsx

**Source data 4** HR scores used in SFig20.xlsx

**Source data 5** HR scores used in SFig22.xlsx

**Source data 6** HR scores for IAQVV to LVKIE mutations in Pikp-HMA.xlsx

**Source data 7** Raw data of Pikm-ancHMA Rmax SPR.xlsx

## REFERENCES

Adachi H, Derevnina L, Kamoun S. 2019. NLR singletons, pairs, and networks: evolution, assembly, and regulation of the intracellular immunoreceptor circuitry of plants. Curr Opin Plant Biol 50:121–131. doi:10.1016/j.pbi.2019.04.007

Alcázar R, García A V., Parker JE, Reymond M. 2009. Incremental steps toward incompatibility revealed by Arabidopsis epistatic interactions modulating salicylic acid pathway activation. Proc Natl Acad Sci U S A 106:334– 339. doi:10.1073/pnas.0811734106

Altschul SF, Gish W, Miller W, Myers EW, Lipman DJ. 1990. Basic local alignment search tool. J Mol Biol 215:403–410. doi:10.1016/S0022-2836(05)80360-2

Ashikawa I, Hayashi N, Yamane H, Kanamori H, Wu J, Matsumoto T, Ono K, Yano M. 2008. Two adjacent nucleotide-binding site-leucine-rich repeat class genes are required to confer Pikm-specific rice blast resistance. Genetics 180:2267–2276. doi:10.1534/genetics.108.095034

Ashkenazy H, Penn O, Doron-Faigenboim A, Cohen O, Cannarozzi G, Zomer O, Pupko T. 2012. FastML: A web server for probabilistic reconstruction of ancestral sequences. Nucleic Acids Res 40:580–584. doi:10.1093/nar/gks498

Baggs E, Dagdas G, Krasileva K V. 2017. NLR diversity, helpers and integrated domains: making sense of the NLR IDentity. Curr Opin Plant Biol 38:59–67. doi:10.1016/j.pbi.2017.04.012

Bailey PC, Schudoma C, Jackson W, Baggs E, Dagdas G, Haerty W, Moscou M, Krasileva K V. 2018. Dominant integration locus drives continuous diversification of plant immune receptors with exogenous doFusions. Genome Biol 19:23. doi:10.1186/s13059-018-1392-6

Bao J, Chen M, Zhong Z, Tang W, Lin L, Zhang X, Jiang H, Zhang D, Miao C, Tang H, Zhang J, Lu G, Ming R, Norvienyeku J, Wang B, Wang Z. 2017. PacBio sequencing reveals transposable elements as a key contributor to genomic plasticity and virulence variation in *Magnaporthe oryzae*. Mol Plant 1465–1468. doi:10.1016/j.molp.2017.08.008

Berrow NS, Alderton D, Sainsbury S, Nettleship J, Assenberg R, Rahman N, Stuart DI, Owens RJ. 2007. A versatile ligation-independent cloning method suitable for high-throughput expression screening applications. Nucleic Acids Res 35:e45–e45. doi:10.1093/nar/gkm047

Bhullar NK, Zhang Z, Wicker T, Keller B. 2010. Wheat gene bank accessions as a source of new alleles of the powdery mildew resistance gene *Pm3*: A large scale allele mining project. BMC Plant Biol 10:88. doi:10.1186/1471-2229-10-88

Białas A, Zess EK, De la Concepcion JC, Franceschetti M, Pennington HG, Yoshida K, Upson JL, Chanclud E, Wu C-H, Langner T, Maqbool A, Varden FA, Derevnina L, Belhaj K, Fujisaki K, Saitoh H, Terauchi R, Banfield MJ, Kamoun S. 2018. Lessons in effector and NLR biology of plant-microbe systems. Mol Plant-Microbe Interact 31:34–45. doi:10.1094/MPMI-08-17-0196-FI

Bomblies K, Lempe J, Epple P, Warthmann N, Lanz C, Dangl JL, Weigel D. 2007. Autoimmune response as a mechanism for a Dobzhansky-Muller-type incompatibility syndrome in plants. PLoS Biol 5:e236. doi:10.1371/journal.pbio.0050236

Bonardi V, Tang SJ, Stallmann A, Roberts M, Cherkis K, Dangl JL. 2011. Expanded functions for a family of plant intracellular immune receptors beyond specific recognition of pathogen effectors. Proc Natl Acad Sci U S A 108:16463–16468. doi:10.1073/pnas.1113726108

Bos JIB, Kanneganti T-D, Young C, Cakir C, Huitema E, Win J, Armstrong MR, Birch PRJ, Kamoun S. 2006. The C-terminal half of *Phytophthora infestans* RXLR effector AVR3a is sufficient to trigger R3a-mediated hypersensitivity and suppress INF1-induced cell death in *Nicotiana benthamiana*. Plant J 48:165–76. doi:10.1111/j.1365-313X.2006.02866.x

Bozkurt TO, Schornack S, Win J, Shindo T, Ilyas M, Oliva R, Cano LM, Jones AME, Huitema E, van der Hoorn R a L, Kamoun S. 2011. *Phytophthora infestans* effector AVRblb2 prevents secretion of a plant immune protease at the haustorial interface. Proc Natl Acad Sci U S A 108:20832–20837. doi:10.1073/pnas.1112708109

Brueggeman R, Druka A, Nirmala J, Cavileer T, Drader T, Rostoks N, Mirlohi A, Bennypaul H, Gill U, Kudrna D, Whitelaw C, Kilian A, Han F, Sun Y, Gill K, Steffenson B, Kleinhofs A. 2008. The stem rust resistance gene *Rpg5* encodes a protein with nucleotide-binding-site, leucine-rich, and protein kinase domains. Proc Natl Acad Sci U S A 105:14970–14975. doi:10.1073/pnas.0807270105

Bryan GT, Wu K-S, Farrall L, Jia Y, Hershey HP, McAdams SA, Faulk KN, Donaldson GK, Tarchini R, Valent B. 2000. A single amino acid difference distinguishes resistant and susceptible alleles of the rice blast resistance gene *Pi-ta*. Plant Cell 12:2033–2045. doi:10.1105/tpc.12.11.2033

Bulgarelli D, Biselli C, Collins NC, Consonni G, Stanca AM, Schulze-Lefert P, Valè G. 2010. The CC-NB-LRR-type *Rdg2a* resistance gene confers immunity to the seed-borne barley leaf stripe pathogen in the absence of hypersensitive cell death. PLoS One 5:e12599. doi:10.1371/journal.pone.0012599

Cesari S. 2018. Multiple strategies for pathogen perception by plant immune receptors. New Phytol 219:17–24. doi:10.1111/nph.14877

Cesari S, Bernoux M, Moncuquet P, Kroj T, Dodds PN. 2014a. A novel conserved mechanism for plant NLR protein pairs: the “integrated decoy” hypothesis. Front Plant Sci 5:606. doi:10.3389/fpls.2014.00606

Cesari S, Kanzaki H, Fujiwara T, Bernoux M, Chalvon V, Kawano Y, Shimamoto K, Dodds P, Terauchi R, Kroj T. 2014b. The NB-LRR proteins RGA4 and RGA5 interact functionally and physically to confer disease resistance. EMBO J 33:1941–1959. doi:10.15252/embj.201487923

Cesari S, Thilliez G, Ribot C, Chalvon V, Michel C, Jauneau A, Rivas S, Alaux L, Kanzaki H, Okuyama Y, Morel J-BJ-B, Fournier E, Tharreau D, Terauchi R, Kroj T. 2013. The rice resistance protein pair RGA4/RGA5 recognizes the *Magnaporthe oryzae* effectors AVR-Pia and AVR1-CO39 by direct binding. Plant Cell 25:1463–1481. doi:10.1105/tpc.112.107201

Chae E, Tran DTN, Weigel D. 2016. Cooperation and conflict in the plant immune system. PLOS Pathog 12:e1005452. doi:10.1371/journal.ppat.1005452

Chen J, Huang Q, Gao D, Wang J, Lang Y, Liu T, Li B, Bai Z, Luis Goicoechea J, Liang C, Chen C, Zhang W, Sun S, Liao Y, Zhang X, Yang L, Song C, Wang M, Shi J, Liu G, Liu J, Zhou H, Zhou W, Yu Q, An N, Chen Y, Cai Q, Wang B, Liu B, Min J, Huang Y, Wu H, Li Z, Zhang Y, Yin Y, Song W, Jiang J, Jackson S a, Wing R a, Wang J, Chen M. 2013. Whole-genome sequencing of Oryza *brachyantha* reveals mechanisms underlying *Oryza* genome evolution. Nat Commun 4:1595. doi:10.1038/ncomms2596

Chen VB, Arendall WB, Headd JJ, Keedy DA, Immormino RM, Kapral GJ, Murray LW, Richardson JS, Richardson DC. 2010. MolProbity: all-atom structure validation for macromolecular crystallography. Acta Crystallogr Sect D Biol Crystallogr 66:12–21. doi:10.1107/S0907444909042073

Cohen O, Pupko T. 2011. Inference of gain and loss events from phyletic patterns using stochastic mapping and maximum parsimony-A simulation study. Genome Biol Evol 3:1265–1275. doi:10.1093/gbe/evr101

Collins N, Drake J, Ayliffe M, Sun Q, Ellis J, Hulbert S, Pryor T. 1999. Molecular characterization of the maize Rp1-D rust resistance haplotype and its mutants. Plant Cell 11:1365–1376. doi:10.1105/tpc.11.7.1365

Costanzo S, Jia Y. 2010. Sequence variation at the rice blast resistance gene *Pi-km* locus: Implications for the development of allele specific markers. Plant Sci 178:523–530. doi:10.1016/j.plantsci.2010.02.014

Couch BC, Fudal I, Lebrun M-HH, Tharreau D, Valent B, Van Kim P, Nottéghem J-LL, Kohn LM. 2005. Origins of host-specific populations of the blast pathogen *Magnaporthe oryzae* in crop domestication with subsequent expansion of pandemic clones on rice and weeds of rice. Genetics 170:613–630. doi:10.1534/genetics.105.041780

De Abreu-Neto JB, Turchetto-Zolet AC, De Oliveira LFV, Bodanese Zanettini MH, Margis-Pinheiro M. 2013. Heavy metal-associated isoprenylated plant protein (HIPP): Characterization of a family of proteins exclusive to plants. FEBS J 280:1604–1616. doi:10.1111/febs.12159

de Guillen K, Ortiz-Vallejo D, Gracy J, Fournier E, Kroj T, Padilla A. 2015. structure analysis uncovers a highly diverse but structurally conserved effector family in phytopathogenic fungi. PLOS Pathog 11:e1005228. doi:10.1371/journal.ppat.1005228

De la Concepcion JC, Franceschetti M, Maclean D, Terauchi R, Kamoun S, Banfield MJ. 2019. Protein engineering expands the effector recognition profile of a rice NLR immune receptor. Elife 8:1–19. doi:10.7554/eLife.47713

De la Concepcion JC, Franceschetti M, Maqbool A, Saitoh H, Terauchi R, Kamoun S, Banfield MJ. 2018. Polymorphic residues in rice NLRs expand binding and response to effectors of the blast pathogen. Nat Plants 4:734–734. doi:10.1038/s41477-018-0248-0

De la Concepcion JC, Maidment JHR, Longya A, Xiao G, Franceschetti M, Banfield MJ. 2020. The allelic rice immune receptor Pikh confers extended resistance to strains of the blast fungus through a single polymorphism in the effector binding interface. bioRxiv. doi:10.1101/2020.09.05.284240

Dean AM, Thornton JW. 2007. Mechanistic approaches to the study of evolution: the functional synthesis. Nat Rev Genet 8:675–688. doi:10.1038/nrg2160

Delaux P-M, Hetherington AJ, Coudert Y, Delwiche C, Dunand C, Gould S, Kenrick P, Li F-W, Philippe H, Rensing SA, Rich M, Strullu-Derrien C, de Vries J. 2019. Reconstructing trait evolution in plant evo–devo studies. Curr Biol 29:R1110–R1118. doi:10.1016/j.cub.2019.09.044

Deng J, Fang L, Zhu X, Zhou B, Zhang T, Zhang D. 2019. A CC-NBS-LRR gene induces hybrid lethality in cotton. J Exp Bot 70:5145–5156. doi:10.1093/jxb/erz312

Dodds PN, Rathjen JP. 2010. Plant immunity: towards an integrated view of plant–pathogen interactions. Nat Rev Genet 11:539–548. doi:10.1038/nrg2812

Dong S, Stam R, Cano LM, Song J, Sklenar J, Yoshida K, Bozkurt TO, Oliva R, Liu Z, Tian M, Win J, Banfield MJ, Jones AME, van der Hoorn RAL, Kamoun S. 2014. Effector specialization in a lineage of the irish potato famine pathogen. Science 343:552–555. doi:10.1126/science.1246300

Eddy SR. 1998. Profile hidden Markov models. Bioinformatics 14:755–763. doi:10.1093/bioinformatics/14.9.755

Edgar RC. 2004. MUSCLE: multiple sequence alignment with high accuracy and high throughput. Nucleic Acids Res 32:1792–1797. doi:10.1093/nar/gkh340

Emsley P, Lohkamp B, Scott WG, Cowtan K. 2010. Features and development of Coot. Acta Crystallogr Sect D Biol Crystallogr 66:486–501. doi:10.1107/S0907444910007493

Evans PR, Murshudov GN. 2013. How good are my data and what is the resolution? Acta Crystallogr Sect D Biol Crystallogr 69:1204–1214. doi:10.1107/S0907444913000061

Feehan JM, Castel B, Bentham AR, Jones JD. 2020. Plant NLRs get by with a little help from their friends. Curr Opin Plant Biol 56:99–108. doi:10.1016/j.pbi.2020.04.006

Felsenstein J. 1985. Confidence Limits on Phylogenies: An Approach Using the Bootstrap. Evolution 39:783. doi:10.2307/2408678

Feuillet C, Travella S, Stein N, Albar L, Nublat A, Keller B. 2003. Map-based isolation of the leaf rust disease resistance gene *Lr10* from the hexaploid wheat (*Triticum aestivum* L.) genome. Proc Natl Acad Sci U S A 100:15253–15258. doi:10.1073/pnas.2435133100

Fujisaki K, Abe Y, Kanzaki E, Ito K, Utsushi H, Saitoh H, Białas A, Banfield MJ, Kamoun S, Terauchi R. 2017. An unconventional NOI/RIN4 domain of a rice NLR protein binds host EXO70 protein to confer fungal immunity. bioRxiv. doi:10.1101/239400

Fukuoka S, Saka N, Koga H, Ono K, Shimizu T, Ebana K, Hayashi N, Takahashi A, Hirochika H, Okuno K, Yano M. 2009. Loss of function of a proline-containing protein confers durable disease resistance in rice. Science 325:998–1001. doi:10.1126/science.1175550

Gao Y, Wang W, Zhang T, Gong Z, Zhao H, Han GZ. 2018. Out of water: The ORI in and earl diversification of plant R-genes. Plant Physiol 177:82–89. doi:10.1104/pp.18.00185

Ginestet C. 2011. ggplot2: Elegant Graphics for Data Analysis. J R Stat Soc Ser A (Statistics Soc) 174:245–246. doi:10.1111/j.1467-985X.2010.00676_9.x

González-Fuente M, Carrère S, Monachello D, Marsella BG, Cazalé AC, Zischek C, Mitra RM, Rezé N, Cottret L, Mukhtar MS, Lurin C, Noël LD, Peeters N. 2020. EffectorK, a comprehensive resource to mine for Ralstonia, Xanthomonas, and other published effector interactors in the Arabidopsis proteome. Mol Plant Pathol 21(10):1257–1270. doi:10.1111/mpp.12965

Griebel T, Maekawa T, Parker JE. 2014. NOD-like receptor cooperativity in effector-triggered immunity. Trends Immunol 35:562–570. doi:10.1016/j.it.2014.09.005

Guo L, Cesari S, de Guillen K, Chalvon V, Mammri L, Ma M, Meusnier I, Bonnot F, Padilla A, Peng Y-L, Liu J, Kroj T. 2018. Specific recognition of two MAX effectors by integrated HMA domains in plant immune receptors involves distinct binding surfaces. Proc Natl Acad Sci U S A 115:11637–11642. doi:10.1073/pnas.1810705115

Halterman DA, Wise RP. 2004. A single-amino acid substitution in the sixth leucine-rich repeat of barley MLA6 and MLA13 alleviates dependence on RAR1 for disease resistance signaling. Plant J 38:215–226. doi:10.1111/j.1365-313X.2004.02032.x

Harms MJ, Thornton JW. 2013. Evolutionary biochemistry: revealing the historical and physical causes of protein properties. Nat Rev Genet 14:559–571. doi:10.1038/nrg3540

Harris CJ, Slootweg EJ, Goverse A, Baulcombe DC. 2013. Stepwise artificial evolution of a plant disease resistance gene. Proc Natl Acad Sci U S A 110:21189–21194. doi:10.1073/pnas.1311134110

Hayashi K, Yoshida H. 2009. Refunctionalization of the ancient rice blast disease resistance gene *Pit* by the recruitment of a retrotransposon as a promoter. Plant J 57:413–425. doi:10.1111/j.1365-313X.2008.03694.x

Heidrich K, Tsuda K, Blanvillain-Baufumé S, Wirthmueller L, Bautor J, Parker JE. 2013. Arabidopsis TNL-WRKY domain receptor RRS1 contributes to temperature-conditioned RPS4 auto-immunity. Front Plant Sci 4:403. doi:10.3389/fpls.2013.00403

Hodkinson TR. 2018. Evolution and Taxonomy of the Grasses (Poaceae): A Model Family for the Study of Species-Rich Groups. Annual Plant Reviews Online. Wiley. pp. 255–294. doi:10.1002/9781119312994.apr0622

Huang L, Brooks S, Li W, Fellers J, Nelson JC, Gill B. 2009. Evolution of new disease specificity at a simple resistance locus in a crop-weed complex: Reconstitution of the *Lr21* gene in wheat. Genetics 182:595–602. doi:10.1534/genetics.108.099614

Hurni S, Brunner S, Buchmann G, Herren G, Jordan T, Krukowski P, Wicker T, Yahiaoui N, Mago R, Keller B. 2013. Rye *Pm8* and wheat *Pm3* are orthologous genes and show evolutionary conservation of resistance function against powdery mildew. Plant J 76:957–969. doi:10.1111/tpj.12345

Jacob F, Vernaldi S, Maekawa T. 2013. Evolution and Conservation of Plant NLR Functions. Front Immunol 4:297. doi:10.3389/fimmu.2013.00297

Jacquemin J, Chaparro C, Laudié M, Berger A, Gavory F, Goicoechea JL, Wing RA, Cooke R. 2011. Long-range and targeted ectopic recombination between the two homeologous chromosomes 11 and 12 in *Oryza* species. Mol Biol Evol 28:3139–3150. doi:10.1093/molbev/msr144

Jia Y, Lee FlN, Mcclung A. 2009. Determination of resistance spectra of the *Pi-ta* and *Pi-k* genes to U.S. races of Magnaporthe oryzae causing rice blast in a recombinant inbred line population. Plant Dis 93:639–644. doi:10.1094/PDIS-93-6-0639

Jones DT, Taylor WR, Thornton JM. The rapid generation of mutation data matrices from protein sequences. Bioinformatics. 1992;8(3):275–282. doi:10.1093/bioinformatics/8.3.275

Jones JDG, Vance RE, Dangl JL. 2016. Intracellular innate immune surveillance devices in plants and animals. Science 354:aaf6395–aaf6395. doi:10.1126/science.aaf6395

Kanzaki H, Yoshida K, Saitoh H, Fujisaki K, Hirabuchi A, Alaux L, Fournier E, Tharreau D, Terauchi R. 2012. Arms race co-evolution of *Magnaporthe oryzae AVR-Pik* and rice *Pik* genes driven by their physical interactions. Plant J 72:894–907. doi:10.1111/j.1365-313X.2012.05110.x

Karasov TL, Kniskern JM, Gao L, Deyoung BJ, Ding J, Dubiella U, Lastra RO, Nallu S, Roux F, Innes RW, Barrett LG, Hudson RR, Bergelson J. 2014. The long-term maintenance of a resistance polymorphism through diffuse interactions. Nature 512:436–440. doi:10.1038/nature13439

Kourelis J, Kamoun S. 2020. RefPlantNLR: a comprehensive collection of experimentally validated plant NLRs. bioRxiv 1–14. doi:10.1101/2020.07.08.193961

Kourelis J, van der Hoorn RAL. 2018. Defended to the Nines: 25 years of Resistance Gene Cloning Identifies Nine Mechanisms for R Protein Function. Plant Cell 30:tpc.00579.2017. doi:10.1105/tpc.17.00579

Kroj T, Chanclud E, Michel-Romiti C, Grand X, Morel JB. 2016. Integration of decoy domains derived from protein targets of pathogen effectors into plant immune receptors is widespread. New Phytol 210:618–626. doi:10.1111/nph.13869

Kuang H, Woo SS, Meyers BC, Nevo E, Michelmore RW. Multiple genetic processes result in heterogeneous rates of evolution within the major cluster disease resistance genes in lettuce. Plant Cell. 2004;16(11):2870–2894. doi:10.1105/tpc.104.025502

Kumar S, Stecher G, Li M, Knyaz C, Tamura K. 2018. MEGA X: Molecular evolutionary genetics analysis across computing platforms. Mol Biol Evol 35:1547–1549. doi:10.1093/molbev/msy096

Kumar S, Stecher G, Suleski M, Hedges SB. 2017. TimeTree: A resource for timelines, timetrees, and divergence times. Mol Biol Evol 34:1812–1819. doi:10.1093/molbev/msx116

Kurata N, Yamazaki Y. 2006. Oryzabase. An integrated biological and genome information database for rice. Plant Physiol 140:12–7. doi:10.1104/pp.105.063008

Langner T, Harant A, Gomez-Luciano LB, Shrestha RK, Win J, Kamoun S. 2020. Genomic rearrangements generate hypervariable mini-chromosomes in host-specific lineages of the blast fungus. bioRxiv. doi:10.1101/2020.01.10.901983

Latorre S, Reyes-Avila CS, Malmgren A, Win J, Kamoun S, Burbano HA. 2020. Differential loss of effector genes in three recently expanded pandemic clonal lineages of the rice blast fungus. BMC Biol 18:88. doi:10.1186/s12915-020-00818-z

Lee RRQ, Chae E. 2020. Variation Patterns of NLR Clusters in *Arabidopsis thaliana* Genomes. Plant Commun 1:100089. doi:10.1016/j.xplc.2020.100089

Lee S-KS, Song M-Y, Seo Y-S, Kim H-K, Ko S, Cao P-J, Suh J-P, Yi G, Roh J-H, Lee S-KS, An G, Hahn T-R, Wang G-L, Ronald P, Jeon J-S. 2009. Rice *Pi5*-mediated resistance to *Magnaporthe oryzae* requires the presence of two coiled-coil–nucleotide-binding–leucine-rich repeat genes. Genetics 181:1627–1638. doi:10.1534/genetics.108.099226

Letunic I, Bork P. 2007. Interactive Tree Of Life (iTOL): an online tool for phylogenetic tree display and annotation. Bioinformatics 23:127–128. doi:10.1093/bioinformatics/btl529

Li J, Wang Q, Li C, Bi Y, Fu X, Wang R. 2019. Novel haplotypes and networks of AVR-Pik alleles in *Magnaporthe oryzae*. BMC Plant Biol 19:1–12. doi:10.1186/s12870-019-1817-8

Li J, Zhang M, Sun J, Mao X, Wang J, Liu H, Zheng H, Li X, Zhao H, Zou D. 2020. Heavy metal stress-associated proteins in rice and arabidopsis: genome-wide identification, phylogenetics, duplication, and expression profiles analysis. Front Genet 11:1–21. doi:10.3389/fgene.2020.00477

Li L, Habring A, Wang K, Weigel D. 2020. Atypical resistance protein RPW8/HR triggers oligomerization of the NLR immune receptor RPP7 and autoimmunity. Cell Host Microbe 27:405–417.e6. doi:10.1016/j.chom.2020.01.012

Liu W, Frick M, Huel R, Nykiforuk CL, Wang X, Gaudet DA, Eudes F, Conner RL, Kuzyk A, Chen Q, Kang Z, Laroche A. 2014. The stripe rust resistance gene *Yr10* encodes an evolutionary-conserved and unique CC–NBS– LRR sequence in wheat. Mol Plant 7:1740–1755. doi:10.1093/mp/ssu112

Longya A, Chaipanya C, Franceschetti M, Maidment JHR, Banfield MJ, Jantasuriyarat C. 2019. Gene Duplication and mutation in the emergence of a novel aggressive allele of the AVR-Pik effector in the rice blast fungus. Mol Plant-Microbe Interact 32:740–749. doi:10.1094/MPMI-09-18-0245-R

Lukasik-Shreepaathy E, Slootweg E, Richter H, Goverse A, Cornelissen BJC, Takken FLW. 2012. Dual regulatory roles of the extended N terminus for activation of the tomato Mi-1.2 resistance protein. Mol Plant-Microbe Interact 25:1045–1057. doi:10.1094/MPMI-11-11-0302

MacLean D. 2019. Besthr. Zenodo. doi: 10.5281/zenodo.3374507

McNicholas S, Potterton E, Wilson KS, Noble ME. 2011. Presenting your structures: the CCP4mg molecular-graphics software. Acta Crystallogr Sect D Biol Crystallogr 40:386–394. doi: 10.1107/S0907444911007281

Maekawa T, Kracher B, Saur IML, Yoshikawa-Maekawa M, Kellner R, Pankin A, von Korff M, Schulze-Lefert P. 2019. Subfamily-specific specialization of RGH1/MLA immune receptors in wild barley. Mol Plant-Microbe Interact 32:107–119. doi:10.1094/MPMI-07-18-0186-FI

Mago R, Zhang P, Vautrin S, Šimková H, Bansal U, Luo M-C, Rouse M, Karaoglu H, Periyannan S, Kolmer J, Jin Y, Ayliffe MA, Bariana H, Park RF, McIntosh R, Doležel J, Bergès H, Spielmeyer W, Lagudah ES, Ellis JG, Dodds PN. 2015. The wheat *Sr50* gene reveals rich diversity at a cereal disease resistance locus. Nat Plants 1:15186. doi:10.1038/nplants.2015.186

Maidment JHR, Franceschetti M, Maqbool A, Saitoh H. 2020. Multiple variants of the blast fungus effector AVR-Pik bind the HMA domain of the rice protein OsHIPP19 with high affinity. bioRxiv. doi:10.1101/2020.12.01.403451

Maqbool A, Saitoh H, Franceschetti M, Stevenson C, Uemura A, Kanzaki H, Kamoun S, Terauchi R, Banfield M. 2015. Structural basis of pathogen recognition by an integrated HMA domain in a plant NLR immune receptor. Elife 4:1–24. doi:10.7554/eLife.08709

McCoy AJ, Grosse-Kunstleve RW, Adams PD, Winn MD, Storoni LC, Read RJ. 2007. Phaser crystallographic software. J Appl Crystallogr 40:658–674. doi:10.1107/S0021889807021206

Mizuno H, Katagiri S, Kanamori H, Mukai Y, Sasaki T, Matsumoto T, Wu J. 2020. Evolutionary dynamics and impacts of chromosome regions carrying R-gene clusters in rice. Sci Rep 10:1–2. doi:10.1038/s41598-020-57729- w

Murshudov GN, Skubák P, Lebedev AA, Pannu NS, Steiner RA, Nicholls RA, Winn MD, Long F, Vagin AA. 2011. REFMAC 5 for the refinement of macromolecular crystal structures. Acta Crystallogr Sect D Biol Crystallogr 67:355–367. doi:10.1107/S0907444911001314

Myszka DG. 1999. Improving biosensor analysis. J Mol Recognit. doi:10.1002/(SICI)1099-1352(199909/10)12:5<279::AID-JMR473>3.0.CO;2-3

Nei M, Gojobori T. 1986. Simple methods for estimating the numbers of synonymous and nonsynonymous nucleotide substitutions. Mol Biol Evol 3:418–426. doi:10.1093/oxfordjournals.molbev.a040410

Oikawa K, Fujisaki K, Shimizu M, Takeda T, Saitoh H, Hirabuchi A, Hiraka Y, Białas A, Langner T, Kellner R, Bozkurt T, Cesari S, Kroj T, Maidment J, Banfield M, Kamoun S, Terauchi R. 2020. The blast pathogen effector AVR-Pik binds and stabilizes rice heavy metal-associated (HMA) proteins to co-opt their function in immunity. bioRxiv. doi:10.1101/2020.12.01.406389

Okuyama Y, Kanzaki H, Abe A, Yoshida K, Tamiru M, Saitoh H, Fujibe T, Matsumura H, Shenton M, Galam DC, Undan J, Ito A, Sone T, Terauchi R. 2011. A multifaceted genomics approach allows the isolation of the rice *Pia*-blast resistance gene consisting of two adjacent NBS-LRR protein genes. Plant J 66:467–479. doi:10.1111/j.1365-313X.2011.04502.x

Petit-Houdenot Y, Langner T, Harant A, Win J, Kamoun S. 2020. A clone resource of *Magnaporthe oryzae* effectors that share sequence and structural similarities across host-specific lineages. Mol Plant-Microbe Interact 33:1032–1035. doi:10.1094/MPMI-03-20-0052-A

Potterton L, Agirre J, Ballard C, Cowtan K, Dodson E, Evans PR, Jenkins HT, Keegan R, Krissinel E, Stevenson K, Lebedev A, McNicholas SJ, Nicholls RA, Noble M, Pannu NS, Roth C, Sheldrick G, Skubak P, Turkenburg J, Uski V, von Delft F, Waterman D, Wilson K, Winn M, Wojdyr M. 2018. CCP 4 i 2: the new graphical user interface to the CCP 4 program suite. Acta Crystallogr Sect D Struct Biol 74:68–84. doi:10.1107/S2059798317016035

Prigozhin DM, Krasileva K V. 2020. Intraspecies diversity reveals a subset of highly variable plant immune receptors and predicts their binding sites. bioRxiv. doi:10.1101/2020.07.10.190785

Pupko T, Pe’er I, Shamir R, Graur D. 2000. A fast algorithm for joint reconstruction of ancestral amino acid sequences. Mol Biol Evol 17:890–896. doi:10.1093/oxfordjournals.molbev.a026369

Qi D, DeYoung BJ, Innes RW. 2012. Structure-function analysis of the coiled-coil and leucine-rich repeat domains of the RPS5 disease resistance protein. Plant Physiol 158:1819–32. doi:10.1104/pp.112.194035

Qu S, Liu G, Zhou B, Bellizzi M, Zeng L, Dai L, Han B, Wang G-L. 2006. The Broad-Spectrum Blast Resistance Gene Pi9 Encodes a Nucleotide-Binding Site–Leucine-Rich Repeat Protein and Is a Member of a Multigene Family in Rice. Genetics 172:1901–1914. doi:10.1534/genetics.105.044891

Radakovic ZS, Anjam MS, Escobar E, Chopra D, Cabrera J, Silva AC, Escobar C, Sobczak M, Grundler FMW, Siddique S. 2018. Arabidopsis HIPP27 is a host susceptibility gene for the beet cyst nematode *Heterodera schachtii*. Mol Plant Pathol 19:1917–1928. doi:10.1111/mpp.12668

Rairdan GJ, Moffett P. 2006. Distinct domains in the ARC region of the potato resistance protein Rx mediate LRR binding and inhibition of activation. Plant Cell 18:2082–2093. doi:10.1105/tpc.106.042747

Randall RN, Radford CE, Roof KA, Natarajan DK, Gaucher EA. 2016. An experimental phylogeny to benchmark ancestral sequence reconstruction. Nat Commun 7:1–6. doi:10.1038/ncomms12847

Read AC, Moscou MJ, Zimin A V., Pertea G, Meyer RS, Purugganan MD, Leach JE, Triplett LR, Salzberg SL, Bogdanove AJ. 2020. Genome assembly and characterization of a complex zfBED-NLR gene-containing disease resistance locus in Carolina Gold Select rice with Nanopore sequencing. PLoS Genet 16:1–24. doi:10.1371/journal.pgen.1008571

Saitou N, Nei M. 1987. The neighbor-joining method: a new method for reconstructing phylogenetic trees. Mol Biol Evol 4:406–425. doi:citeulike-article-id:93683

Sarris PF, Cevik V, Dagdas G, Jones JDG, Krasileva K V. 2016. Comparative analysis of plant immune receptor architectures uncovers host proteins likely targeted by pathogens. BMC Biol 14:8. doi:10.1186/s12915-016-0228-7

Saucet SB, Ma Y, Sarris PF, Furzer OJ, Sohn KH, Jones JDG. 2015. Two linked pairs of Arabidopsis TNL resistance genes independently confer recognition of bacterial effector AvrRps4. Nat Commun 6:1–12. doi:10.1038/ncomms7338

Seeholzer S, Tsuchimatsu T, Jordan T, Bieri S, Pajonk S, Yang W, Jahoor A, Shimizu KK, Keller B, Schulze-Lefert P. 2010. Diversity at the Mla powdery mildew resistance locus from cultivated barley reveals sites of positive selection. Mol Plant-Microbe Interact 23:497–509. doi:10.1094/MPMI-23-4-0497

Segretin ME, Pais M, Franceschetti M, Chaparro-Garcia A, Bos JIB, Banfield MJ, Kamoun S. 2014. Single Amino Acid Mutations in the Potato Immune Receptor R3a Expand Response to *Phytophthora* Effectors. Mol Plant-Microbe Interact 27:624–637. doi:10.1094/MPMI-02-14-0040-R

Shang J, Tao Y, Chen X, Zou Y, Lei C, Wang J, Li X, Zhao X, Zhang M, Lu Z, Xu J, Cheng Z, Wan J, Zhu L. 2009. Identification of a new rice blast resistance gene, *Pid3*, by genome-wide comparison of paired nucleotide-binding site-leucine-rich repeat genes and their pseudogene alleles between the two sequenced rice genomes. Genetics 182:1303–1311. doi:10.1534/genetics.109.102871

Shao Z-QQ, Xue J-YY, Wu P, Zhang Y-MM, Wu Y, Hang Y-YY, Wang B, Chen J-QQ. 2016. Large-scale analyses of angiosperm nucleotide-binding site-leucine-rich repeat genes reveal three anciently diverged classes with distinct evolutionary patterns. Plant Physiol 170:2095–2109. doi:10.1104/pp.15.01487

Sinapidou E, Williams K, Nott L, Bahkt S, Tör M, Crute I, Bittner-Eddy P, Beynon J. 2004. Two TIR:NB:LRR genes are required to specify resistance to *Peronospora parasitica* isolate Cala2 in Arabidopsis. Plant J 38:898–909. doi:10.1111/j.1365-313X.2004.02099.x

Stamatakis A. 2014. RAxML version 8: a tool for phylogenetic analysis and post-analysis of large phylogenies. Bioinformatics 30:1312–1313. doi:10.1093/bioinformatics/btu033

Stein JC, Yu Y, Copetti D, Zwickl DJ, Zhang L, Zhang C, Chougule K, Gao D, Iwata A, Goicoechea JL, Wei S, Wang J, Liao Y, Wang M, Jacquemin J, Becker C, Kudrna D, Zhang J, Londono CEM, Song X, Lee S, Sanchez P, Zuccolo A, Ammiraju JSS, Talag J, Danowitz A, Rivera LF, Gschwend AR, Noutsos C, Wu C, Kao S, Zeng J, Wei F, Zhao Q, Feng Q, El Baidouri M, Carpentier M-C, Lasserre E, Cooke R, Rosa Farias D da, da Maia LC, dos Santos RS, Nyberg KG, McNally KL, Mauleon R, Alexandrov N, Schmutz J, Flowers D, Fan C, Weigel D, Jena KK, Wicker T, Chen M, Han B, Henry R, Hsing YC, Kurata N, de Oliveira AC, Panaud O, Jackson SA, Machado CA, Sanderson MJ, Long M, Ware D, Wing RA. 2018. Genomes of 13 domesticated and wild rice relatives highlight genetic conservation, turnover and innovation across the genus *Oryza*. Nat Genet 50:285–296. doi:10.1038/s41588-018-0040-0

Steuernagel B, Jupe F, Witek K, Jones JDGG, Wulff BBHH. 2015. NLR-parser: Rapid annotation of plant NLR complements. Bioinformatics 31:1665–1667. doi:10.1093/bioinformatics/btv005

Steuernagel B, Periyannan SK, Hernández-Pinzón I, Witek K, Rouse MN, Yu G, Hatta A, Ayliffe M, Bariana H, Jones JDG, Lagudah ES, Wulff BBH. 2016. Rapid cloning of disease-resistance genes in plants using mutagenesis and sequence capture. Nat Biotechnol 34:652–655. doi:10.1038/nbt.3543

Tanaka S, Schweizer G, Rössel N, Fukada F, Thines M, Kahmann R. 2019. Neofunctionalization of the secreted Tin2 effector in the fungal pathogen *Ustilago maydis*. Nat Microbiol 4:251–257. doi:10.1038/s41564-018-0304-6

Tavaré S. Some Probabilistic and Statistical Problems in the Analysis of DNA Sequences. Vol 17.; 1986

Thind AK, Wicker T, Šimková H, Fossati D, Moullet O, Brabant C, Vrána J, Doležel J, Krattinger SG. 2017. Rapid cloning of genes in hexaploid wheat using cultivar-specific long-range chromosome assembly. Nat Biotechnol 35:793–796. doi:10.1038/nbt.3877

Thornton JW. 2004. Resurrecting ancient genes: Experimental analysis of extinct molecules. Nat Rev Genet 5:366– 375. doi:10.1038/nrg1324

Tran DTN, Chung E-H, Habring-Müller A, Demar M, Schwab R, Dangl JL, Weigel D, Chae E. 2017. Activation of a plant NLR complex through heteromeric association with an autoimmune risk variant of another NLR. Curr Biol 27:1148–1160. doi:10.1016/j.cub.2017.03.018

Van de Weyer AL, Monteiro F, Furzer OJ, Nishimura MT, Cevik V, Witek K, Jones JDG, Dangl JL, Weigel D, Bemm F. 2019. A species-wide inventory of NLR genes and alleles in *Arabidopsis thaliana*. Cell 178:1260–1272.e14. doi:10.1016/j.cell.2019.07.038

Varden FA, Saitoh H, Yoshino K, Franceschetti M, Kamoun S, Terauchi R, Banfield MJ. 2019. Cross-reactivity of a rice NLR immune receptor to distinct effectors from the rice blast pathogen *Magnaporthe oryzae* provides partial disease resistance. J Biol Chem 294:13006–13016. doi:10.1074/jbc.RA119.007730

Wang G-F, Ji J, EI-Kasmi F, Dangl JL, Johal G, Balint-Kurti PJ. 2015. molecular and functional analyses of a maize autoactive NB-LRR protein identify precise structural requirements for activity. PLOS Pathog 11:e1004674. doi:10.1371/journal.ppat.1004674

Wang L, Xu X, Lin F, Pan Q. 2009. Characterization of rice blast resistance genes in the *Pik* cluster and fine mapping of the *Pik-p* locus. Phytopathology 99:900–5. doi:10.1094/PHYTO-99-8-0900

Wang Z-X, Yano M, Yamanouchi U, Iwamoto M, Monna L, Hayasaka H, Katayose Y, Sasaki T. 1999. The Pib gene for rice blast resistance belongs to the nucleotide binding and leucine-rich repeat class of plant disease resistance genes. Plant J 19:55–64. doi:10.1046/j.1365-313X.1999.00498.x

Waterhouse A, Bertoni M, Bienert S, Studer G, Tauriello G, Gumienny R, Heer FT, de Beer TAP, Rempfer C, Bordoli L, Lepore R, Schwede T. 2018. SWISS-MODEL: homology modelling of protein structures and complexes. Nucleic Acids Res 46:W296–W303. doi:10.1093/nar/gky427

Weber E, Engler C, Gruetzner R, Werner S, Marillonnet S. 2011. A Modular Cloning System for Standardized Assembly of Multigene Constructs. PLoS One 6:e16765. doi:10.1371/journal.pone.0016765

Win J, Kamoun S. 2003. pCB301-p19: A Binary Plasmid Vector to Enhance Transient Expression of Transgenes by Agroinfiltration. Plant J 33:949–956.

Win J, Kamoun S, Jones AME. 2011. Purification of Effector–Target Protein Complexes via Transient Expression in *Nicotiana benthamiana*, Plant Immunity, Methods in Molecular Biology. Totowa, NJ: Humana Press. doi:10.1007/978-1-61737-998-7

Winter G. 2010. xia2 : an expert system for macromolecular crystallography data reduction. J Appl Crystallogr 43:186–190. doi:10.1107/S0021889809045701

Wu C-H, Derevnina L, Kamoun S. 2018. Receptor networks underpin plant immunity. Science 360:1300–1301. doi:10.1126/science.aat2623

Wu C-H, Krasileva K V., Banfield MJ, Terauchi R, Kamoun S. 2015. The “sensor domains” of plant NLR proteins: more than decoys? Front Plant Sci 6:5–7. doi:10.3389/fpls.2015.00134

Yamamoto E, Takashi T, Morinaka Y, Lin S, Wu J, Matsumoto T, Kitano H, Matsuoka M, Ashikari M. 2010. Gain of deleterious function causes an autoimmune response and Bateson-Dobzhansky-Muller incompatibility in rice. Mol Genet Genomics 283:305–315. doi:10.1007/s00438-010-0514-y

Yang Z. 2005. Bayes Empirical Bayes inference of amino acid sites under positive selection. Mol Biol Evol 22:1107–1118. doi:10.1093/molbev/msi097

Yang Z. 2000. Maximum likelihood estimation on large phylogenies and analysis of adaptive evolution in Human Influenza Virus A. J Mol Evol 51:423–432. doi:10.1007/s002390010105

Yang Z. 1997. PAML: a program package for phylogenetic analysis by maximum likelihood. Bioinformatics 13:555–556. doi:10.1093/bioinformatics/13.5.555

Yang Z, Nielsen R. 2000. estimating synonymous and nonsynonymous substitution rates under realistic evolutionary models. Mol Biol Evol 17:32–43. doi:10.1093/oxfordjournals.molbev.a026236

Yang Z, Nielsen R. 1998. Synonymous and nonsynonymous rate variation in nuclear genes of mammals. J Mol Evol 46:409–418. doi:10.1007/PL00006320

Yang Z, Swanson WJ, Vacquier VD. 2000. Maximum-likelihood analysis of molecular adaptation in abalone sperm lysin reveals variable selective pressures among lineages and sites. Mol Biol Evol 17:1446–1455. doi:10.1093/oxfordjournals.molbev.a026245

Yoshida K, Saunders DGOO, Mitsuoka C, Natsume S, Kosugi S, Saitoh H, Inoue Y, Chuma I, Tosa Y, Cano LM, Kamoun S, Terauchi R. 2016. Host specialization of the blast fungus *Magnaporthe oryzae* is associated with dynamic gain and loss of genes linked to transposable elements. BMC Genomics 17:370. doi:10.1186/s12864-016-2690-6

Yoshimura S, Yamanouchi U, Katayose Y, Toki S, Wang ZXZ-X, Kono I, Kurata N, Yano M, Iwata N, Sasaki T. 1998. Expression of *Xa1*, a bacterial blight-resistance gene in rice, is induced by bacterial inoculation. Proc Natl Acad Sci U S A 95:1663–1668. doi:10.1073/pnas.95.4.1663

Yuan B, Zhai C, Wang W, Zeng X, Xu X, Hu H, Lin F, Wang L, Pan Q. 2011. The Pik-p resistance to *Magnaporthe oryzae* in rice is mediated by a pair of closely linked CC-NBS-LRR genes. Theor Appl Genet 122:1017– 1028. doi:10.1007/s00122-010-1506-3

Zdrzałek R, Kamoun S, Terauchi R, Saitoh H, Banfield MJ. 2020. The rice NLR pair Pikp-1/Pikp-2 initiates cell death through receptor cooperation rather than negative regulation. PLoS One 15:1–15. doi:10.1371/journal.pone.0238616

Zess EK, Białas A, Kamoun S. 2019. Old fungus, new trick. Nat Microbiol 4:210–211. doi:10.1038/s41564-018-0351-z

Zhai C, Lin F, Dong Z, He X, Yuan B, Zeng X, Wang L, Pan Q. 2011. The isolation and characterization of *Pik*, a rice blast resistance gene which emerged after rice domestication. New Phytol 189:321–334. doi:10.1111/j.1469-8137.2010.03462.x

Zhai C, Zhang Y, Yao N, Lin F, Liu Z, Dong Z, Wang L, Pan Q. 2014. Function and interaction of the coupled genes responsible for Pik-h encoded rice blast resistance. PLoS One 9:e98067. doi:10.1371/journal.pone.0098067

Zhou B, Qu S, Liu G, Dolan M, Sakai H, Lu G, Bellizzi M, Wang G-L. 2006. The eight amino-acid differences within three leucine-rich repeats between Pi2 and Piz-t resistance proteins determine the resistance specificity to *Magnaporthe grisea*. Mol Plant-Microbe Interact 19:1216–1228. doi:10.1094/MPMI-19-1216

Zschiesche W, Barth O, Daniel K, Böhme S, Rausche J, Humbeck K. 2015. The zinc-binding nuclear protein HIPP3 acts as an upstream regulator of the salicylate-dependent plant immunity pathway and of flowering time in *Arabidopsis thaliana*. New Phytol 207:1084–1096. doi:10.1111/nph.13419

