## Supplementary material for "Two NLR immune receptors acquired high-affinity binding to a fungal effector through convergent evolution of their integrated domain": List_of_supp_files.docx

**Supplementary File 01** Full list of all Poaceae NLRs and filtering details.xlsx

**Supplementary File 02** Site selection test for K-type HMAs.xlsx

**Supplementary File 03** ancHMA prediction probabilities.xlsx

**Source File 01** Selection test for Pik-1 vs Pik-2 orthologues.xlsx

**Source File 02** Selection test for Pik-1-HMA vs NB-ARC.xlsx

**Source File 03** Raw data of Pikp-ancHMA Rmax SPR.xlsx

**Source File 04** HR scores used in SFig20.xlsx

**Source File 05** HR scores used in SFig22.xlsx

**Source File 06** HR scores for IAQVV to LVKIE mutations in Pikp-HMA.xlsx

**Source File 07** Raw data of Pikm-ancHMA Rmax SPR.xlsx
